# A computational framework to explore cellular response mechanisms from multi-omics datasets

**DOI:** 10.1101/2020.03.02.974121

**Authors:** James C. Pino, Alexander L. R. Lubbock, Leonard A. Harris, Danielle B. Gutierrez, Melissa A. Farrow, Nicole Muszynski, Tina Tsui, Jeremy L. Norris, Richard M. Caprioli, John P. Wikswo, Carlos F. Lopez

## Abstract

Recent technological advances have made it feasible to collect multi-condition transcriptome and proteome time-courses of cellular response to perturbation. The increasing size and complexity of these datasets impedes mechanism of action discovery due to challenges in data management, analysis, visualization, and interpretation. Here, we introduce MAGINE, a software framework to explore complex time-course multi-omics datasets and build mechanistic hypotheses of dynamic cellular response. MAGINE combines data management, enrichment, and network analysis and visualization within an interactive, Jupyter notebook-based environment to enable human-in-the-loop inquiry of complex datasets. We demonstrate how measurements from HL-60 cellular response to bendamustine treatment can be used to build a mechanistic, multi-resolution description of cellular commitment to fate. We present a systems-level description of signal execution from cellular DNA-damage response, to cell cycle arrest, and eventual commitment to apoptosis, mediated by over 2000 biochemical species. We further show that MAGINE can reveal unexpected, non-canonical effects of bendamustine treatment, including disruption of cellular pathways relevant to HIV infection response. MAGINE is available from https://github.com/lolab-vu/magine.

## 1 Introduction

Cellular response to perturbations can elicit molecular responses across multiple processes such as gene expression modulation, changes in protein and metabolic activity, and in extreme cases,changes in DNA structure (e.g., mutations). Modern, accessible technologies—most notably mass spectrometry (MS) and RNA-sequencing (RNAseq)—have enabled the measurement of biochemical interactions at molecular resolution for whole cellular genomes, proteomes, and metabolomes^1–3^. Recent work by our labs and others have already shown the potential of these kinds of datasets to gain a systems-level understanding of cellular response mechanisms to perturbations, with measurements that can easily number in the thousands to millions of data points^4–6^. Although these measurements in principle contain the molecular details necessary to formulate mechanistic hypotheses about cellular response to perturbations, the analysis of these datasets currently entails multiple tools, most notably enrichment- and network-based methods.

Enrichment analysis can provide insights about relevant cellular processes by comparing multiple experimental conditions following perturbations such as drug treatments^7,8^. This approach can be used to identify biological processes with altered activity by identifying groups of genes or proteins that are up- or down-regulated following treatment. Unfortunately, for the purposes of mechanistic exploration, these approaches fail to provide insights about molecular interactions that could drive a specific cellular process. For the purposes of large multi-omics and multi-experiment exploration, popular web-based enrichment analysis tools such as EnrichR^9^ and Webgestalt^10^ can only handle one sample at a time through their web interfaces, thus posing a major limitation given the high-throughput needed for multi-omics analysis.

In contrast to enrichment-based analysis methods, network-based analysis produces “maps” of biochemical species and their interactions. Networks can then be explicitly analyzed – e.g. using graph theoretic methods – to find paths or groups of relevant interactions between two or more network points. However, these become difficult to visualize and interrogate when the graph grows beyond a few tens of nodes, as seen in genome-wide networks. leading to the familiar “hairball” problem. Network tools, most notably Cytoscape^11^, partially address the needs for combined enrichment and network analysis through the use of plugins, but their capabilities for multi-sample analysis are limited. Ingenuity Pathway Analysis^12^, a useful pathway analysis tool available for multi-omics data, can provide cellular process exploration, but its closed, proprietary nature limits extension by users to meet the needs of the field.

Recognizing the need for multi-omics analysis, other tools have been built that can handle small multi-omics datasets^9,10,13–15^. However, these tools were not designed for the emerging needs posed by large and complex, multi-time point or multi-sample datasets. Thus there is an unmet need for novel tools to (*i*) integrate-omics datasets from multiple experimental modalities, (*ii*) provide a platform where multiple analysis tools can be used in tandem, and (*iii*) enable human-guided mechanism exploration within a reproducible, shareable workflow environment^16–21^

To address the needs for modern multi-omics data analysis, we developed the <u>M</u>echanism of <u>A</u>ction <u>G</u>enerator <u>I</u>nvolving <u>NE</u>twork analysis (MAGINE). MAGINE is a Python-based mechanism-of-action exploration framework that unifies enrichment and network analyses, enabling the user to explore interactions across multiple cellular processes along with the molecular interactions that drive these processes. MAGINE embraces a *literate programming* paradigm^22^ in which biological knoweldge, data exploration, and visualization are integrated within a live document within an executable web-based environment such as Jupyter notebooks, thus facilitating access to the Python scientific ecosystem.

To demonstrate the capabilities of MAGINE, we explore the time-course response of HL-60 cells to bendamustine treatment. Bendamustine is a well-established DNA-damage agent used for cancer treatments in the clinic with a consensus mechanism of action^23^. Our analysis reveals detailed, systems-level, dynamic molecular mechanisms, which comprise thousands of biochemical interactions across multiple cellular processes. These results significantly expand upon the consensus mechanism accepted for Bendamustine in cellular perturbations, which comprises a few tens of molecular interactions^23,24^. We also demonstrate how MAGINE can be used to explore bendamustine side-effects; specifically, protein interactions that could provide a mechanistic explanation for previous reports of bendamustine treatments in HIV patients. Finally, all our MAGINE-based analysis is documented using Jupyter notebooks, which offer a means to transparently report complete analyses, suitable for distribution across members of the scientific community to evaluate and expand on as desired. In summary, MAGINE unifies multiple practices in the field onto a common, open, scalable platform that enables the extraction of cellular response mechanisms from large, complex-omics datasets.

## 2 Results

### MAGINE: A framework to explore cellular response mechanisms using multi-omics

MAGINE is implemented in Python, a suitable language for a biological exploration platform due to its ease of use, large user base, and integration with over 200,000 packages in the Python Package Index as of the writing of this manuscript. The platform has been tested on Windows, Mac OS, and Linux. It can be used within the Python console, interactive Jupyter Notebooks, or within data processing and analysis pipelines. We envisage the majority of users are best served through the Jupyter notebook option, and thus describe that in the most detail, including a tutorial notebook (Supplementary Information).

MAGINE comprises three main modules (Figure 1): data management and visualization, enrichment analysis, and network analysis. Each module can be used independently, but data can be easily shared across modules. We present a brief summary of each module below, followed by an applied case study.

**Figure 1:**
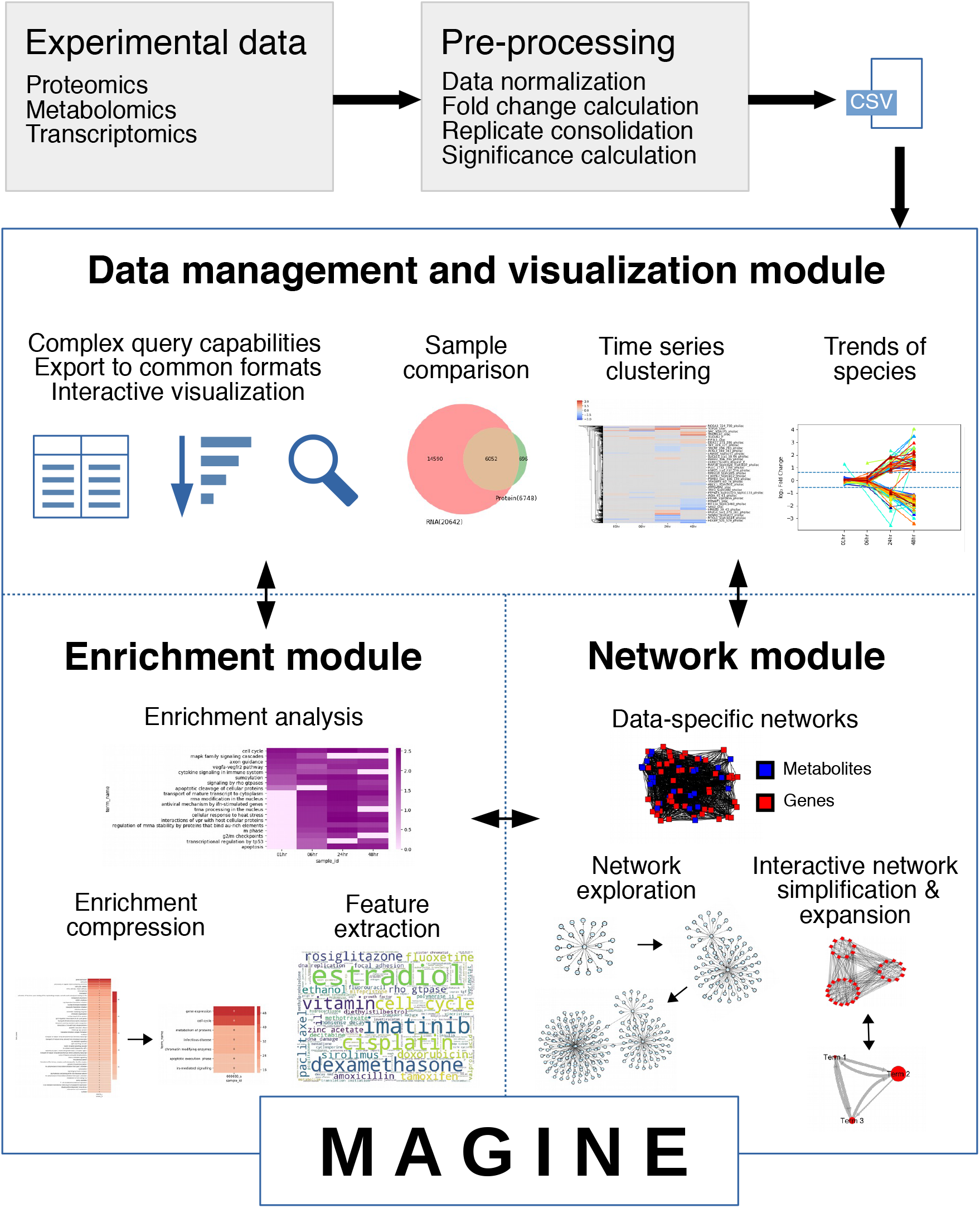
The MAGINE platform. MAGINE is designed for quantitative time-series multi-omic data. It is built around three concepts: data management, enrichment analysis, and network exploration. The modular design allows flow of information among the data, enrichment, and network modules, allowing an iterative cycle with varying levels of resolution. Example outputs of each module are provided in each panel.

### Data analysis module

The data management module handles data storage, access, and analysis. MAGINE utilizes the *pandas*^25^ library to provide database-like capabilities for-omics data querying. Data are loaded using a tabular, comma-separated values (CSV) file, with one measurement per row (see *Methods*). MAGINE stores these in an *ExperimentalData* class, which provides a simple, high-level interface to access, filter, and search these data as needed. This module also provides visualization capabilities, which include sample comparisons, time-series clustering, and species differential expression trends over time (Figure 1). A summary of these methods is shown in Supplementary Table **S2**, and example usage can be found online or in Supplementary Notebook S1.

### Enrichment module

The enrichment analysis module identifies over- or under-represented biochemical species that share common annotation (e.g., biological processes or molecular functions) compared to random background sets^26^. MAGINE leverages the capabilities of *EnrichR*, which includes over 120 gene set “libraries”^9,27^. For analysis, users provide one or more lists of genes, which can be manually constructed or created by the *ExperimentalData* class (e.g., all up-regulated genes, species detected on a specific experimental platform, or filtered by time point). MAGINE automates analysis through EnrichR, as shown in Supplementary Figure **S2**. A single command is provided to query EnrichR across all time points and data platforms present in a dataset. The results are stored in an *EnrichmentResult* class, which includes methods to further query, filter, and visualize enrichment terms. Terms can be compared, ranked, grouped, and visualized with built-in methods (e.g., time-series heat map, word cloud). Genes corresponding to each term can be extracted and used to subset ExperimentalData or create subgraphs.

Even through the use of multiple databases, traditional enrichment analyses can yield terms of varied granularity, ranging from very broad (e.g., “biological process”), to highly specific (e.g., “cysteine-type endopeptidase inhibitor activity involved in apoptotic process”), within a single results output. The problem is exacerbated due to each gene mapping to multiple terms, thus introducing term redundancy, increasing the total number of explorable terms and hindering human interpretation. To address these issues, we developed an ontology compression method to aggregate terms based on gene content similarity (described in *Methods*). This significantly reduces enrichment term redundancy, which in turn greatly aids human interpretation. For example, on the bendamustine dataset analyzed herein, 84 terms from traditional enrichment analysis of our data can be compressed to 17 terms, an 80% reduction (see *Results*). A summary of functions available in the enrichment module is shown in Supplementary Table **S3**, and example usage can be found online or in Supplementary Notebook S3).

### Network module

MAGINE’s network module allows users to build, query, and visualize molecular and gene annotation networks. A summary of the network module’s methods is shown in Supplementary Table **S4** and example usage can be found online or in Supplementary Notebook S2. The network module utilizes connectivity information from multiple databases, including KEGG^28,29^, SIGNOR^30^, Reactome Functional Interactions^31^, BioGrid^32,33^, and HMDB^34,35^. The graphs underlying these database are merged into a single network, which can be used to perform queries (e.g., find paths between nodes, apply clustering methods) or construct context-specific networks based on a user-provided seed species list. Seed nodes can be obtained from various sources: significantly changed species, mutational evidence, literature review, or manual curation. The module iterates through the databases and expands the network by adding edges and nodes based on connectivity to the seed species in those databases (Supplementary Figure **S4**). The resulting networks are a subset of the background network, focused around the specific molecular seed species. These networks can be large (>20 000 nodes and >100 000 edges). MAGINE users can use the *Subgraph* class to generate subnetworks (Supplementary Table **S4**). For example, these functions enable users to find paths between species, or find neighbors (upstream or downstream) of nodes of interest. Options are provided to limit the network expansion, such as setting a maximum distance from specific nodes.

Additionally, we introduce a method to create a coarse-grained network from gene sets, which we refer to as an Annotated Gene-set Network (AGN). The AGN is motivated by the desire to combine dynamic, high-level information about biological processes from enrichment analysis with inter-process communication provided by molecular networks. This results in a coarsegrained network, where nodes are biological process terms and the edges are connections between the sets of nodes in the molecular network. This can be expanded into a fine-grained network, which contains the chemical species and their connections, thus enabling multi-resolution exploration. MAGINE’s network module also provides various tools for network visualization, allowing users to overlay data or update network attributes. Users can visualize networks in Jupyter notebooks using *cytoscape.js*^36^, modify a *cytoscape* session via *py2cytoscape*^37^, or create a sequence of figures of network activity through *matplotlib*^38^, *igraph*^39^, or *graphviz*^40^. Networks can be exported using *networkx*^41^, for further external manipulation.

### Case study: Using MAGINE to explore the temporal response of bendamustine-treated leukemia cells

To demonstrate the capabilities of MAGINE, we extracted the mechanism of action from a multi-omics dataset containing molecular measurements for the response of HL-60 cells^42^ to treatment with bendamustine, a nitrogen mustard alkylating compound used for the treatment of chronic lymphocytic leukemia and non-Hodgkin’s lymphoma^24^. Bendamustine is a cytotoxic agent that can induce DNA interstrand cross links (ICLs), which inhibits replication and transcription, leading to cell cycle arrest, and ultimately, cell death^23,24^.

The main goal of this study was to probe and expand the consensus understanding of molecular species and biological processes involved in cellular response to bendamustine treatment. Data were gathered across three proteomics platforms (label-free, SILAC, phospho-SILAC/phSILAC) and RNAseq, spanning between 30 seconds and 72 hours post-drug exposure with matching untreated controls (see Methods; Supplementary Figure **S1**). The analysis is summarized and documented in Supplementary Notebook S1.

In total, we detected 54 818 unique molecular species across all platforms, of which 14426 were significantly changed versus control in at least one time point upon drug treatment. By platform, we saw 5 115 changed species versus control from phSILAC MS, 1 653 from label-free MS, 133 from SILAC MS, and 736 from RNAseq (Supplementary Figure **S1**b). Early changes were detected at the phosphorylation level; phSILAC detected >700 such changes at the 30s time point. Phosphorylation events occurred relatively evenly across all sample time points, while a gradual increase of significant differences versus control occurred over time in overall protein and RNA (Supplementary File S1). We next examined the overlap between significantly changed species across the experimental platforms (Supplementary Figure **S1**c and d). The highest overlap among proteomic platforms was between phSILAC and label-free (643 species). Only 88 species were significantly changed versus respective controls in both RNA and at least one of the proteomic platforms (phSILAC, SILAC, and label-free; Supplementary Figure **S1**). Indeed, even within the protein platforms, a majority (>2 500) of measured species were unique to a single platform, while 688 species were measured in at least two platforms, and only 22 species were measured in all three platforms. Within-platform detection is generally reliable and repeatable^4^, thus the low overlap between significant species changes across platforms demonstrates the value of integrative, multiplatform analysis.

We proceeded to further analyze the dataset with a custom MAGINE workflow, integrating both network and enrichment analyses (Figure 2). First, we constructed a master signaling network utilizing significantly changed species across all platforms and time points as seed species. Separately, we performed an enrichment analysis across 52 reference gene sets (provided by EnrichR) for each time point and experimental platform; MAGINE automates this entire process using a single instruction, `run_enrichment()`. Each enrichment result set was arranged by three change types: up-regulated, down-regulated, and significantly changed (both up- and down-regulated) species; and across 12 time points and four experimental platforms, resulting in 78 total result sets (not all combinations are represented). In total, there were 20 758 enriched terms across the 52 reference gene sets with an adjusted p-value ≤0.05.

**Figure 2:**
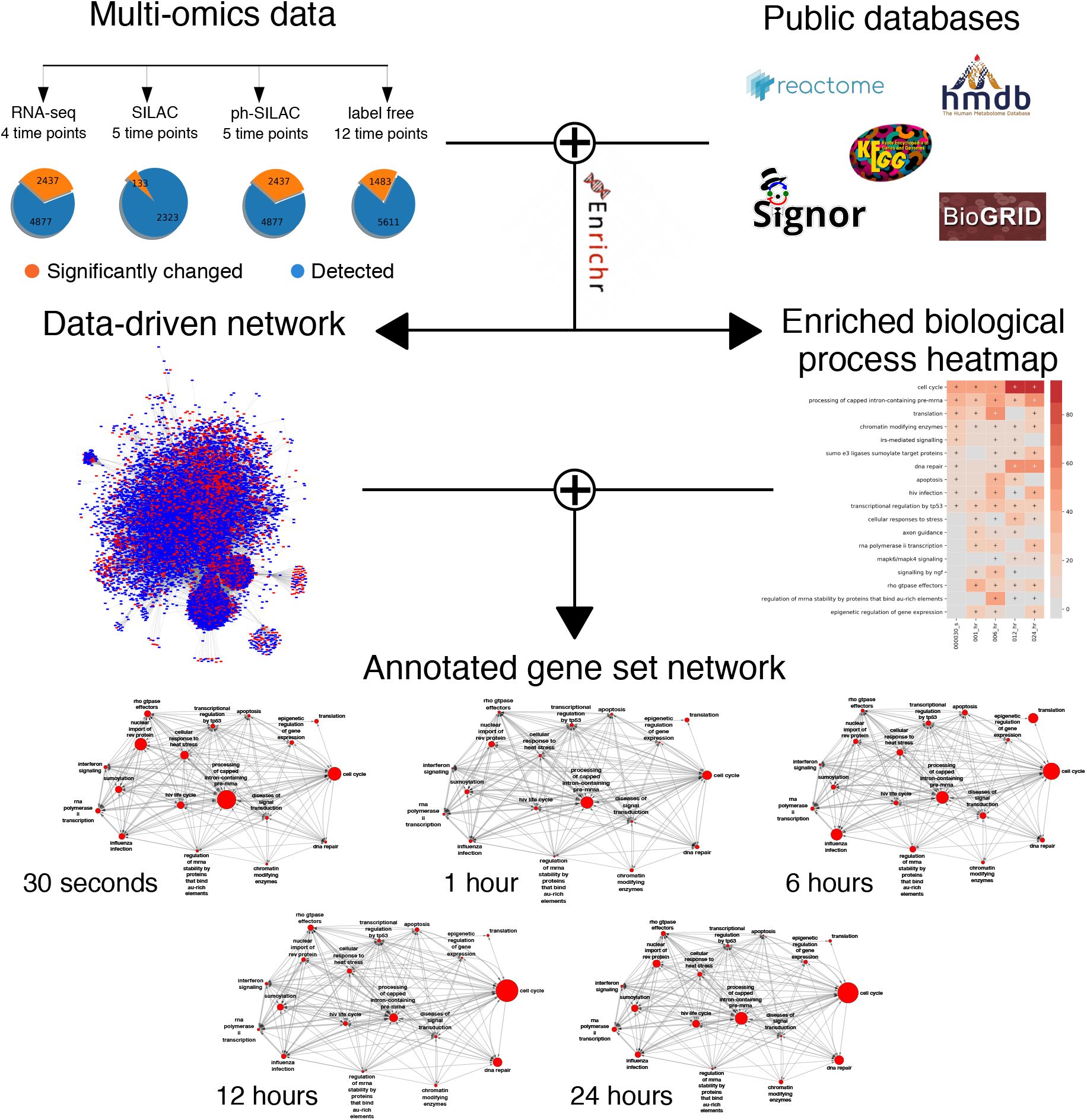
Example workflow of bendamustine mechanism extraction. Multi-omics datasets (*top left*) are combined with public databases (*top right*) to generate detailed interaction networks (*middle left*) and enriched annotated gene set (AGS) heatmaps (*middle right*). The data-derived network is then pruned by extracting a subnetwork of genes from high-ranking AGS terms and collapsed to produce annotation-level networks (*bottom*), where each node is labeled by an AGS term and scaled according to enrichment of that term; the width of each edge represents the number of edges between the terms. The network can be viewed at each time point or as an animation (Supplementary File S5).

We initially focused on phospho-protein changes to elucidate signaling activity. Using the MAGINE filtering capability, we selected only enriched terms originating from phSILAC data with the *Reactome_2016* reference gene set, resulting in 561 unique terms. Next, we required terms to be significantly enriched in at least 3 of 5 time points, resulting in 84 terms. To reduce redundancy of closely-related terms (e.g., “cell cycle”, “cell cycle, mitotic”), we applied the *remove_redundancy()* method, resulting in 17 terms (Supplementary Figure **S3**); a manageable number for visualization.

We created a heatmap to examine the dynamics of these 17 terms over time (Figure 2, center right), which include *cell cycle, processing of capped intron-containing pre-mRNA, DNA repair, apoptosis*, and *HIV infection*. As shown in Supplementary Figure **S3**c, we were able to expand resulting terms such as *cell cycle* to view aggregated and redundant terms removed in the compression process described above, such as *nuclear envelope breakdown, DNA replication*, and *G2/M checkpoints*, all of which are known bendamustine responses^23^.

An enrichment analysis by itself provides no detail on how terms affect one another. Therefore, we constructed an annotated gene set network (AGN) for each time point (Figure 2, bottom). Each node in the AGN represents an enriched biological process, and the edges between the nodes reflect the directed connectivity between species that are annotated with that node’s biological process term. As shown in Figure 2 and Supplementary File S5, this network can be visualized over time, where the node size will be adjusted based on the enriched value of the term at each time point, providing a high-level representation of the dynamics of signal flow, with animation if desired. These figures give insights into the high-level organization of signaling dynamics. At 30 seconds, the ontology term with the most changes involves processing of pre-mRNA, suggesting alterations in transcriptional activity. By 6 hours, terms involving genes associated with apoptosis, protein translation, and HIV response (explored in the next section) have their relative peaks at 6 hours. However, by the 12 and 24 hour time points, cell cycle becomes the dominant ontology term. This suggests early response is mediated through phosphorylation and changes in transcription, while later activity manifests through signaling activity. Looking at network connectivity, *RNA polymerase II transcription* is regulated by four terms (*cellular response to stress, cell cycle, sumo E3 ligases sumoylate target proteins, HIV infection*), but only regulates a single term (*rho GTPase effectors*), suggesting the transcription is highly regulated in response to bendamustine. The *cell cycle* node is connected to all other nodes in the AGN, in agreement with its importance in regulating response to bendamustine^23^. Overall, the AGN provides a broad representation of the data that allows users to visualize major cellular processes and their mutual interactions. Users can then identify terms of importance to be used as a starting point for further exploration.

### MAGINE elucidates DNA repair, cell cycle, and cell death proteins central to bendamustine response

We next examined how our data recapitulate existing knowledge of bendamustine’s mechanism of action (the “canonical mechanism”). We extracted a molecular network from the AGN focused on the terms most commonly associated with bendamustine response^23^: *DNA repair, cell cycle*, and *apoptosis* (Figure 3, top center). There are a total of 65, 146, and 28 species in the network for the terms *DNA repair, cell cycle*, and *apoptosis* respectively. 33 species were classified as both *DNA repair* and *cell cycle*, 37 genes were classified as *cell cycle* and *apoptosis*, two genes were in all three terms, while no genes were classified as both *DNA repair* and *apoptosis*. We saw higher connectivity between *DNA repair* and *cell cycle* (132 outgoing and 117 incoming edges) and *cell cycle* and *apoptosis* (154 outgoing and 609 incoming edges) compared to *DNA repair* and *apoptosis* (24 outgoing and 56 incoming edges).

**Figure 3:**
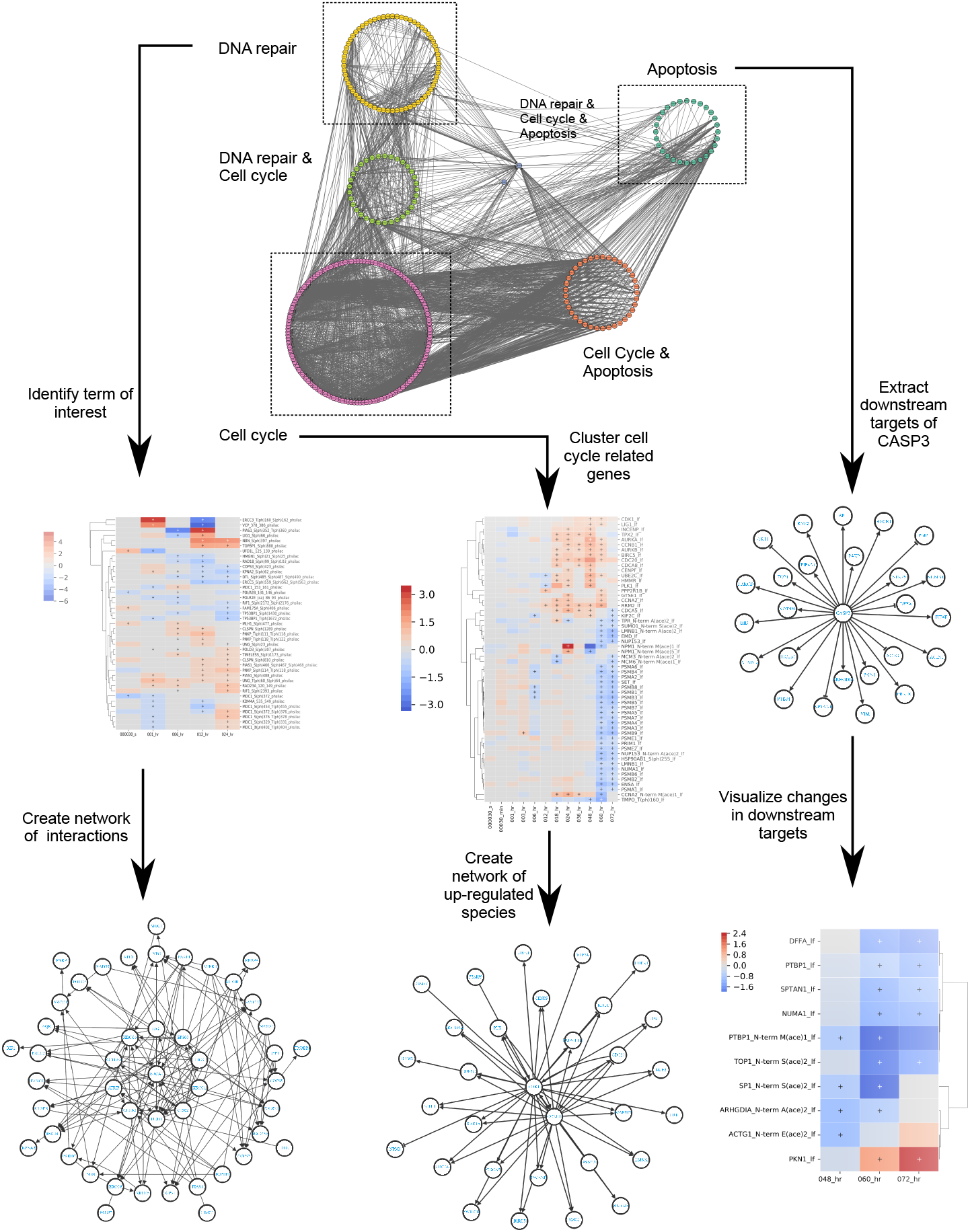
Custom workflows for exploring known biological processes of bendamustine. Molecular network constructed from DNA repair, cell cycle, and apoptosis classified species, the canonical processes involved in bendamustine response (*top*). The network is an expanded version of the annotated gene network from Figure 2. From the term “DNA repair”, MAGINE can visualize (*center left*) as well as construct a signaling network corresponding to the term’s molecular species (*bottom left*). Due to CASP3 importance in executing apoptosis, we constructed a network of downstream species (*center right*). We then filtered the experimental data to generate a heatmap (*bottom right*). Eight of nine species are down-regulated, indicating CASP3 activity. We then visualized the changes to cell cycle-related genes over time (*center*), and subsequently constructed a molecular network based on CCNB1 and CDK1 (*middle bottom*), two genes responsible for G2-M transition of cell cycle.

We next examined each of these ontology terms at the molecular level using MAGINE’s network module. In the *DNA repair* term, we saw 173 phosphorylation events across 80 proteins, 43 changes in protein expression, and 5 changes of RNA expression (Figure 3, left). Of these, we identified proteins involved in DNA ligase (LIG1, LIG2), required to alleviate interstrand crosslinks, and double stranded break response MRN complex (RAD50, MRE11A, NBN)^43^, suggesting that the interstrand cross-links are corrected during DNA replication creation of double stranded breaks. We also identified proteins involved with base excision repair pathways (DDB1, XRCC1, XRCC2, XRCC5, XCCC6) and nucleotide excision repair (ERCC3, ERCC5, ERRC6)^44^, suggesting that the pyrimidine analog properties of bendamustine also contribute to DNA damage in addition to interstrand cross-links.

We next extracted and visualized changes in cell cycle status (Figure 3, center). For the “cell cycle” term, we saw 444 significant phosphorylation events across 153 proteins, 111 changes in protein expression, and 17 changes of RNA expression. We clustered protein expression changes and created a sub-network of up-regulated species. We saw up-regulation of CDK1, AURKA, AURKB, CCNB1, CDC20, and PLK1^45–47^. Of these, CDC20, CDK1, and CCNB1 regulate chromosome separation^48^, while expression of AURKA/AURKB/PLK1 regulates progress through the G2-M checkpoint^46^. Without this checkpoint, activated apoptotic signals could lead to mitotic catastrophe during metaphase/anaphase transition, a mechanism known to occur in response to bendamustine^23^.

Finally, we used the *Subgraph* module to explore downstream targets of CASP3, an effector caspase whose cleavage marks commitment to apoptosis^49^. Although significant changes in CASP3 concentration were not detected from the experimental datasets, enough biological evidence about CASP3 activation exists that motivate exploration of its downstream targets. To accomplish this, We created a network of downstream targets of CASP3 and visualized their expression levels over time (Figure 3, right). We saw 9 proteins that were down-regulated and one up-regulated at late time points, after apoptosis was induced. Thus, despite not measuring changes in CASP3, MAGINE allowed us to gather evidence suggesting its activation in a signaling capacity.

### Bendamustine regulation of nuclear pore proteins and regulators of nuclear trafficking: potential clinical significance to HIV infection

Throughout the exploration of the HL-60 response to bendamustine dataset, we noticed that the term *HIV infection* was a significantly enriched ontology term that apparently played no role in the cell-death response associated with bendamustine. However, since HIV infection hijacks and modulates cell cycle and DNA repair^50,51^, we examined whether a plausible explanation of the enrichment in proteins associated with HIV infection is the overlap of underlying species with these expected biological process terms. The data showed that this overlap did not completely account for the finding (Figure 4). Proteins down-regulated in response to bendamustine marked with the *HIV infection* term include those related to the nuclear pore (NUP107, NUP133, NUP153, NUP155, NUP160, NUP188, NUP210, NUP214, NUP35, NUP37, NUP50, NUP62, NUP88, NUP93, NUP98) as well as regulators of the trafficking through the nuclear pore (RANBP1, RANBP2). Perturbations to RANBP2 are known to disrupt the HIV virus’ ability to shuttle the HIV-1 Rev protein between the cytoplasm and nucleus, making it a potential target for inhibition of HIV^52^. Additionally, CCNT1, a co-factor of the HIV-1 Tat protein necessary for full activation of viral transcription, is down-regulated at the 60 and 72 hour time points. Thus, the ability of bendamustine to perturb HIV infection-related genes could have clinical relevance. While treatment with bendamustine (for other disease indications) in HIV-positive patients is rare, case reports state that two HIV-positive patients with chronic lymphocytic leukemia received bendamustine without adverse outcomes^53^. Therefore, unbiased analysis of this dataset enabled side effects exploration that could lead to further leads in HIV treatment. Although such results clearly require further validation, they demonstrates the ability of MAGINE to identify non-canonical cellular pathways effected by drug treatment.

**Figure 4:**
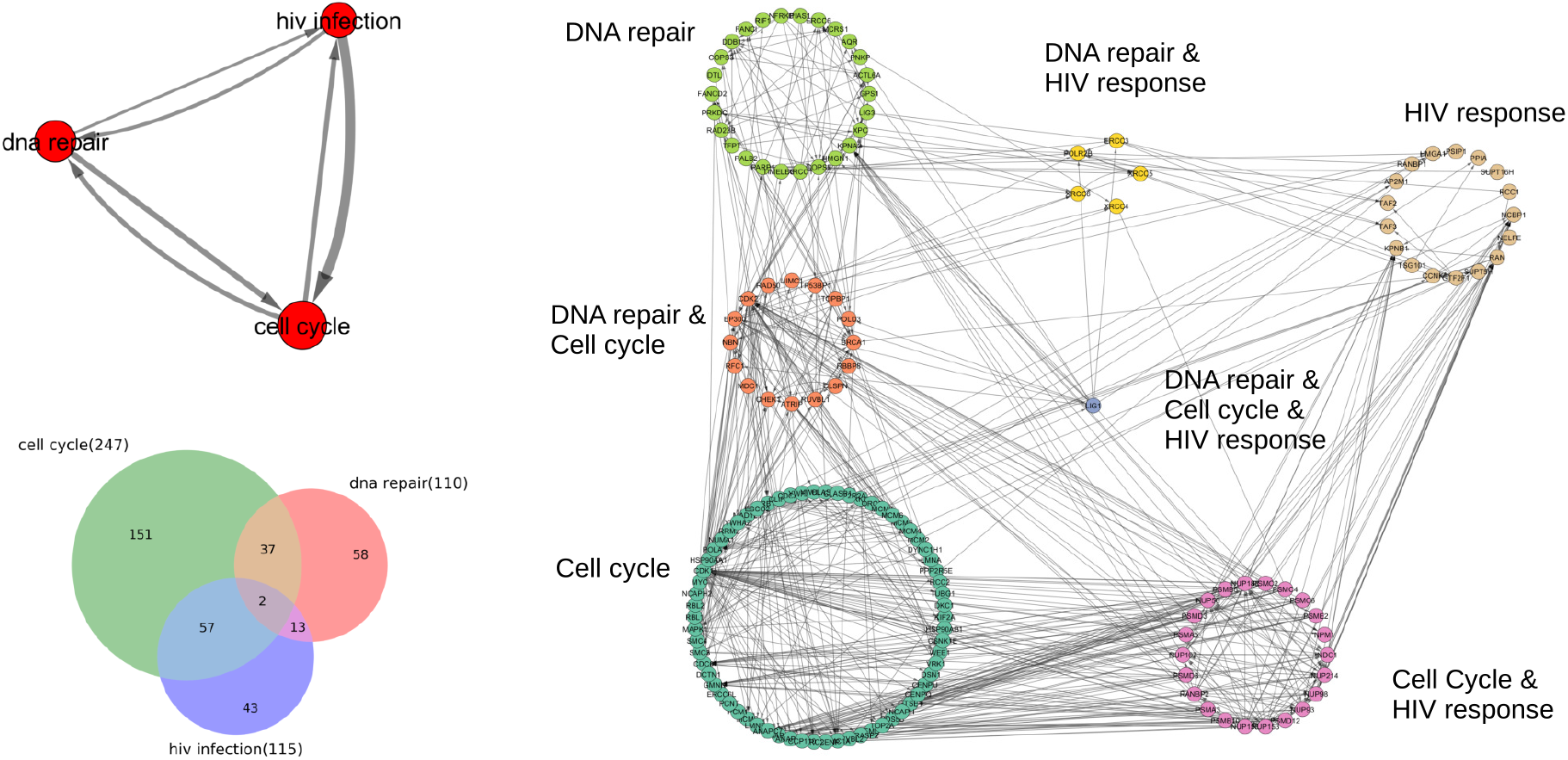
Common genes between bendamustine mechanism processes and the GO term *HIV infection*. An unexpected outcome of the analysis revealed that proteins related to the GO term *HIV infection* were modulated with bendamustine. An annotated gene set network of the interaction between the GO terms *DNA repair, HIV infection*, and *cell cycle* (*top left*). The overlap between genes labeled in each pathway labeled by Reactome (*bottom left*). Molecular interaction network of all significantly changed species grouped according to their classified pathway (*right*).

## 3 DISCUSSION

Experimental advances in-omics data generation have provided a wealth of data and opportunities to advance currently available tools. The most common analysis types include network and enrichment analysis, which were originally designed for single sample, single platform studies. These analyses are complementary: network analyses give molecular interaction insights, while enrichment analyses show the coordination of molecular activity at the level of biological processes. Although both approaches have been widely used, computational tools allowing users to employ both analyses in a complementary fashion on multi-sample, multi-omics data have been lacking. Here we demonstrated that MAGINE can integrate multi-sample-omic data with enrichment analysis and data-specific signaling networks in a single computational environment, which allows users to gain many types of biological insights and to explore their data interactively and efficiently.

MAGINE enables users to construct custom analysis protocols in Jupyter notebooks that are both reproducible and transferable. Existing software such as Biojupies^54^ and PaDua^55^ have demonstrated the power of this approach on RNAseq and phospho-proteomics analysis. We expect the use of Jupyter notebooks to increase and see MAGINE as highly complementary to such pipelines.

Multi-omics mesaurements provide larger cellular coverage than traditional *in vitro*/in cell measurements (Western blots or ELISA) and identification of unexpected significantly changed biochemical species is common. MAGINE enables users to elucidate these findings through exploration of the molecular data and their biological network context, which is often a slow step. Here, we demonstrated MAGINE’s ability to quickly explore why the term *HIV infection* was detected and its relationship to known bendamustine responses. We expect this utility to be useful in identifying and exploring non-canonical cellular pathways that are often disregarded or ignored.

Our results show the complexity in interpreting multi-omics data, as the response to bendamustine involves multiple genes and cellular pathways. Single time point measurements do not capture an entire mechanism, with some events occurring early (phosphorylation events of DNA repair proteins), and some later (up-regulation of cell cycle proteins, cleavage events of apoptotic CASP3). Interactive exploration of these data enables users to piece together the mechanism and design further experiments for validation and follow-up exploration.

In summary, MAGINE enables users to easily switch between enrichment analysis, networks, and experimental data within an iterative, interactive framework. This allows users to vary resolutions (molecular/biological process, static/dynamic) depending on the question at hand, and generate new hypotheses. As improvements and falling costs with-omic data generation allow multiple sample acquisitions per perturbation, we envision the need for integrative tools such as MAGINE increasing significantly.

## METHODS

### MAGINE software implementation

MAGINE is implemented as a Python package. It leverages several existing Python packages, including *pandas*^25^, *NumPy*^56^, *Bioservices*^57^, *Matplotlib*^38^, *Seaborn*^58^, *matplotlib-venn*, and *networkx*^41^ packages. Optional but recommended dependencies include *jupyter-notebook*^59^, *python-igraph*^39^, *py2cytoscape*^37^, and *plotly*^60^ Python packages and *graphviz*^40^ and *Cytoscape*^11^ software for additional visualization features.

### Data format

MAGINE is designed for quantitative datasets. These data are loaded from a single file in a tabular format, with columns identifying the sample, time point, and experimental platform. Due to the large number of possible experimental platforms, we generalized naming of columns to support a wide variety of data types. MAGINE requires the following column labels: *identifier, label, species_type, significant, fold_change, p_value, source*, and *sample_id*, with a sample table shown in Supplementary Table **S1**. This format uses a *stacked* data file notation (*denormalized*, in database parlance), which provides ease of editing and compatibility (a single, text format table is all that is needed) at the expense of data redundancy. For *identifier*, we use HGNC^61^ and Human Metabolite Database (HMDB)^35^IDs for genes and compounds, respectively (we also provide tools to map other identifiers to these formats). The *label* column can be used to store any other information, such as post-translational modifications, aliases, or molecular weights. The *source* and *sample_id* are used to label the experimental platform and sample condition (i.e., time point). The *species_type* column is used to state if the species is a metabolite or gene (we use “gene” to label all possible gene products and allow the *source* column to specify the type). The *fold_change* column and *p_value* column are for the fold-change and statistical significance of the data compared to control (supplied from the raw data processing step performed before data are loaded into MAGINE). Note that the *p_value* should be the adjusted value if using any form of multiple hypothesis testing. Finally, the *significant* column is used to identify if a given measurement is significant compared to control. The definition of significant is deliberately left to the user: in the situation where each experimental platform is handled by a different team, those teams can apply their own standards or software to defining significance, if desired.

Once loaded, data are stored in an instance of MAGINE’s *ExperimentalData* class, built on the Pandas DataFrame with added methods designed for biological data. Our goal was to simplify and provide shorthand properties/methods to minimize the code needed to perform common operations such as filtering, querying, aggregating, and visualization of the data. A summary of these methods is shown in Supplementary Table **S2** and example usage can be found online or in Supplementary Notebook S1). MAGINE allows for custom workflows to be assembled from its component modules and methods (Figure 5). Python *properties* are constructed around the *significant, source* and *sample_id* columns, allowing users to gather only significant changes (*data.sig*), extract only label-free proteomics data (*data.label_free*), extract out only sample id rows labeled “wild type” (*data.wild_type*), or filter up-regulated fold-changed species (*data.up*). Importantly, filters can be chained together to create complex queries, with less complex syntax, and methods directly applied to the resulting information (*data.label_free.sig.up.heatmap()*). Identifiers or labels of these queries can then be aggregated (*data.id_list* or *data.label_list*) and passed to enrichment analysis or to construct a network. Additionally, the class has direct access to common biological plotting functions such as volcano plots (*data.volcano()*), heatmaps (*data.heatmap()*), or individual species plots *(data.plotspecies(…)*).

**Figure 5:**
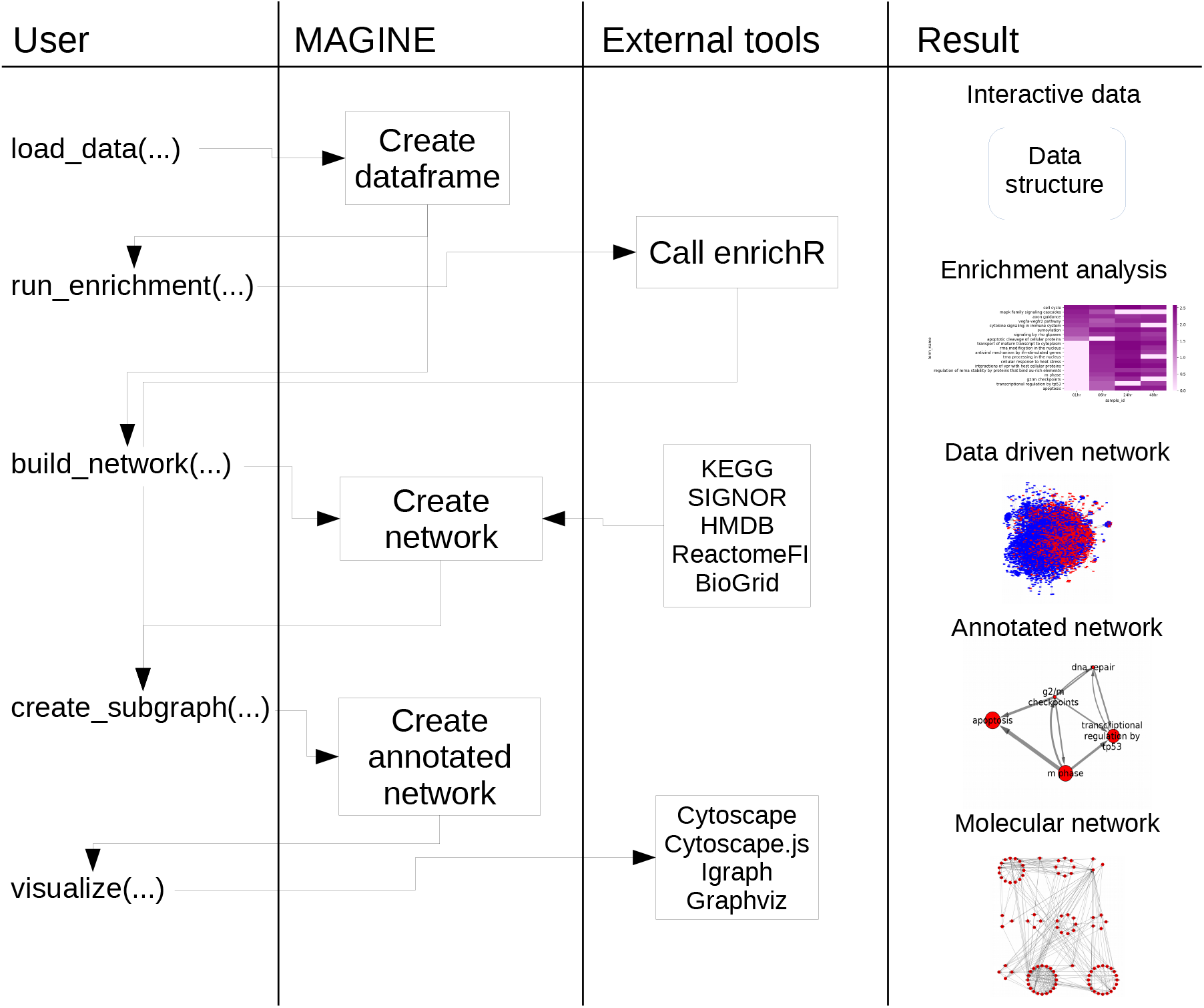
Example of MAGINE workflow. The figure shows example function calls made by the user (*left column*) and how these are handled by MAGINE and external tools (*middle columns*), and examples of resulting output (*right column*). This is a typical processing workflow where users desire to run enrichment analysis, construct a data-specific network, and construct a compressed annotated gene set network.

### Enrichment term aggregation

MAGINE contains a method, *remove_redundancy*, for reducing the number of gene enrichment terms by aggregating redundant terms. First, it calculates the ratio of the sizes of the intersection and union (Jaccard index^62^) between genes within all term pairs. It then ranks all terms based on either their combined score (from EnrichR), number of genes in the term, or p-value. Starting from the highest ranked, it compares all lower ranked terms and removes them if their similarity is above a user-defined threshold, as shown in Supplementary Text 1 and demonstrated in Figure **S3**. This allows the user to minimize the number of total terms while maintaining a level of information content that preserves total information.

### Annotated gene set network construction

Annotated gene set networks start with a set of annotation terms, which can be selected based on expert knowledge, rank of enrichment, all compressed terms, or any other criteria. We first extract the set of nodes from a molecular network identified with the selected annotation terms. From there, we search through all possible combinations of pairs between the terms. For example, if term 1 has genes (A, B, C) and term 2 has (D, E), we count the number of edges from the possible sets ((A, D), (A, E), (B, D), (B, E), (C, D), (C, E)) that are found in the network edges. We then do the reverse (term 2 to term 1). If a node is in both sets, we consider the edges that connect the other term, not edges that are within the term. This is demonstrated in Supplementary Figure **S5**.

### Software availability and requirements

MAGINE is released as open source software under the GNU General Public License, version 3. Source code for *MAGINE* can be found at github.com/LoLab-VU/Magine. Full documentation can be found at magine.readthedocs.io. MAGINE has a full software test suite, which is executed using continuous integration and checked for code coverage. The project is version-controlled on GitHub. We encourage the community to post issues, questions, enhancement requests, and code contributions.

### Experimental methods

Briefly, HL-60 cells were plated and exposed to 100 M bendamustine in triplicate for each time point, as described previously^4^. Sample preparation for transcriptomics, label-free proteomics, and metabolomics analyses was performed as previously described^4^. SILAC and phSILAC samples were prepared in a similar manner as reported^5^, with the exception that SILAC samples were run by 1D chromatography rather than MudPIT. After introduction of drug, measurements were collected at 12 time points ranging from 30 seconds to 72 hours across 6 experimental platforms as shown in Figure **S1**a. Each dataset was then formatted and combined into a single CSV to be used with MAGINE.

## Supporting information

S1-Jupyter Notebook [Data Exploration]

S2-Jupyter Notebook [Network Creation and Exploration]

S3-Jupyter Notebook [Enrichment Analysis]

S4-Jupyter Notebook [Annotated Gene Set Networks]

Animated Annotated Gene-Set Network Dynamics

## 5 ACKNOWLEDGEMENTS

We thank Randi Gant-Branum, Lauren D. Palmer, Stacy D. Sherrod, D. Borden Lacy, Eric P. Skaar, and John A. McLean for data generation and feedback. We also thank Oscar Ortega, Lisa Poole, and Nate Braman for input and ideas, and Michael Ripperger for discussions regarding visualization tools.

Research was sponsored by the U.S. Army Research Office and the Defense Advanced Research Projects Agency and was accomplished under Cooperative Agreement no. W911 NF-14-2-0022. The views and conclusions contained in this document are those of the authors and should not be interpreted as representing the official policies, either expressed or implied, of the Army Research Office, DARPA, or the U.S. Government. The U.S. Government is authorized to reproduce and distribute reprints for Government purposes notwithstanding any copyright notation herein. This work was supported in part by the National Science Foundation grant MCB-1411482 (C.F.L., J.C.P). This work was also supported in part by the National Institutes of Health cooperative agreement 1U01CA215845-01 (C.F.L.).

