## Supplementary methods, tables, and figures for "A computational framework to explore cellular response mechanisms from multi-omics datasets"

#### 1 Supplementary Information

##### 1.1 Notes

All scripts, Jupyter notebooks, and data are available as part of the supplement, and are also available online at https://github.com/LoLab-VU/MAGINE_Supplement_notebooks. Scripts were created using Python 3.7 and tested on Windows 10 x64. MAGINE can be installed with the command: `pip install magine`.

##### 1.2 Jupyter Notebooks

- **Supplementary File S1**: `supplement_notebook_1_data_exploration.pdf`
  contains examples in data exploration.
- **Supplementary File S2**: `supplement_notebook_2_network_creation_and_exploration.pdf`
  contains examples to generate, visualize, and explore networks.
- **Supplementary File S3**: `supplement_notebook_3_enrichment_analysis.pdf`
  contains examples for running, querying, and visualizing enrichment analysis.
- **Supplementary File S4**: `supplement_notebook_4_agn.pdf`
  contains the example workflow to generate AGN.

##### 1.3 Animations and Videos

- **Supplementary File S5**: `supplement_agn_trajectory.gif`
  Network demonstrating aggregation of ontology terms performed at each time point (time-series animation). The area of each node is proportional to the enrichment of that ontology term at the indicated time point. The thickness of edges is related to the number of edges connecting the molecular nodes underlying the two term nodes (not time-dependent).

#### Methods

##### Enriched term compression

We created a method that filters the number of enriched terms based on redundancy in gene content, starting with a ranked array with columns term_name, rank, and set of genes. We then rank all terms based on either their combined score (from EnrichR), number of genes in the term, or p-value. Starting from the highest ranked, we compare all lower ranked terms and remove them if their similarity is above a user defined threshold. We use the Jaccard index[1], which is the size of the intersection divided by the size of the union of any two sets, as our similarity metric.

Using this approach, we are able to minimize the total number of terms while maintaining their enrichment content, as shown in 1 and demonstrated in Figure **S3**. In the bendamustine dataset described in the main manuscript, 84 terms were filtered to 17 terms, a reduction of 80%. It should also be noted that we can take the compressed terms and back calculate from the original enrichment array which terms were removed, as demonstrated in Figure **S3** C.

**Algorithm 1** Enrichment compression

```
procedure FILTER
    entry = [term, rank, geneSet]
    array = sorted list of entries                          ▷ EnrichmentResult class
    to_remove = set()
    for i in array do
        for j in array not including i do
            score = jaccard_index(i.geneSet,j.geneSet)
            if score > threshold then                       ▷ i contains more information than j
                to_remove.add(j)                            ▷ remove to_remove from array
```

##### Generation of Annotated Gene Set networks

Enriched terms provide us with big picture information about processes and when they occur, yet they do not provide us with details about the interconnection among them. *Why is DNA damage repair up regulated at 1 hour and apoptosis up-regulated at 24? Is there a cause and effect? Are they independent?* To answer these questions, we created an algorithm to extract trimmed signaling networks derived from the enriched terms. The terms can be selected based on expert knowledge, rank of enrichment, all compressed terms, or any other criteria.

Once the terms are selected, we find the nodes in the network for each term. From there, we search through all possible combinations of pairs between the terms. For example, if term 1 has genes (A, B, C) and term 2 has (D, E), we count the number of edges from the possible sets ((A, D), (A, E), (B, D), (B, E), (C, D), (C, E)) that are found in the network edges. We then do the reverse (term 2 to term 1). If a node is in both sets, we consider only the edges that connect the other term, not edges that are within the term. This is demonstrated in Figure **S5** (N). This results in one coarse-grained network, where nodes are terms and the edges are frequency of edges between any two terms, and the fine-grained network, which contains the species and the connections between them. By using both, we are able explore at a high level and zoom in on detail when required.

#### 2 Supplementary Tables

| identifier | label | type | significant | fold_change | p_value | source | sample_id |
|---|---|---|---|---|---|---|---|
| BAX | BAX_S(ph)292 | gene | True | 4 | 0.01 | SILAC | 01hr |
| HMDB00012 | Deoxyuridine | metabolite | True | -2 | 0.01 | HILIC | 02hr |

**Supplementary Table S1**: Example input data for MAGINE. Data are stored in a comma-separated value (CSV) file.

| | Name | Description |
|---|---|---|
| Function | load_data | Load MAGINE formatted csv |
| | create_summary_table | Create summary of counts per experimental method and sample_id |
| | subset | Filter data based on list of species |
| | require_n_sig | Filter data to include species that were measured at least N times |
| Plots | heatmap | Create heatmap or clustermap of species |
| | volcano_plot | Create volcano plot |
| | volcano_by_sample | Create a series of volcano plots for each sample |
| | plot_species | Plot scatter plots of selected species |
| | plot_histogram | Create histogram of fold change values |
| Properties | sig | Filter data to include only significant flagged entries |
| | up | Filter data to include fold change >0 |
| | down | Filter data to include fold change <0 |
| | id_list | Compile a list of unique species from sample |
| | by_sample | Generate list of unique species for each sample |

**Supplementary Table S2**: Categorized list of key functions in MAGINE's `data` module.

| | Name | Description |
|---|---|---|
| Functions | load_enrichment | Load MAGINE created enrichment analysis csv |
| | run | Run enrichment analysis for list of genes across selected gene sets |
| | run_samples | Run enrichment analysis for multiple lists of genes across selected gene sets |
| | run_enrichment_for_project | Run enrichment for entire ExperimentalData instance |
| | require_n_sig | Filter data to include species that were measured at least N times |
| | filter_multi | Filter by multiple criteria (p_value, combined_score, database, etc.) |
| | find_similar_terms | Rank order terms by similar gene sets |
| | filter_based_on_words | Filter by key words |
| | all_genes_from_df | Generate list of all genes in current EnrichmentResult instance |
| | term_to_genes | Generate list of genes for given term |
| | remove_redundant | Remove terms that are less enriched but highly similar |
| Plots | heatmap | Find all paths between list |
| | dist_matrix | Plot distance matrix of all terms |

**Supplementary Table S3**: Categorized list of key functions in MAGINE's `enrichment` module

| Name | Description |
| --- | --- |
| build_network | Generate network centered around provided species |
| create_subnetwork | Create annotated set network from enriched terms and background network |
| paths_between_pair | Find shortest path(s) between pair |
| paths_between_list | Find all paths between list |
| paths_between_two_lists | Find paths between two lists |
| neighbors | Find neighbors (upstream and/or downstream) of node |
| expand_neighbors | Add neighbors of node to network |
| draw | Draw using igraph, matplotlib, graphviz, cytoscape.js |
| RenderModel | Create Cytoscape instance of network through py2cytoscape |

**Supplementary Table S4**: Categorized list of key functions in MAGINE's `network` module

### 3 Supplementary Figures

a

| | 30s | 30m | 01h | 03h | 06h | 12h | 18h | 24h | 36h | 48h | 60h | 72h |
|---|---|---|---|---|---|---|---|---|---|---|---|---|
| Label-free | ✓ | ✓ | ✓ | ✓ | ✓ | ✓ | ✓ | ✓ | ✓ | ✓ | ✓ | ✓ |
| phSILAC | ✓ | - | ✓ | - | ✓ | ✓ | - | ✓ | - | - | - | - |
| SILAC | ✓ | - | ✓ | - | ✓ | ✓ | - | ✓ | - | - | - | - |
| RNAseq | - | - | ✓ | - | ✓ | ✓ | - | ✓ | - | - | - | - |

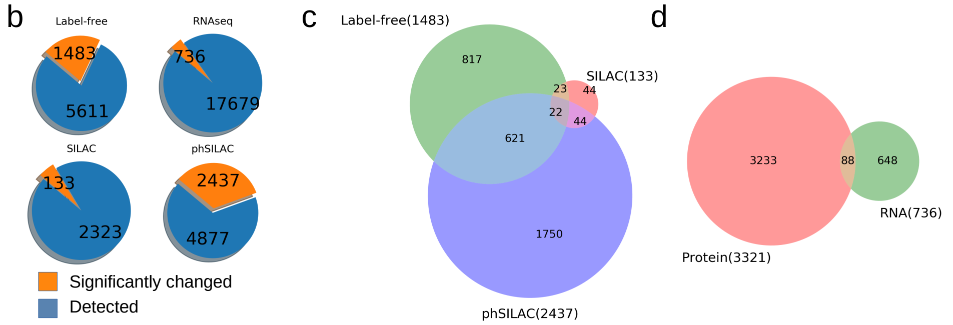


**Supplementary Fig. S1**: **Overlap between experimental measurements.** a.) Table of time points per experimental platform that were measured for the bendamustine study. b.) Summary of number of significant changed species vs detected. c.) Venn diagram comparing phSILAC, label-free, and SILAC. d.) Venn diagram comparing proteomics (phSILAC, label-free, SILAC) vs RNAseq.

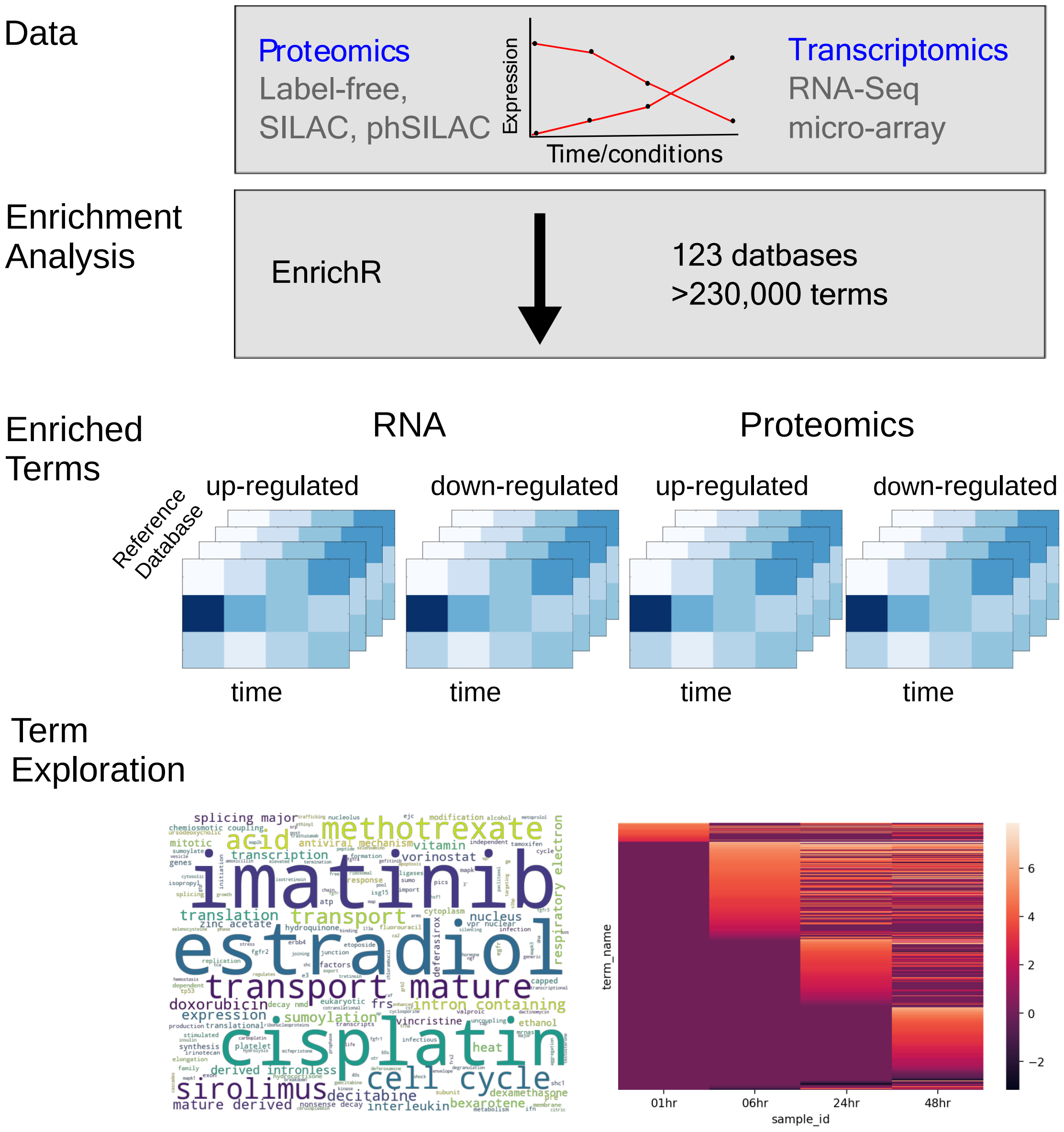


**Supplementary Fig. S2**: **Time series enrichment analysis pipeline**. Abstract representation of the automation of running enrichment analysis on multiple time points and experimental platforms. We utilize EnrichR to access various databases. We then organize the data. This allows us to explore the output in various ways. We can use word clouds to compress various databases to see common terms across them all. We can also plot enriched terms over times to see trends of biological processes.

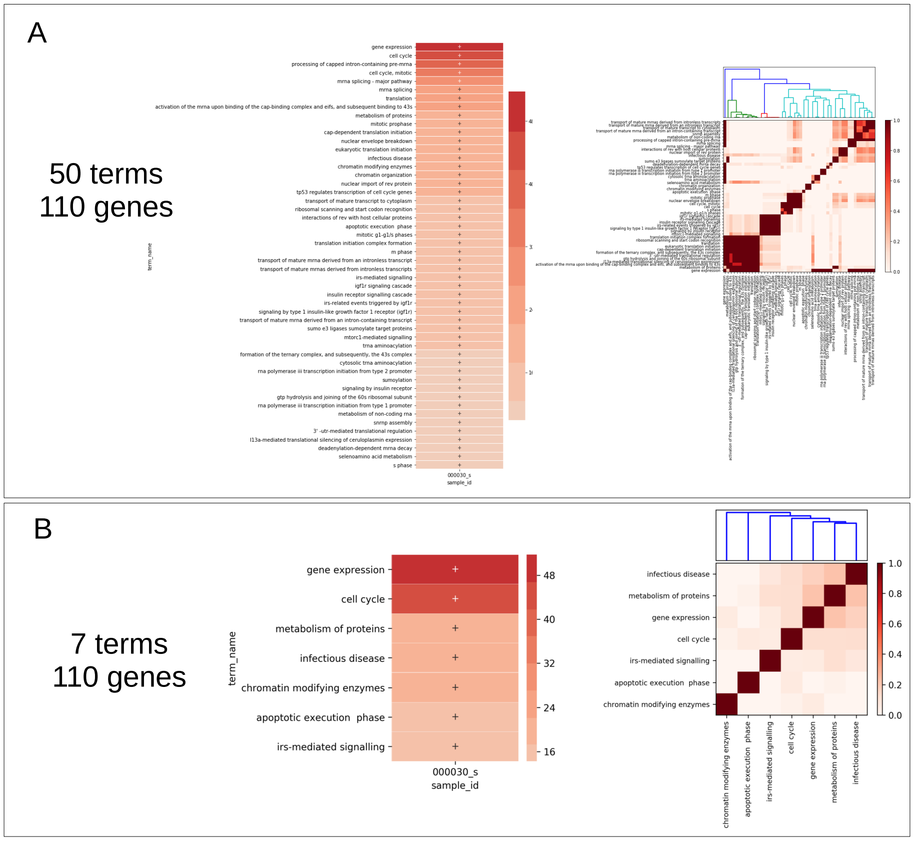


C

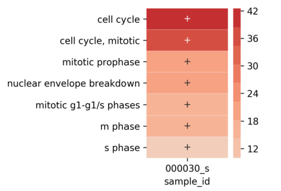


| term_name | genes |
|---|---|
| cell cycle | CLASP1,DYNC1H1,FAM175A,LMNA,LMNB1,MAPK1,MCM2,MCM4, MCM7,MLH1,NCAPH2,NEK7,NPM1,NUP107,NUP153,NUP188,PCM1, POLA1,PPP1R12A,PSMD6,RAD50,RB1,RBL1,RBL2,RUVBL1,SMC3, TOP2A,YWHAE |
| cell cycle, mitotic | CLASP1,DYNC1H1,LMNA,LMNB1,MAPK1,MCM2,MCM4,MCM7, NCAPH2,NEK7,NUP107,NUP153,NUP188,PCM1,POLA1,PPP1R12A, PSMD6,RB1,RBL1,RBL2,SMC3,TOP2A,YWHAE |
| mitotic prophase | LMNA,LMNB1,MAPK1,NCAPH2,NEK7,NUP107,NUP153,NUP188,RB1 |
| nuclear envelope breakdown | LMNA,LMNB1,NEK7,NUP107,NUP153,NUP188 |
| mitotic g1-g1/s phases | MCM2,MCM4,MCM7,POLA1,PSMD6,RB1,RBL1,RBL2,TOP2A |
| m phase | CLASP1,LMNA,LMNB1,MAPK1,NCAPH2,NEK7,NUP107,NUP153, NUP188,PSMD6,RB1,SMC3 |
| s phase | MCM2,MCM4,MCM7,POLA1,PSMD6,RB1,SMC3 |

**Supplementary Fig. S3**: **Demonstration of enrichment compression** A. Top 50 terms from Reactome for the 30 second phSILAC timepoint. The left figure shows the terms and their enriched values while the right shows the Jaccard index between all pairs. B. After filtering based on Jaccard index similarity. The resulting left figure has 7 terms, retaining all 110 genes that were contained in the terms from A. The right figure demonstrates that we have minimal overlap between corresponding terms. C. Demonstration of the collapsing of terms to a parent term. The left shows all terms that were collapsed from the term "cell cycle". The table on the right provides the genes that make up each term. Note that if a term was too broad ("cell cycle"), the user could remove it and repeat the process to maintain the level of detail which they require.

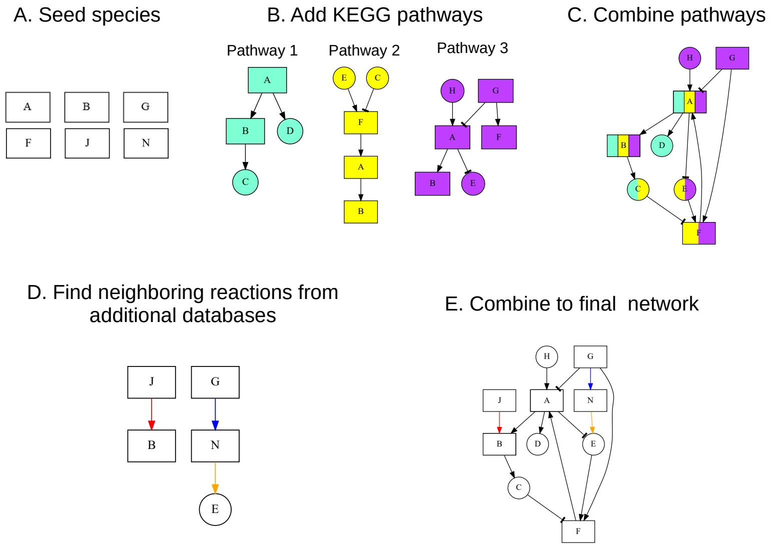


**Supplementary Fig. S4**: **Construction of data seeded network.** Example set of seed species to build network: (a) User provides seed species list; (b) KEGG pathways containing seed species are downloaded and merged; (c) Edges between seed species and nodes added by the KEGG pathways are identifier; (d) All nodes and edges are combined into a single network, (e).

| | Enrichment Value | Species |
|---|---|---|
| Term 1 | 15 | A, B, C |
| Term 2 | 10 | C, D, E |

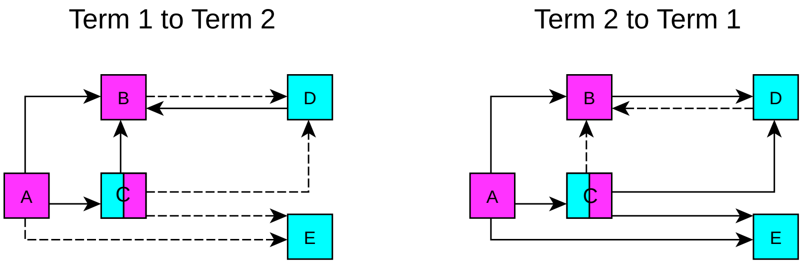


| | | Term 1 to Term 2 | Term 2 to Term 1 |
|---|---|---|---|
| # edges | Actual | 4 | 2 |
| | Possible | 6 | 6 |

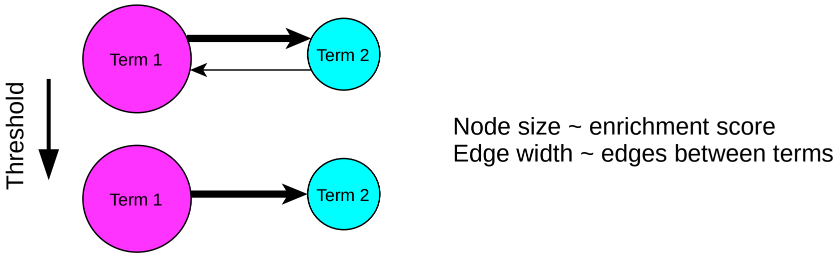


**Supplementary Fig. S5**: **Annotated gene set network construction** Nodes belonging to Term 1 are labeled as pink, Term 2 as blue, and nodes in both are colored both. We calculate the number of edges between nodes in Term 1 to Term 2 and Term 2 and Term 1 (dashed lines), shown in middle table. We then create nodes for each Term, with the size of the node corresponding to the enrichment value. Edges are created between the terms and width set according to the number of edges between the terms. Finally, we applied a minimum of 3 edges threshold between terms to arrive at our final network.

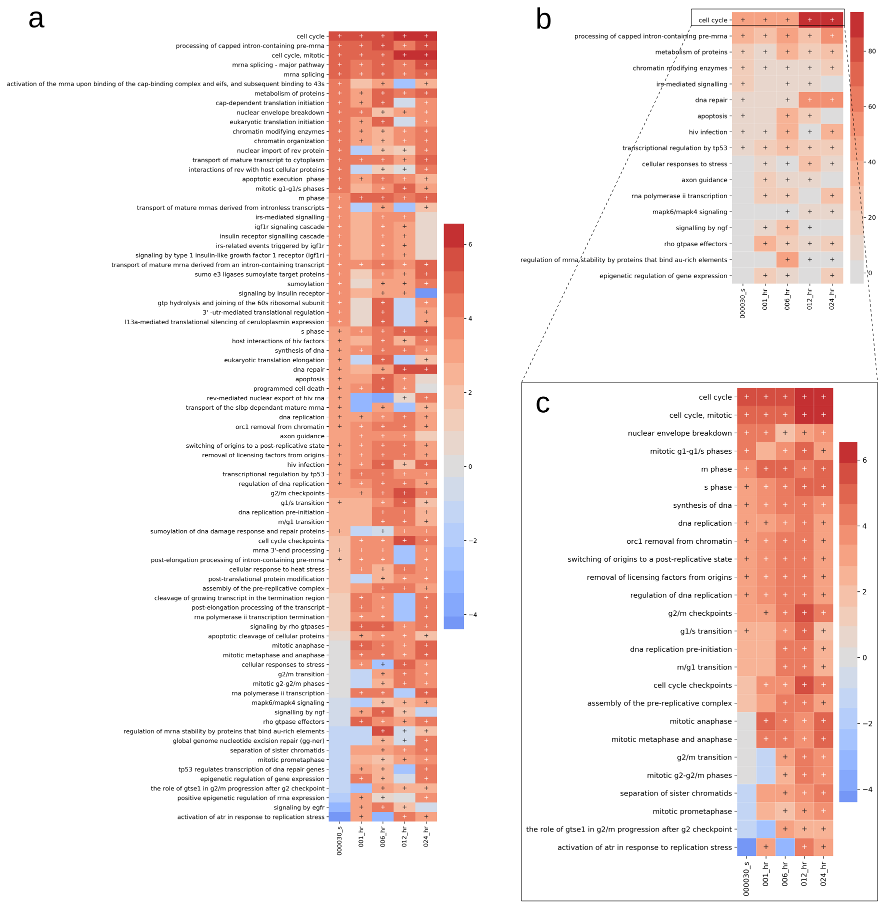


**Supplementary Fig. S6**: **MAGINE provides enrichment compression** A. Enrichment results for the Reactome gene set. The left figure shows the terms and their enriched values, while the right shows the Jaccard index between all pairs. B. Resulting array after filtering based on Jaccard index similarity. The resulting left figure has 17 terms. C. Terms that have high overlap with *cell cycle*. MAGINE not only provides a method to compress the information of the enrichment array, it also provides methods to extract more detailed, but less enriched terms.
