## Supplementary material for "A computational framework to explore cellular response mechanisms from multi-omics datasets": S1-Jupyter Notebook [Data Exploration]

### supplement_notebook_1_data_exploration

December 10, 2019

#### 1 Supplement notebook 1

##### 1.1 ExperimentalData class demonstration

This purpose of this notebook is to introduce and demonstrate the ExperimentalData.

First we import python packages that we need. Please refer to tutorials on Scipy, NumPy, Matplotlib, and Seaborn if unfamilar with these powerful tools.

[1]:
```
from IPython.display import display
%matplotlib inline
import pandas as pd
import matplotlib.pyplot as plt
import seaborn as sns
import numpy as np
from scipy.stats import pearsonr, spearmanr
```

[2]:
```
import magine.data.tools as dt
from magine.plotting.wordcloud_tools import create_wordcloud
from magine.plotting.venn_diagram_maker import create_venn2, create_venn3
```

##### 1.2 ExperimentalData class structure

Since MAGINE is built for multi-sample, multi-omics data, it is no surprise that the data is the most important aspect. Here we should how to use the :py:class:`ExperimentalData` class.

[3]:
```
# load the experimental data
from magine.data.experimental_data import load_data
```

[4]:
```
help(load_data)
```

```
Help on function load_data in module magine.data.experimental_data:

load_data(file_name, **kwargs)
    Load data into EnrichmentResult data class

    Parameters
    ----------
    file_name : str
    kwargs :
```

```
        Flags to pass to pandas.

    Returns
    -------
    df : EnrichmentResult
```

[5]:
```python
exp_data = load_data(
    'Data/bendamustine.csv.gz', # filename and location
    # Following args are passed to pandas.read_csv
    low_memory=False,
    index_col=0
)
```

##### 1.2.1 Getting counts from data

First, lets quickly view the stats about the data.

[6]:
```python
display(exp_data.create_summary_table())
display(exp_data.create_summary_table(index='label'))
```

```
sample_id  000030_s 00030_min  001_hr 003_hr  006_hr  012_hr 018_hr  024_hr  \
source
C18            5735      5114    5721   5834    6313    6574   6201    4531
HILIC         11891      9412   11880   7882   14215   14702  12666   10451
label_free     3215      3451    3113   4098    2907    3150   4273    4374
ph_silac       3240         -    3495      -    3327    3756      -    3212
rna_seq           -         -   16550      -   15887   16017      -   16418
silac          1629         -    1883      -    1761    1650      -    1664

sample_id  036_hr 048_hr 060_hr 072_hr  Total Unique Across
source
C18          7825   6267   4751   4773                19570
HILIC       11902  13013  10804   6350                27959
label_free   4100   4448   4188   2628                 5611
ph_silac        -      -      -      -                 4877
rna_seq         -      -      -      -                17679
silac           -      -      -      -                 2323


sample_id  000030_s 00030_min  001_hr 003_hr  006_hr  012_hr 018_hr  024_hr  \
source
C18            5629      5014    5626   5729    6188    6454   6074    4458
HILIC         11754      9314   11758   7781   14038   14530  12493   10322
label_free     3730      4113    3553   4717    3329    3645   4953    5058
ph_silac      12224         -   14512      -   12709   15472      -   12252
rna_seq           -         -   16550      -   15887   16017      -   16418
silac          1629         -    1883      -    1761    1650      -    1664
```

| sample_id | 036_hr | 048_hr | 060_hr | 072_hr | Total Unique Across |
|---|---|---|---|---|---|
| source | | | | | |
| C18 | 7707 | 6155 | 4670 | 4687 | 19212 |
| HILIC | 11746 | 12869 | 10684 | 6271 | 27572 |
| label_free | 4651 | 5263 | 4767 | 2911 | 7428 |
| ph_silac | - | - | - | - | 25613 |
| rna_seq | - | - | - | - | 17679 |
| silac | - | - | - | - | 2323 |

From here, we can see that we have 12 time points and 6 experimental platforms for the data. This is of all the data. We can filter by significantly measured or by looking at `label` column (default is `identifier`

[7]:
```python
display(exp_data.create_summary_table(sig=True))
display(exp_data.create_summary_table(sig=True, index='label'))
```

| sample_id | 000030_s | 00030_min | 001_hr | 003_hr | 006_hr | 012_hr | 018_hr | 024_hr | \ |
|---|---|---|---|---|---|---|---|---|---|
| source | | | | | | | | | |
| C18 | 870 | 121 | 454 | 555 | 444 | 322 | 293 | 341 | |
| HILIC | 729 | 452 | 226 | 891 | 354 | 1053 | 732 | 410 | |
| label_free | 14 | 18 | 22 | 37 | 113 | 18 | 85 | 161 | |
| ph_silac | 609 | - | 883 | - | 1091 | 722 | - | 944 | |
| rna_seq | - | - | 51 | - | 51 | 69 | - | 611 | |
| silac | 20 | - | 30 | - | 19 | 20 | - | 58 | |

| sample_id | 036_hr | 048_hr | 060_hr | 072_hr | Total Unique Across |
|---|---|---|---|---|---|
| source | | | | | |
| C18 | 1032 | 684 | 787 | 1224 | 5414 |
| HILIC | 83 | 2118 | 116 | 137 | 6244 |
| label_free | 39 | 162 | 853 | 542 | 1483 |
| ph_silac | - | - | - | - | 2437 |
| rna_seq | - | - | - | - | 736 |
| silac | - | - | - | - | 133 |

| sample_id | 000030_s | 00030_min | 001_hr | 003_hr | 006_hr | 012_hr | 018_hr | 024_hr | \ |
|---|---|---|---|---|---|---|---|---|---|
| source | | | | | | | | | |
| C18 | 840 | 119 | 443 | 542 | 435 | 314 | 289 | 335 | |
| HILIC | 699 | 437 | 215 | 866 | 353 | 1044 | 713 | 394 | |
| label_free | 14 | 18 | 22 | 37 | 114 | 18 | 86 | 168 | |
| ph_silac | 755 | - | 1189 | - | 1525 | 975 | - | 1570 | |
| rna_seq | - | - | 51 | - | 51 | 69 | - | 611 | |
| silac | 20 | - | 30 | - | 19 | 20 | - | 58 | |

| sample_id | 036_hr | 048_hr | 060_hr | 072_hr | Total Unique Across |
|---|---|---|---|---|---|
| source | | | | | |
| C18 | 1014 | 670 | 776 | 1198 | 5296 |

```
HILIC            83    2094    115     134                     6138
label_free       39     170    925     591                     1653
ph_silac          -       -      -       -                     5115
rna_seq           -       -      -       -                      736
silac             -       -      -       -                      133
```

The `.species` index aggregates all data. Since we utilize a `pandas.DataFrame`, we can use the `.head` method to glance at the data.

[8]:
```
exp_data.species.head(5)
```

[8]:
```
  identifier           label  fold_change  significant  p_value species_type  \
0       UBA6      UBA6_silac    -1.049913        False      1.0      protein
1       MTDH      MTDH_silac    -1.038867        False      1.0      protein
2   SLC25A24  SLC25A24_silac     1.014615        False      1.0      protein
3    ANKRD22   ANKRD22_silac    -1.058937        False      1.0      protein
4        AGK       AGK_silac     1.001600        False      1.0      protein

  sample_id source
0    001_hr  silac
1    001_hr  silac
2    001_hr  silac
3    001_hr  silac
4    001_hr  silac
```

We can filter the data by `source` using the `.name`, where name is anything in the `source` column. We can get a list of these by printing `exp_data.exp_methods`

[9]:
```
exp_data.exp_methods
```

[9]:
```
['silac', 'ph_silac', 'HILIC', 'C18', 'label_free', 'rna_seq']
```

[10]:
```
# filters to only the 'label_free'
exp_data.label_free.shape
```

[10]:
```
(50736, 8)
```

[11]:
```
exp_data.label_free.head(5)
```

[11]:
```
       identifier                  label  fold_change  significant  p_value  \
515804    RHOXF2B             RHOXF2B_lf         1.51         True   0.0003
515805      KIF11               KIF11_lf         1.70         True   0.0008
515806    MYBBP1A  MYBBP1A_S(ph)1163_lf         1.40        False   0.0009
515807      CDC20               CDC20_lf         2.78         True   0.0012
515808       TMPO        TMPO_T(ph)160_lf        -1.47        False   0.0016

       species_type sample_id      source
515804      protein    036_hr  label_free
515805      protein    036_hr  label_free
515806      protein    036_hr  label_free
515807      protein    036_hr  label_free
515808      protein    036_hr  label_free
```

[12]:
```
exp_data.rna_seq.head(5)
```

[12]:
```
         identifier                label  fold_change  significant   p_value  \
566540        CDC20         CDC20_rnaseq    -1.528232         True  0.016321
566541         ASPM          ASPM_rnaseq    -1.346689        False  0.038802
566542  FO538757.2  FO538757.2_rnaseq     2.133888         True  0.038802
566543         GNL3          GNL3_rnaseq    -2.313025         True  0.016321
566544      SNORD19       SNORD19_rnaseq    -2.313025         True  0.016321

       species_type sample_id   source
566540      rna_seq    012_hr  rna_seq
566541      rna_seq    012_hr  rna_seq
566542      rna_seq    012_hr  rna_seq
566543      rna_seq    012_hr  rna_seq
566544      rna_seq    012_hr  rna_seq
```

##### 1.2.2 Significant filter

We can use the `significant` column to filter that data to only contain those species.

[13]:
```
exp_data.species.shape
```

[13]:
```
(558025, 8)
```

[14]:
```
exp_data.species.sig.shape
```

[14]:
```
(24620, 8)
```

##### 1.2.3 Filter data to up or down regulated species.

For enrichment analysis, we will want to access up-regulated and down-regulated species using `.up` and `.down`.

[15]:
```
exp_data.rna_seq.up.head(5)
```

[15]:
```
            identifier                 label  fold_change  significant  \
566542      FO538757.2     FO538757.2_rnaseq     2.133888         True
566546  RP4-669L17.10  RP4-669L17.10_rnaseq     3.788399         True
566547   RP4-669L17.4   RP4-669L17.4_rnaseq     3.788399         True
566548     RNU6-513P      RNU6-513P_rnaseq     2.462533         True
566549          RRP7B          RRP7B_rnaseq     2.462533         True

         p_value species_type sample_id   source
566542  0.038802      rna_seq    012_hr  rna_seq
566546  0.029804      rna_seq    012_hr  rna_seq
566547  0.029804      rna_seq    012_hr  rna_seq
566548  0.016321      rna_seq    012_hr  rna_seq
566549  0.016321      rna_seq    012_hr  rna_seq
```

[16]:
```
exp_data.rna_seq.down.head(5)
```

[16]:
```
       identifier             label  fold_change  significant   p_value  \
566540      CDC20      CDC20_rnaseq    -1.528232         True  0.016321
566543       GNL3       GNL3_rnaseq    -2.313025         True  0.016321
566544    SNORD19    SNORD19_rnaseq    -2.313025         True  0.016321
566545   SNORD19B   SNORD19B_rnaseq    -2.313025         True  0.016321
566550     FAM73A     FAM73A_rnaseq    -9.644115         True  0.016321

       species_type sample_id   source
566540      rna_seq    012_hr  rna_seq
566543      rna_seq    012_hr  rna_seq
566544      rna_seq    012_hr  rna_seq
566545      rna_seq    012_hr  rna_seq
566550      rna_seq    012_hr  rna_seq
```

##### 1.2.4 Extracting by sample (time point)

We can filter by `sample_id`.

[17]:
```
exp_data.sample_ids
```

[17]:
```
['000030_s',
 '00030_min',
 '001_hr',
 '003_hr',
 '006_hr',
 '012_hr',
 '018_hr',
 '024_hr',
 '036_hr',
 '048_hr',
 '060_hr',
 '072_hr']
```

[18]:
```
exp_data['000030_s'].head(5)
```

[18]:
```
     identifier         label  fold_change  significant  p_value species_type  \
5200       UBA6    UBA6_silac    -1.088427        False      1.0      protein
5201     AKR1A1  AKR1A1_silac     1.065195        False      1.0      protein
5202     MTHFD2  MTHFD2_silac    -1.308449        False      1.0      protein
5203       DLAT    DLAT_silac     1.029963        False      1.0      protein
5204        CNP     CNP_silac    -1.244300        False      1.0      protein

     sample_id source
5200  000030_s  silac
5201  000030_s  silac
5202  000030_s  silac
5203  000030_s  silac
5204  000030_s  silac
```

##### 1.2.5 Calculating overlaps between time points

[19]:
```python
from itertools import combinations

def overlap(vals):
    return len(vals[0].intersection(vals[1]))

def calc_dist(names, gene_sets, figsize=(12, 12)):

    n_dim = len(names)
    scores = list(map(overlap, combinations(gene_sets, 2)))

    dist_mat = np.zeros((n_dim, n_dim), dtype=float)
    ind = 0
    for i in range(n_dim):
        for j in range(i, n_dim):
            if i == j:
                dist_mat[i, i] = np.nan
                continue
            elif i>= j:
                continue
            dist_mat[i, j] = scores[ind]
            dist_mat[j, i] = scores[ind]
            ind += 1

    fig = plt.figure(figsize=figsize)
    ax = fig.add_subplot(111)
    cmap=plt.cm.Reds
    cmap.set_under(".5")
    fig = sns.heatmap(dist_mat, cmap=cmap, fmt='3g', annot=True, linewidths=0.1,
                xticklabels=names, yticklabels=names, ax=ax, square=False)
    ax.xaxis.tick_top()
    ax.set_xticklabels(names, minor=False, rotation=90, fontsize=12)
    ax.set_yticklabels(names, minor=False, rotation=0, fontsize=12)


def overlap_by_source(source_name, figsize=(10, 10)):
    sample_ids = np.array(exp_data[source_name].sample_ids)
    sample_sets = exp_data[source_name].sig.by_sample
    calc_dist(sample_ids, sample_sets, figsize)

overlap_by_source('label_free')
overlap_by_source('rna_seq', figsize=(6, 6))
overlap_by_source('silac', figsize=(6, 6))
overlap_by_source('ph_silac', figsize=(4, 4))
plt.savefig("ph_silac_overlap_by_time.png", dpi=300, bbox_inches='tight')
```

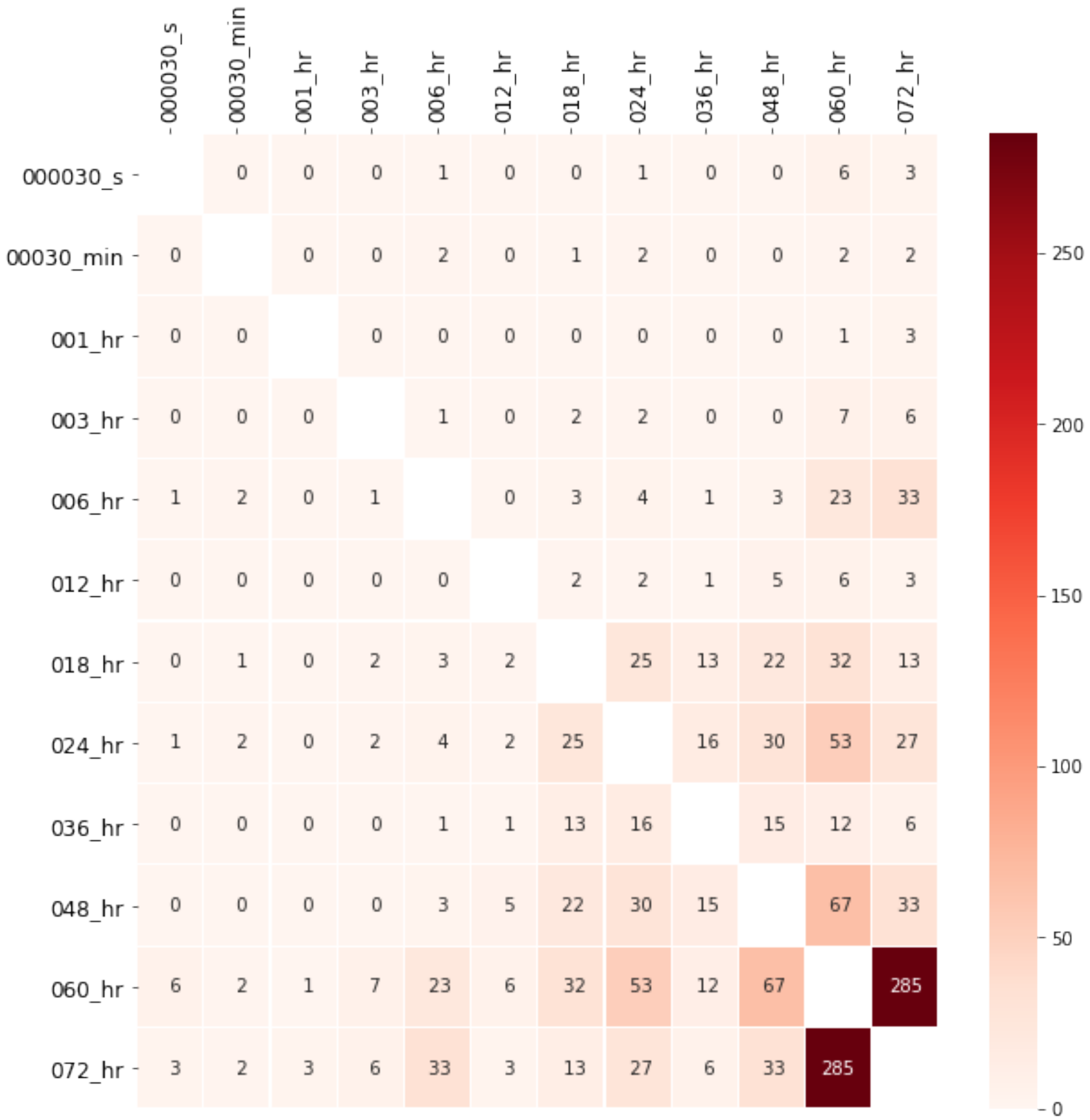
000030_s
00030_min
001_hr
003_hr
006_hr
012_hr
018_hr
024_hr
036_hr
048_hr
060_hr
072_hr
000030_s 0 0 0 1 0 0 1 0 0 6 3
00030_min 0 0 0 2 0 1 2 0 0 2 2
001_hr 0 0 0 0 0 0 0 0 0 1 3
003_hr 0 0 0 1 0 2 2 0 0 7 6
006_hr 1 2 0 1 0 3 4 1 3 23 33
012_hr 0 0 0 0 0 2 2 1 5 6 3
018_hr 0 1 0 2 3 2 25 13 22 32 13
024_hr 1 2 0 2 4 2 25 16 30 53 27
036_hr 0 0 0 0 1 1 13 16 15 12 6
048_hr 0 0 0 0 3 5 22 30 15 67 33
060_hr 6 2 1 7 23 6 32 53 12 67 285
072_hr 3 2 3 6 33 3 13 27 6 33 285
250
200
150
100
50
0

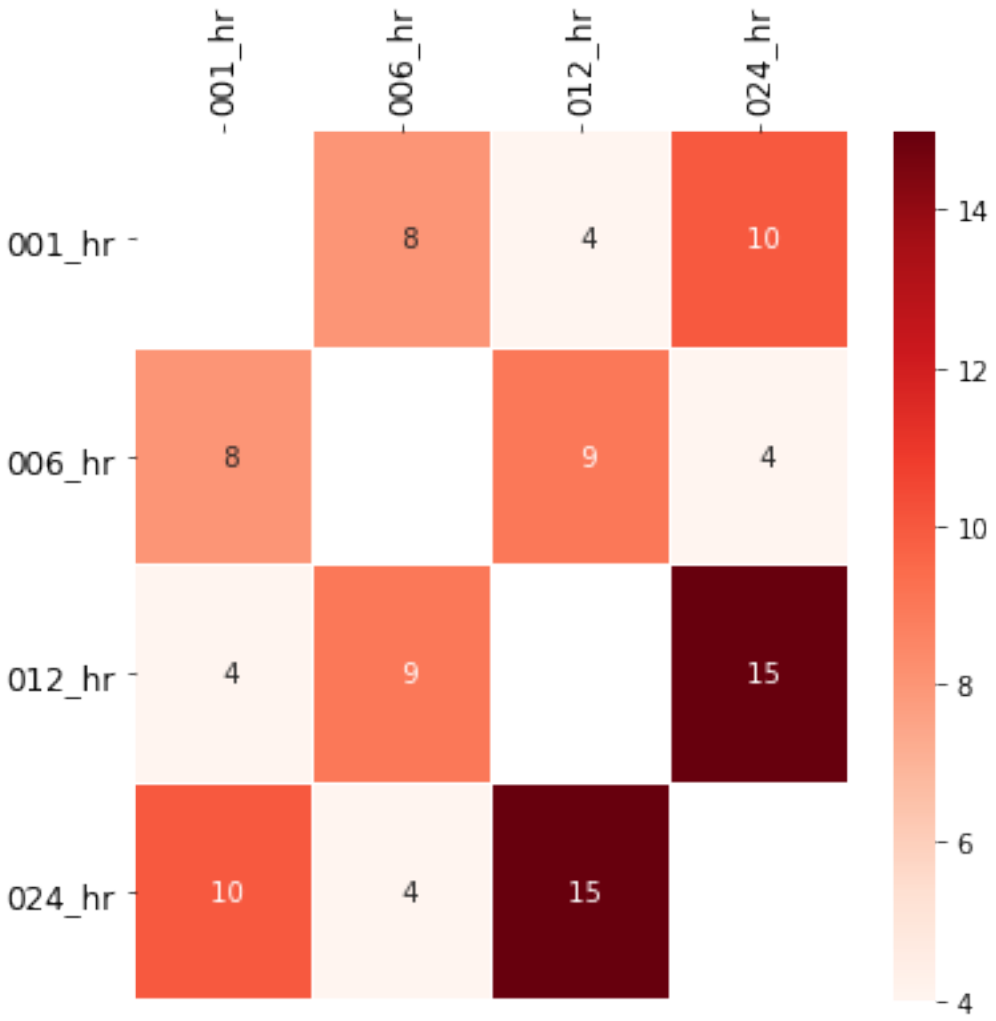
001_hr
006_hr
012_hr
024_hr
001_hr
006_hr
012_hr
024_hr
8
4
10
8
9
4
4
9
15
10
4
15
14
12
10
8
6
4

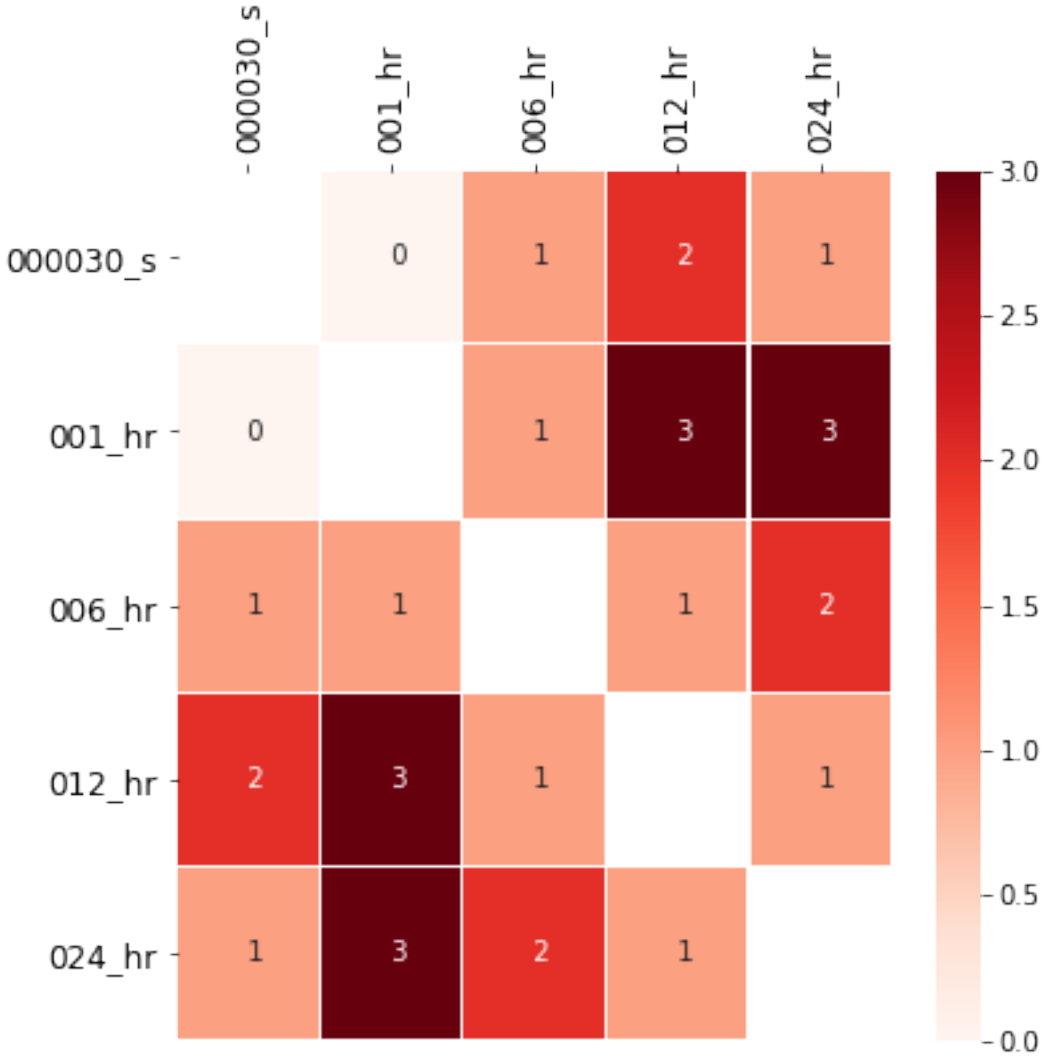
000030_s
001_hr
006_hr
012_hr
024_hr
0
1
2
3
0.0
0.5
1.0
1.5
2.0
2.5
3.0

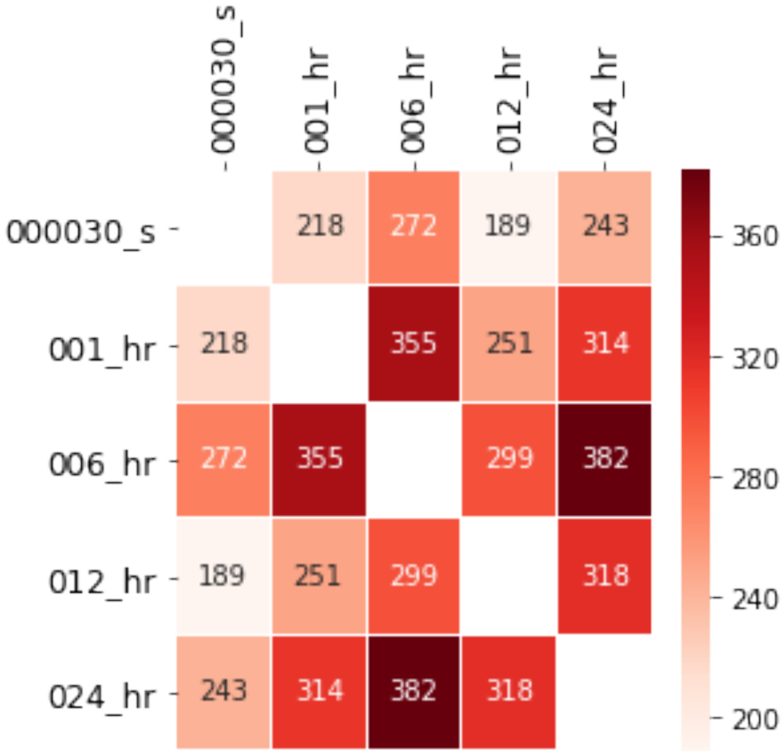


We can apply the same function to compare across experimental platforms.

[20]:
```
names, sets = [], []
for i in exp_data.exp_methods:
    sets.append(exp_data[i].sig.id_list)
    names.append(i)
calc_dist(names, sets, figsize=(4, 4))
plt.savefig("overlap_experiments.png", dpi=300, bbox_inches='tight')
```

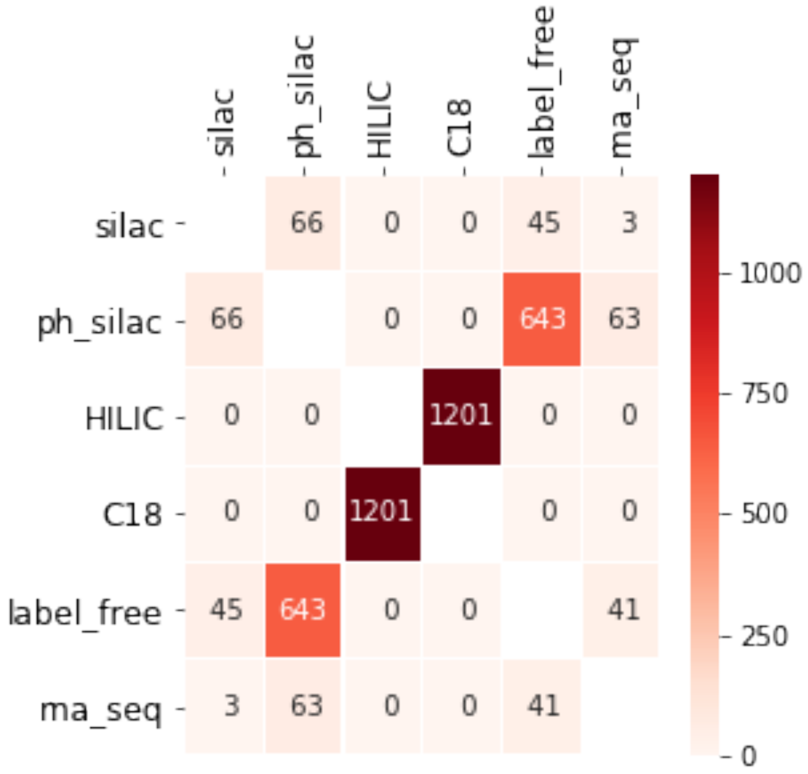


##### 1.2.6 Pivot table to get table across time

[21]:
```
exp_data.rna_seq.pivoter(
    convert_to_log=False,
    index='identifier',
    columns='sample_id',
    values=['fold_change', 'p_value']
).head(5)
```

[21]:
```
           fold_change                                  p_value          \
sample_id       001_hr    006_hr    012_hr    024_hr     001_hr   006_hr
identifier
7SK           1.139800 -1.148785 -1.014452  1.037751   0.999631  0.99995
A1BG         -2.402129  1.288824  1.610149  1.168061   0.999631  0.99995
A1BG-AS1     -1.081850  1.257409  1.056497 -1.099355   0.999631  0.99995
A2ML1        -1.372659  1.307877 -1.343714  1.793100   0.999631  0.99995
AAAS          1.203116 -1.058658 -1.012985  1.007011   0.999631  0.99995


sample_id     012_hr    024_hr
identifier
```

```
7SK          0.999660  0.864095
A1BG         0.803284  0.673174
A1BG-AS1     0.999660  0.752914
A2ML1        0.999660  0.274612
AAAS         0.999660  0.970896
```

[22]:
```python
exp_data.ph_silac.pivoter(
    convert_to_log=False,
    index='label',
    columns='sample_id',
    values=['fold_change', 'p_value']
).head(5)
```

[22]:
```
                      fold_change                                           \
sample_id                000030_s    001_hr    006_hr     012_hr    024_hr
label
A2M_1004_1014_phsilac         NaN -1.060800       NaN  43.747479       NaN
A2M_339_345_phsilac           NaN       NaN       NaN  -1.157100       NaN
AAAS_S(ph)495_phsilac    1.121971 -1.230439 -1.159737  -1.288961 -1.486965
AAGAB_134_161_phsilac    1.134301       NaN       NaN        NaN       NaN
AAGAB_274_284_phsilac         NaN       NaN       NaN  -1.123760       NaN

                       p_value
sample_id             000030_s 001_hr 006_hr 012_hr 024_hr
label
A2M_1004_1014_phsilac      NaN    1.0    NaN  0.049    NaN
A2M_339_345_phsilac        NaN    NaN    NaN  1.000    NaN
AAAS_S(ph)495_phsilac      1.0    1.0    1.0  1.000    1.0
AAGAB_134_161_phsilac      1.0    NaN    NaN    NaN    NaN
AAGAB_274_284_phsilac      NaN    NaN    NaN  1.000    NaN
```

Note that in the previous example, we find that there are NaN values. This is because there might be measurements missing in our experimental data. We can easy check what species are not found in all samples.

[23]:
```python
measured_in_all = exp_data.ph_silac.present_in_all_columns(
    index='label',
    columns='sample_id',
)
```

```
Number in index went from 25613 to 5595
```

This shows that out of the 25613 unique species measured in ph_silac proteomics, only 5595 were measured in all time points. What one can do with this information is dependent on the analysis. For now, we will keep using the full dataset.

[24]:
```python
measured_in_all.pivoter(
    convert_to_log=False,
    index='label',
    columns='sample_id',
    values=['fold_change', 'p_value']
```

```
).head(10)
```

[24]:

```
                                      fold_change                     \
sample_id                                000030_s    001_hr    006_hr
label
AAAS_S(ph)495_phsilac                    1.121971 -1.230439 -1.159737
AAGAB_S(ph)310_S(ph)311_phsilac         -0.031090  1.711100 -1.557812
AAK1_T(ph)606_phsilac                   -1.105709 -1.013440  1.033784
AAK1_T(ph)620_S(ph)623_phsilac           1.003735 -1.002832 -0.015575
AARS_(ca)_173_194_phsilac                1.039944  1.243000  1.254237
AARS_225_235_phsilac                     1.094279  1.107042  1.051156
AASDHPPT_253_267_phsilac                 1.007512  1.013656  1.002298
AATF_S(ph)316_S(ph)320_S(ph)321_phsilac -0.020251 -1.538099  1.119900
ABCE1_(ox)_542_557_phsilac              -1.254600 -1.226600  1.005787
ABCE1_213_224_phsilac                    1.072050 -1.006802  1.097534

                                                         p_value         \
sample_id                                  012_hr    024_hr 000030_s 001_hr
label
AAAS_S(ph)495_phsilac                   -1.288961 -1.486965      1.0  1.000
AAGAB_S(ph)310_S(ph)311_phsilac         -0.000927 -1.657294      1.0  0.049
AAK1_T(ph)606_phsilac                   -2.708685 -2.866159      1.0  1.000
AAK1_T(ph)620_S(ph)623_phsilac          -1.082963 -1.051561      1.0  1.000
AARS_(ca)_173_194_phsilac               -1.347523 -1.034186      1.0  1.000
AARS_225_235_phsilac                    -1.199240  1.101794      1.0  1.000
AASDHPPT_253_267_phsilac                -1.222638  1.148329      1.0  1.000
AATF_S(ph)316_S(ph)320_S(ph)321_phsilac -1.107462 -1.393000      1.0  0.049
ABCE1_(ox)_542_557_phsilac              -1.732200  1.001636      1.0  1.000
ABCE1_213_224_phsilac                   -1.155031 -1.085661      1.0  1.000


sample_id                               006_hr 012_hr 024_hr
label
AAAS_S(ph)495_phsilac                    1.000  1.000  1.000
AAGAB_S(ph)310_S(ph)311_phsilac          0.049  1.000  1.000
AAK1_T(ph)606_phsilac                    1.000  0.049  0.049
AAK1_T(ph)620_S(ph)623_phsilac           1.000  1.000  1.000
AARS_(ca)_173_194_phsilac                1.000  1.000  1.000
AARS_225_235_phsilac                     1.000  1.000  1.000
AASDHPPT_253_267_phsilac                 1.000  1.000  1.000
AATF_S(ph)316_S(ph)320_S(ph)321_phsilac  1.000  1.000  1.000
ABCE1_(ox)_542_557_phsilac               1.000  0.049  1.000
ABCE1_213_224_phsilac                    1.000  1.000  1.000
```

We can use this same principle and filter that data by requiring a species to be significant in `n` `sample_id`.

[25]:

```
lf_4_tp = exp_data.label_free.require_n_sig(
    index='label',
```

```
    columns='sample_id',
    n_sig=4
)
display(lf_4_tp.head(10))
```

```
       identifier                      label  fold_change  significant  \
515805      KIF11                   KIF11_lf         1.70         True
515807     CDC20                    CDC20_lf         2.78         True
515810      CKAP2                   CKAP2_lf         1.63         True
515813       TPX2                    TPX2_lf         1.93         True
515814      KIFC1                   KIFC1_lf         1.73         True
515822       RRM2                    RRM2_lf         1.92         True
515825     NUSAP1                  NUSAP1_lf         1.92         True
515827      CDCA5                   CDCA5_lf         1.76         True
515831      UBE2S                   UBE2S_lf         1.52         True
515835   HISTIH1B  HIST1H1B_N-term S(ace)2_lf        -1.51         True

        p_value species_type sample_id      source
515805   0.0008      protein    036_hr  label_free
515807   0.0012      protein    036_hr  label_free
515810   0.0024      protein    036_hr  label_free
515813   0.0038      protein    036_hr  label_free
515814   0.0040      protein    036_hr  label_free
515822   0.0075      protein    036_hr  label_free
515825   0.0087      protein    036_hr  label_free
515827   0.0108      protein    036_hr  label_free
515831   0.0120      protein    036_hr  label_free
515835   0.0131      protein    036_hr  label_free
```

##### 1.2.7 Visualization

**Volcano plots**

[26]:
```
exp_data.label_free.volcano_plot();
```

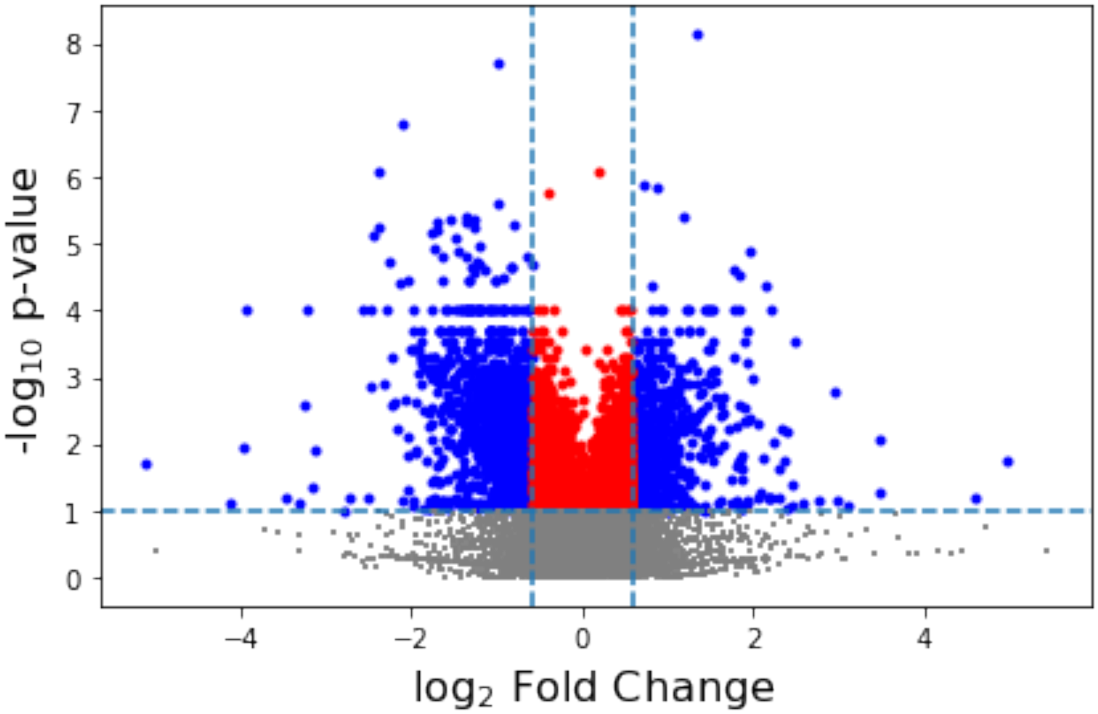


[27]:
```
exp_data.label_free.volcano_by_sample(sig_column=True);
```

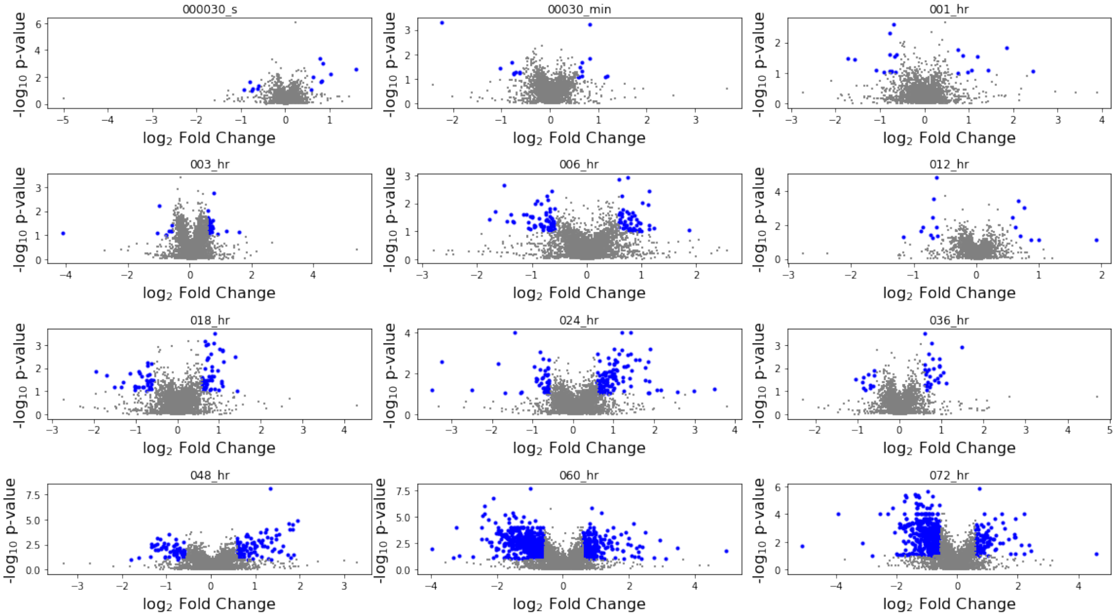


**Histogram**

[28]:
```
exp_data.label_free.plot_histogram();
```

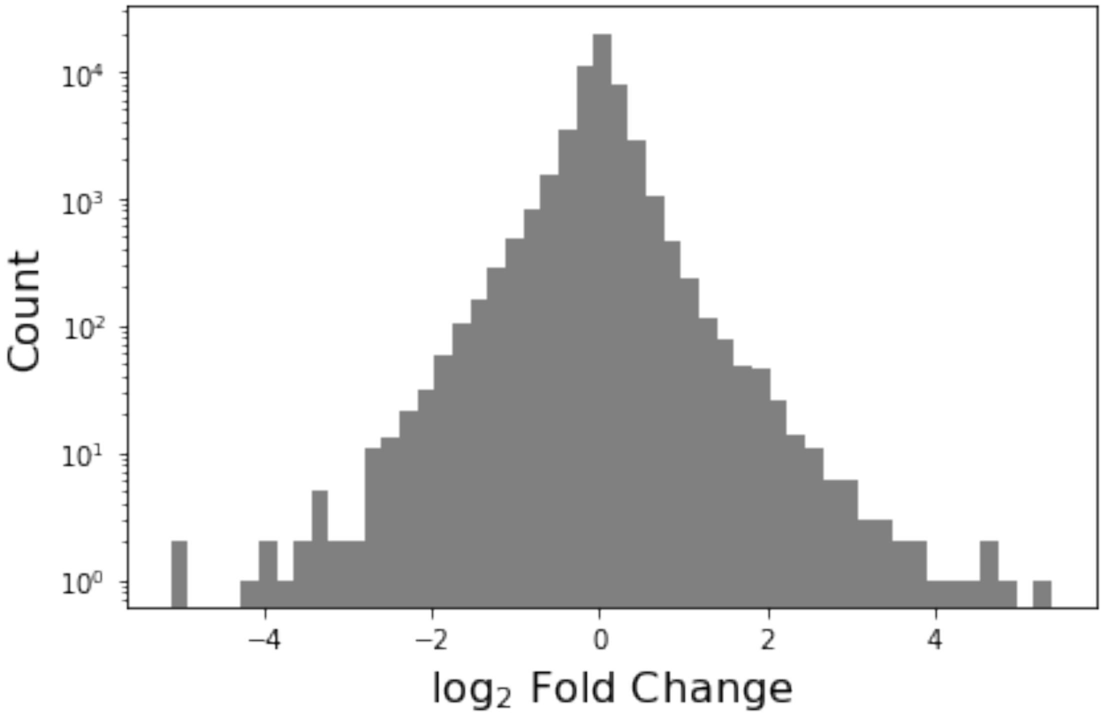


**Plotting subset of species** We provide the a few plotting interfaces to explore that subsets of the data. Basically, you create a list of species and provide it to the function. It filters based on these and then returns the results.

**Time series using ploty and matplotlib**

[29]:
```
# sample list for demo purposes
interesting_list = ['CCNA2', 'CDCA5', 'CDC20', 'AURKA', 'AURKB']

exp_data.label_free.plot_species(interesting_list, plot_type='matplotlib');
```

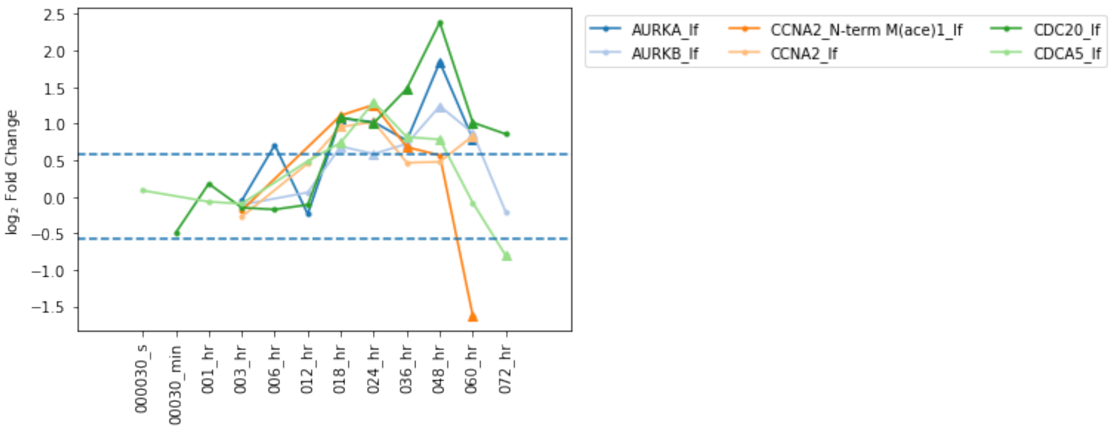


[30]:
```
exp_data.label_free.plot_species(interesting_list, plot_type='plotly')
```

**Heatplots**

[31]:
```
exp_data.label_free.heatmap(interesting_list, linewidths=0.01, figsize=(4,3));
```

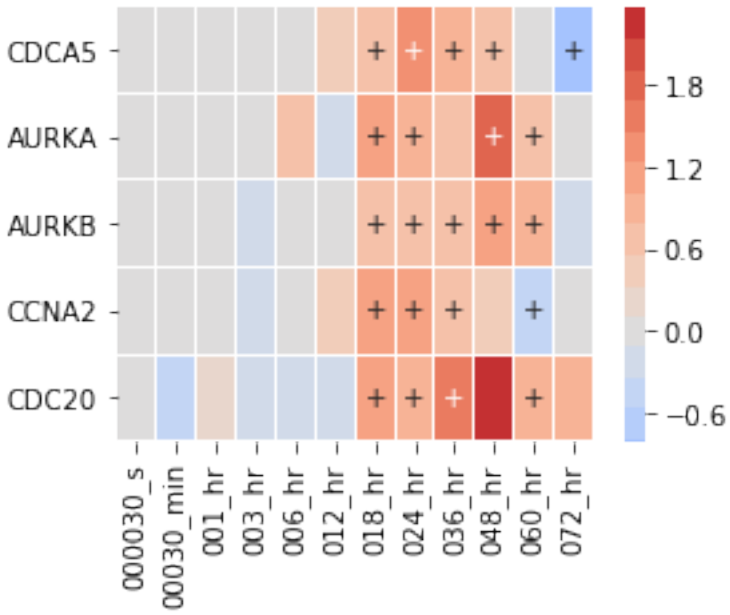


Notice that the above plot doesn't show any of the modifiers of `CCNA2_N-term M(ace)1_lf`. This is because the default index to pivot plots is the `identifier` column. You can set the `label` column for plotting by passing index=`label` to the function. Note, if you want to filter the data using the more generic `identifier` column, you just specify that with subset_index=`identifier`

[32]:
```
exp_data.label_free.heatmap(
    interesting_list,
```

```
    index='label',
    subset_index='identifier',
    linewidths=0.01,
    figsize=(4,3)
);
```

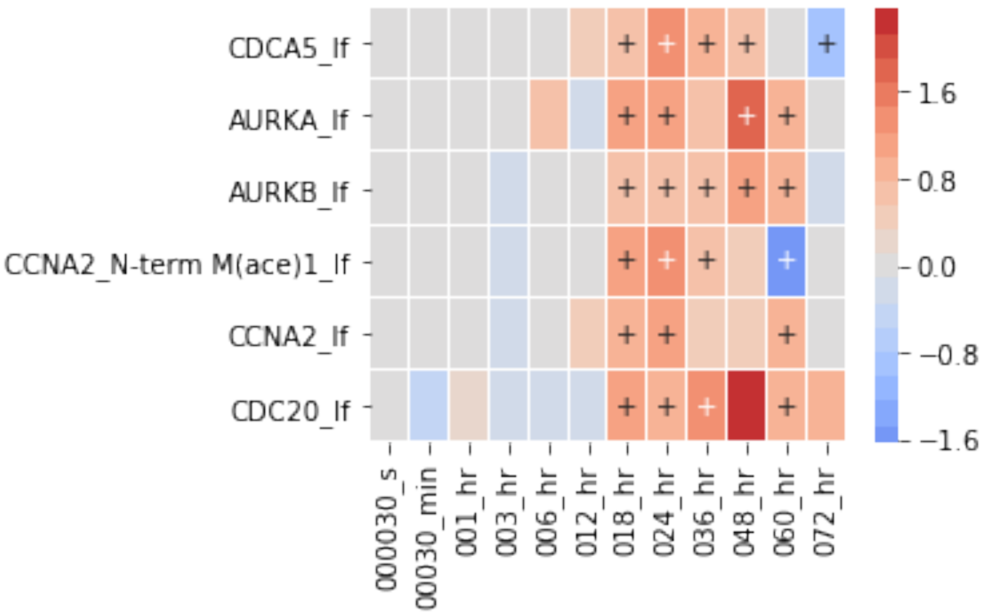


##### 1.2.8 Examples

Here are a few examples how all the above commands can be chained together to create plots with varying degrees of critera.

**Query 1:**

`Heatmap of label-free proteomics that are signficantly change in at least 3 time points. Extract clusters and visualize separately.`

[33]:
```
lf_sig = exp_data.label_free.require_n_sig(
    index='label',
    columns='sample_id',
    n_sig=3
)
fig = lf_sig.heatmap(
    convert_to_log=True,
    cluster_row=True,
    index='label',
    values='fold_change',
```

```
    columns='sample_id',
    annotate_sig=True,
    figsize=(8, 12),
    div_colors=True,
    num_colors=21,
    linewidths=0.01
);
for i, j in fig.row_clusters.items():
    lf_sig.heatmap(j, index='label', linewidths=0.01, figsize=(6, 8))
    plt.show()
```

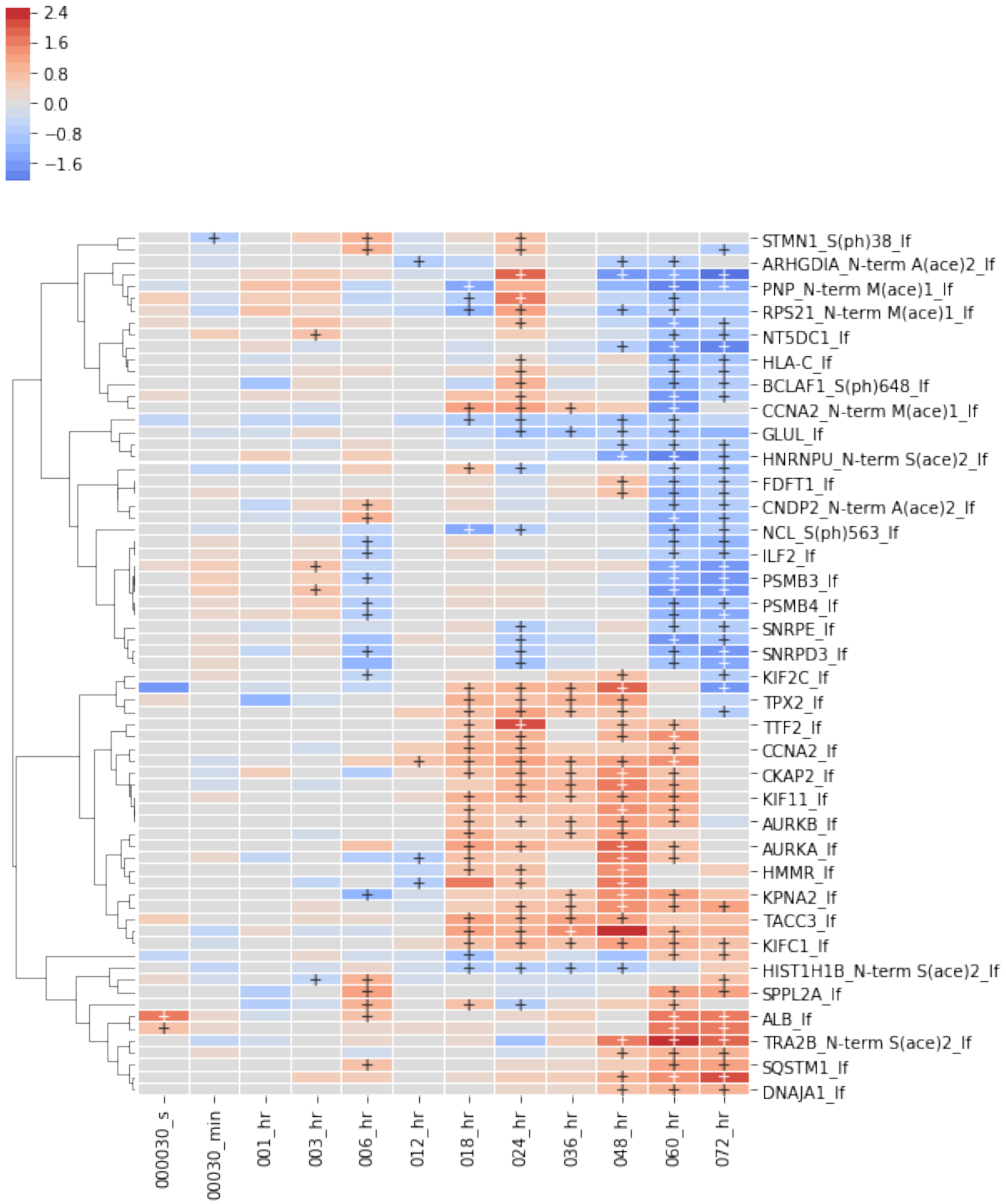

2.4
1.6
0.8
0.0
−0.8
−1.6
STMN1_S(ph)38_lf
ARHGDIA_N-term A(ace)2_lf
PNP_N-term M(ace)1_lf
RPS21_N-term M(ace)1_lf
NT5DC1_lf
HLA-C_lf
BCLAF1_S(ph)648_lf
CCNA2_N-term M(ace)1_lf
GLUL_lf
HNRNPU_N-term S(ace)2_lf
FDFT1_lf
CNDP2_N-term A(ace)2_lf
NCL_S(ph)563_lf
ILF2_lf
PSMB3_lf
PSMB4_lf
SNRPE_lf
SNRPD3_lf
KIF2C_lf
TPX2_lf
TTF2_lf
CCNA2_lf
CKAP2_lf
KIF11_lf
AURKB_lf
AURKA_lf
HMMR_lf
KPNA2_lf
TACC3_lf
KIFC1_lf
HIST1H1B_N-term S(ace)2_lf
SPPL2A_lf
ALB_lf
TRA2B_N-term S(ace)2_lf
SQSTM1_lf
DNAJA1_lf
000030_s
00030_min
001_hr
003_hr
006_hr
012_hr
018_hr
024_hr
036_hr
048_hr
060_hr
072_hr

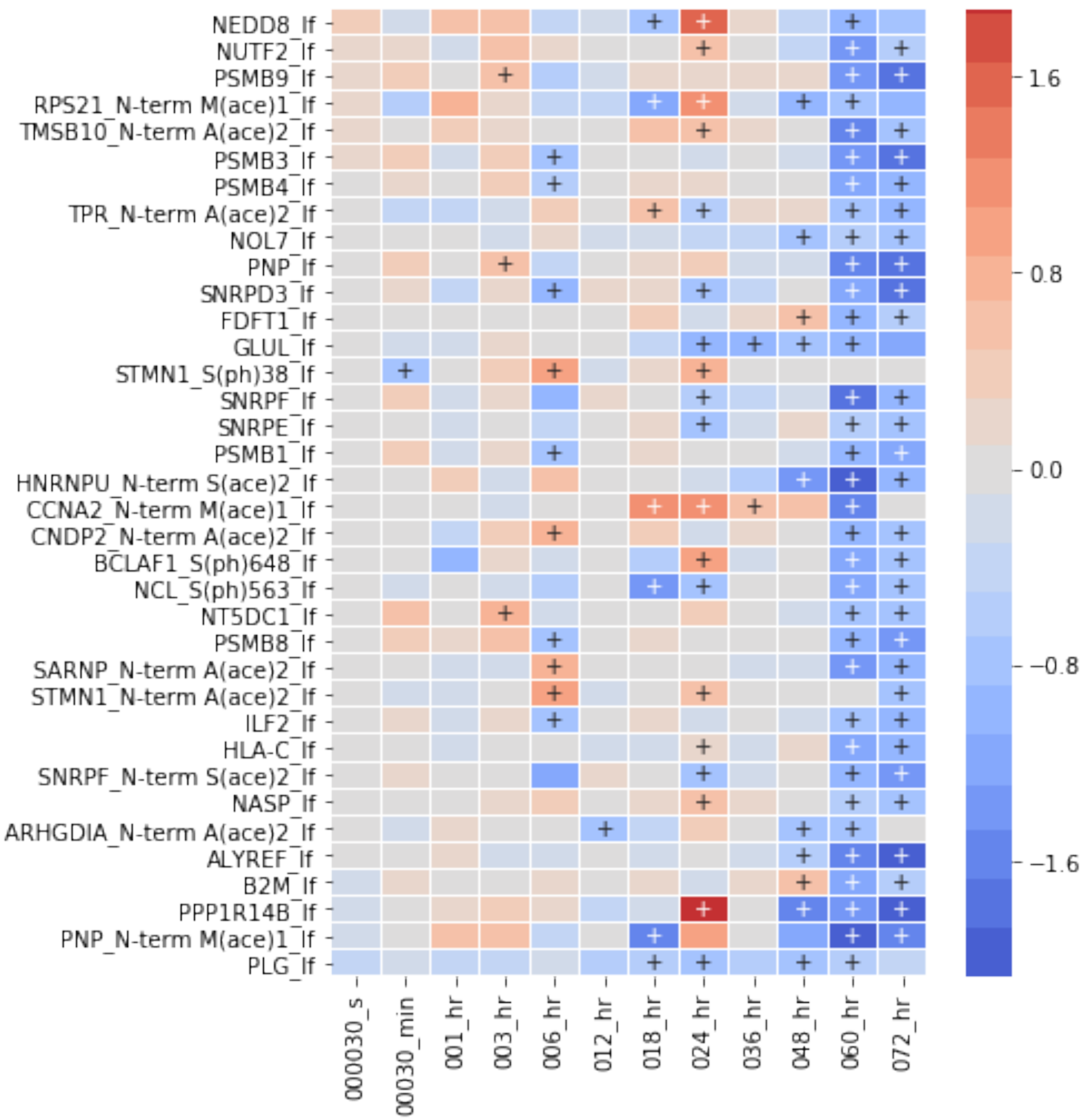
NEDD8_lf
NUTF2_lf
PSMB9_lf
RPS21_N-term M(ace)1_lf
TMSB10_N-term A(ace)2_lf
PSMB3_lf
PSMB4_lf
TPR_N-term A(ace)2_lf
NOL7_lf
PNP_lf
SNRPD3_lf
FDFT1_lf
GLUL_lf
STMN1_S(ph)38_lf
SNRPF_lf
SNRPE_lf
PSMB1_lf
HNRNPU_N-term S(ace)2_lf
CCNA2_N-term M(ace)1_lf
CNDP2_N-term A(ace)2_lf
BCLAF1_S(ph)648_lf
NCL_S(ph)563_lf
NT5DC1_lf
PSMB8_lf
SARNP_N-term A(ace)2_lf
STMN1_N-term A(ace)2_lf
ILF2_lf
HLA-C_lf
SNRPF_N-term S(ace)2_lf
NASP_lf
ARHGDIA_N-term A(ace)2_lf
ALYREF_lf
B2M_lf
PPP1R14B_lf
PNP_N-term M(ace)1_lf
PLG_lf
000030_s
00030_min
001_hr
003_hr
006_hr
012_hr
018_hr
024_hr
036_hr
048_hr
060_hr
072_hr
1.6
0.8
0.0
−0.8
−1.6

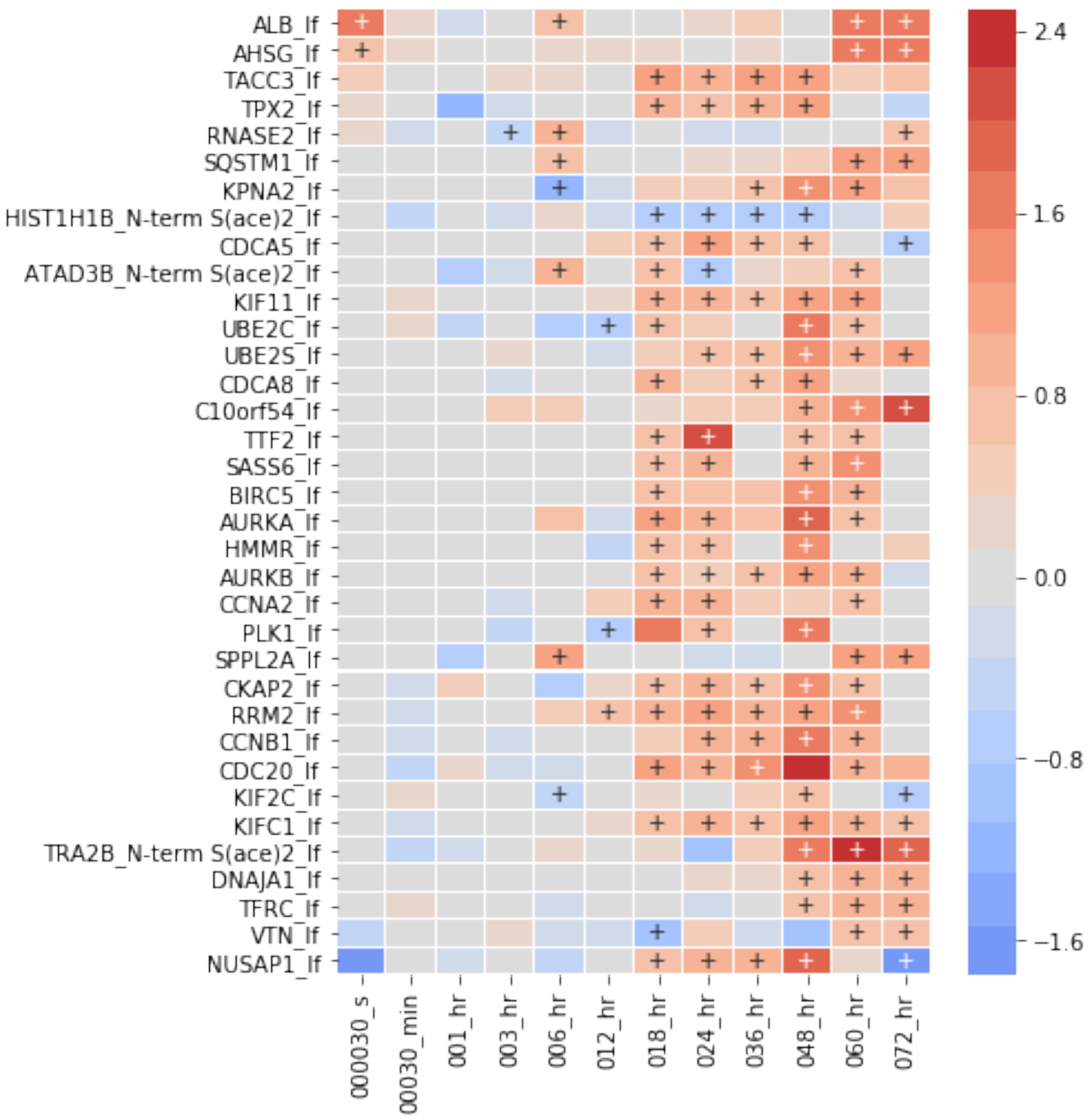


**Query 2:**

Changes that happen at all 2 timepoints for silac.

```
[34]: exp_data.silac.require_n_sig(
          n_sig=2, index='label'
      ).plot_species(plot_type='matplotlib');
```

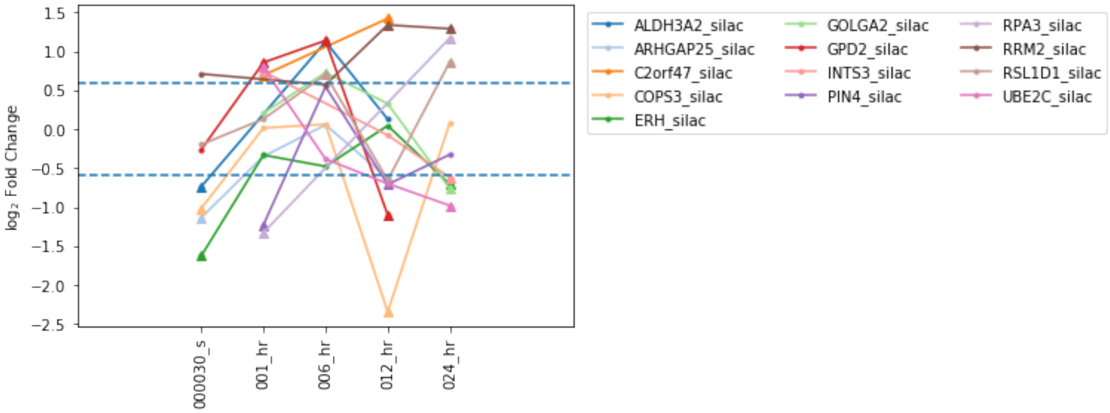


[35]:
```
int_species = 'CDK'
exp_data.label_free.heatmap(
    int_species,
    subset_index='identifier',
    index='label',
    min_sig=1,
    linewidths=0.01,
    figsize=(4, 3)
);
```

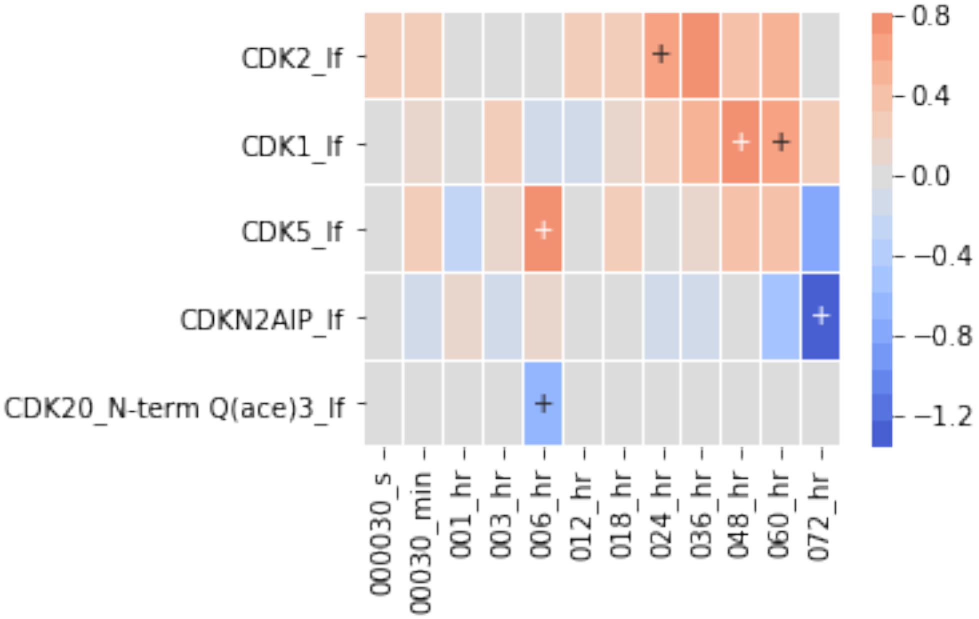

##### 1.2.9 Extending to other plots

Since our exp_data is built off a pandas.DataFrame, we can use other packages that take that data format. Seaborn is one such tool that provides some very nice plots.

[36]:
```
label_free = exp_data.label_free.copy()
label_free.log2_normalize_df(column='fold_change', inplace=True)

g = sns.PairGrid(label_free,
                 x_vars=('sample_id'),
                 y_vars=('fold_change', 'p_value'),
                 hue='source',
                 aspect=3.25, height=3.5)
g.map(
    sns.violinplot,
    palette="pastel",
    split=True,
    order=label_free.sample_ids
);
```

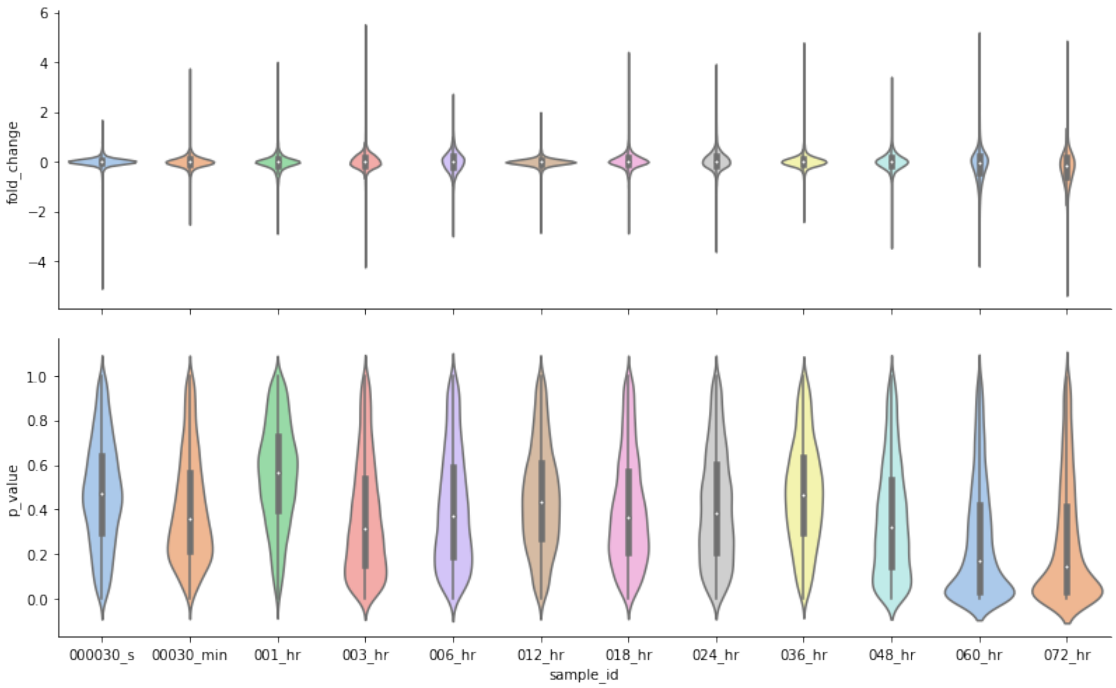


###### Visualizing signficant fraction of measured species

[37]:
```
fig = plt.figure(figsize=(5,5))
exp_data.label_free.plot_pie_sig_ratio('pie_label_free', fig.add_subplot(221))
plt.title("label_free");
exp_data.rna.plot_pie_sig_ratio('pie_rna_seq', fig.add_subplot(222))
```

```
plt.title("rna_seq");
exp_data.silac.plot_pie_sig_ratio('pie_silac', fig.add_subplot(223))
plt.title("silac");
exp_data.ph_silac.plot_pie_sig_ratio('pie_ph_silac', fig.add_subplot(224))
plt.title("ph_silac");
plt.savefig("pie_sig_omics.png", dpi=300, bbox_inches='tight')
```

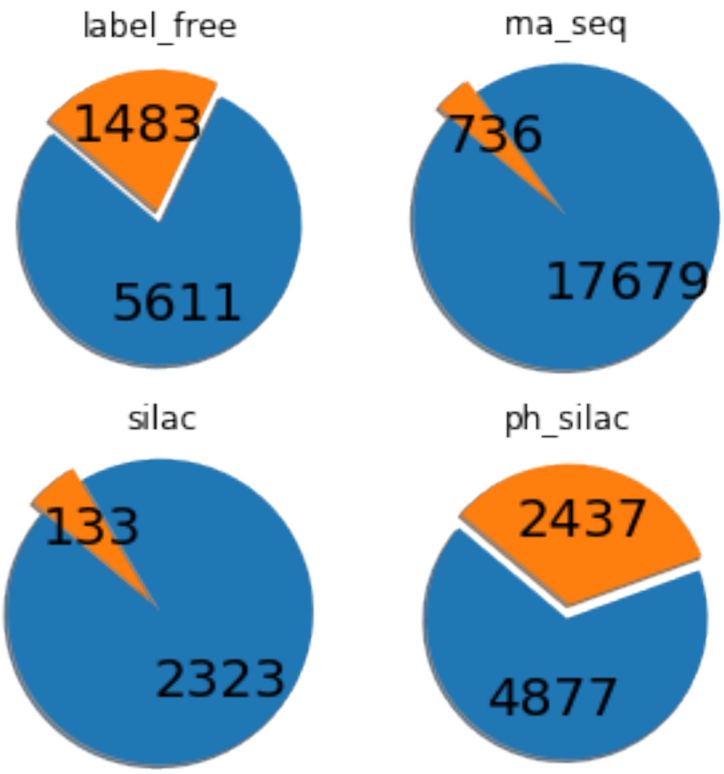


**Venn diagram comparisons between measurements**

[38]:
```
from magine.plotting.venn_diagram_maker import create_venn2, create_venn3

lf = exp_data.label_free.sig.id_list
silac = exp_data.silac.sig.id_list
phsilac = exp_data.ph_silac.sig.id_list
rna_names = exp_data.rna_seq.sig.id_list
hilic = exp_data.HILIC.sig.id_list
rplc = exp_data.C18.sig.id_list
fig = plt.figure(figsize=(8,6))

create_venn3(lf, silac, phsilac, 'LF', 'SILAC', 'ph-SILAC', ax=fig.
 ↪add_subplot(221));
plt.title("Compare proteomics");
```

```
create_venn2(set.union(*[lf, silac, phsilac]), rna_names,
             'Protein', 'RNA', ax=fig.add_subplot(223));
plt.title("Compare rna vs protein");
create_venn2(hilic, rplc, 'HILIC', 'RPLC', ax=fig.add_subplot(122));
plt.title("Compare metabolomics");
plt.tight_layout()
plt.savefig("venn_diagrams.png", dpi=300, bbox_inches='tight')
```

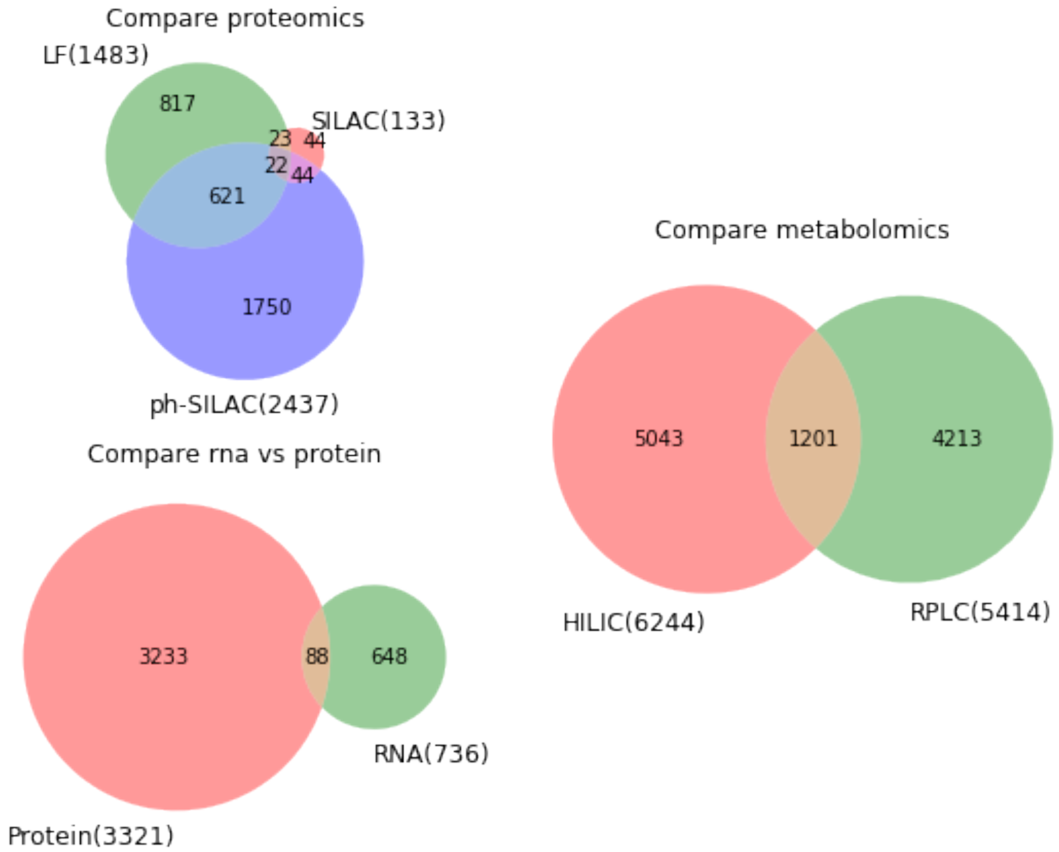


**Query 3:**

Extract out species that are significantly changed in all three omics (label-free, silac, ph-si

[39]:
```
all_three_omics = lf.intersection(silac).intersection(phsilac)
```

[40]:
```
exp_data.species.subset(all_three_omics).heatmap(
    all_three_omics,
    subset_index='identifier',
    columns=['source', 'sample_id',],
    min_sig=0,
    figsize=(8, 12), cluster_row=True,
```

```
    y_tick_labels=True, linewidths=0.01
);
```

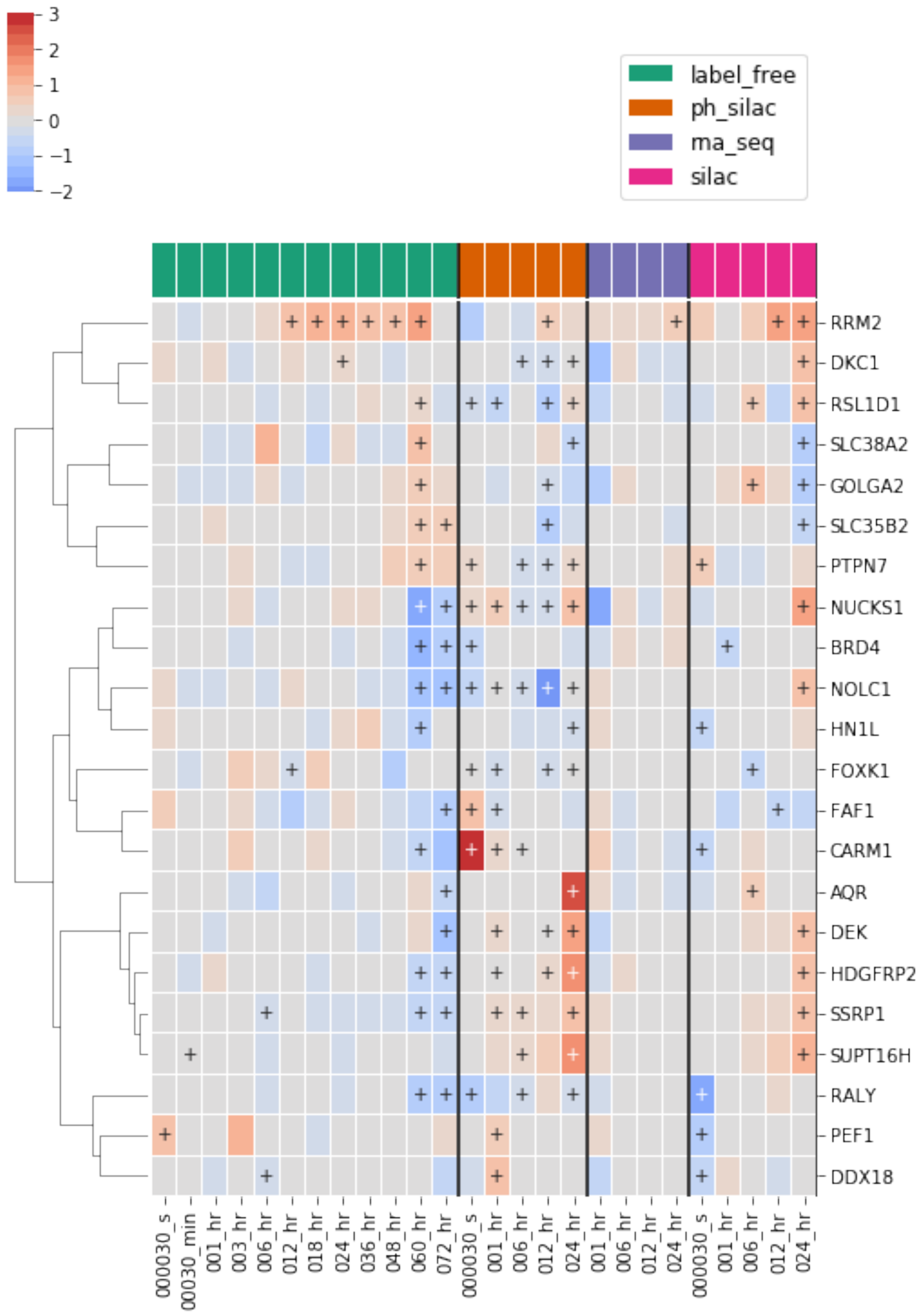

##### 1.2.10 Plot lists of interest

[41]:
```
# sample list for demo purposes
mrn_complex = ['NBN', 'RAD50', 'MRE11A']

exp_data.species.heatmap(
    mrn_complex,
    index='label',
    subset_index='identifier',
    min_sig=1,
    linewidths=0.01,
    figsize=(4,6)
);
```

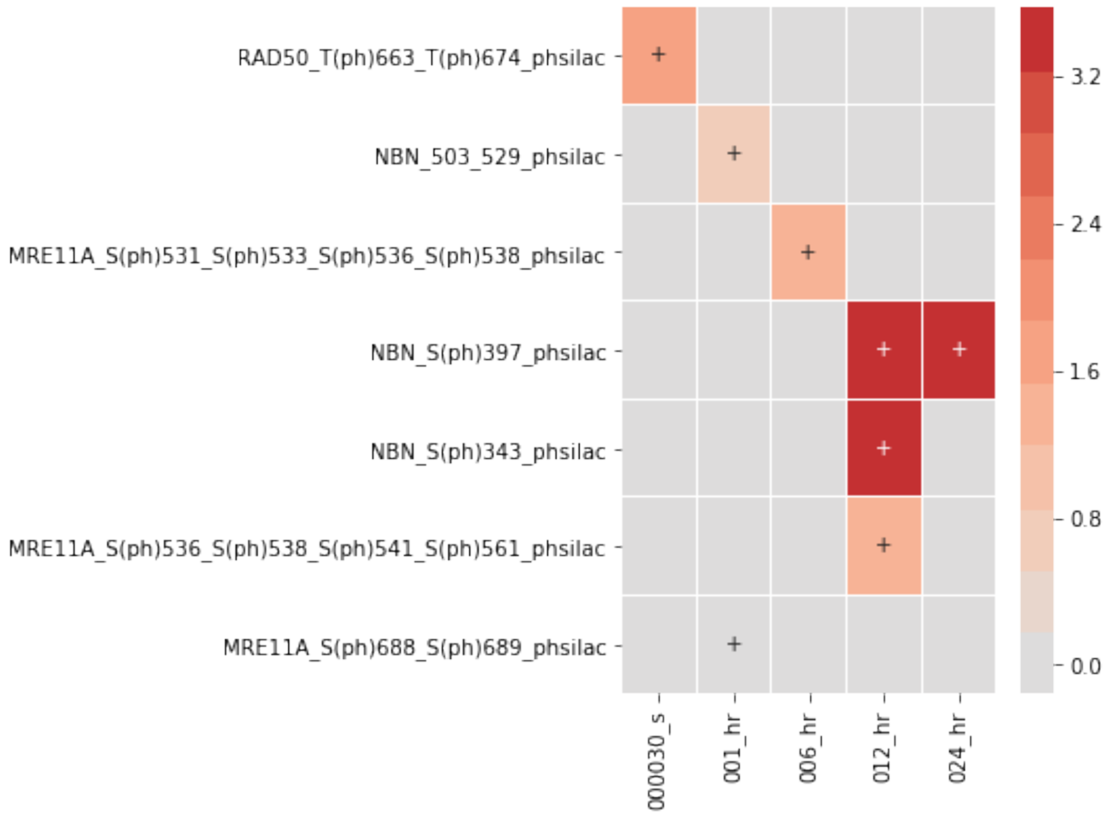


[42]:
```
# sample list for demo purposes
interesting_list = ['CHEK1', 'CHEK2']

exp_data.species.heatmap(
    interesting_list,
```

```
    index='label',
    subset_index='identifier',
    min_sig=1,
    linewidths=0.01,
    figsize=(4,6)
);
```

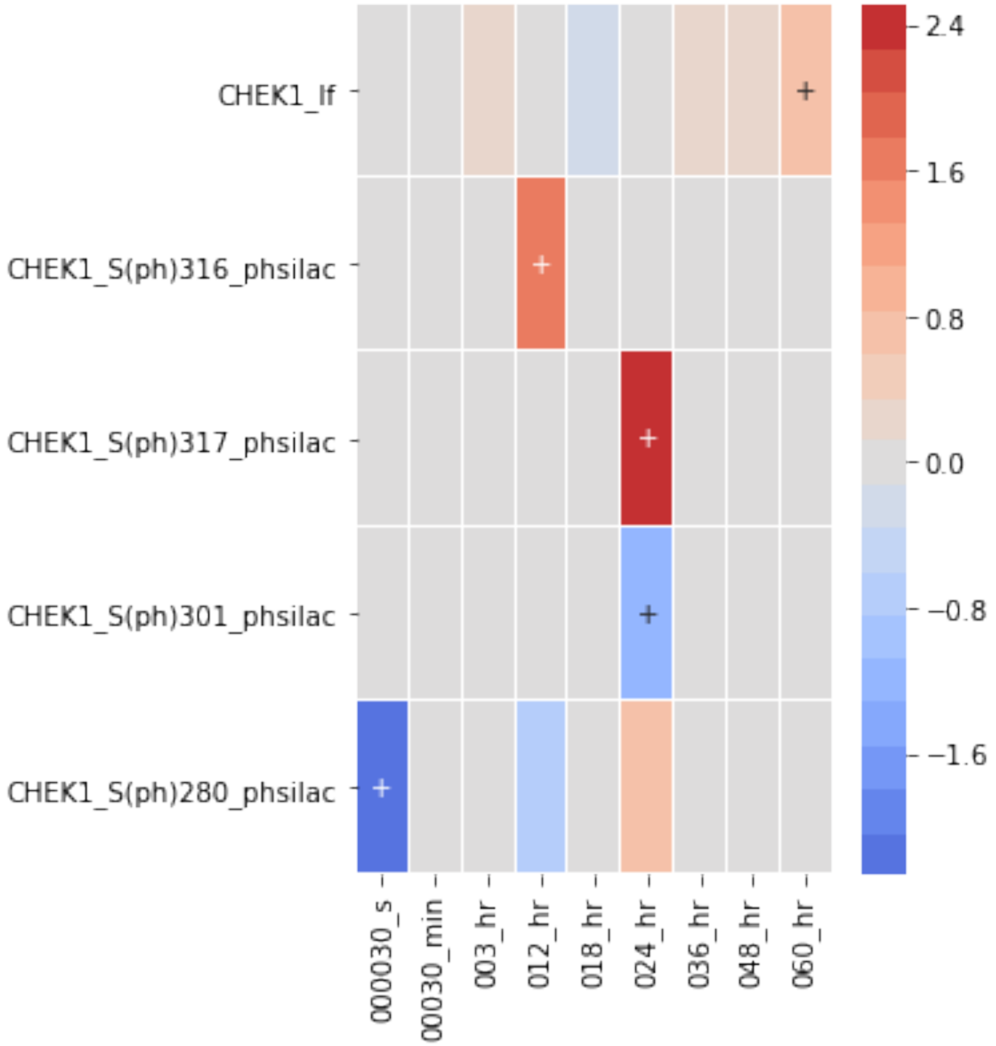


[43]:
```
# sample list for demo purposes
interesting_list = ['ATRIP', 'ERCC5']

exp_data.species.heatmap(
    interesting_list,
    index='label',
    subset_index='identifier',
    min_sig=1,
```

```
    linewidths=0.01,
    figsize=(4,6)
);
```

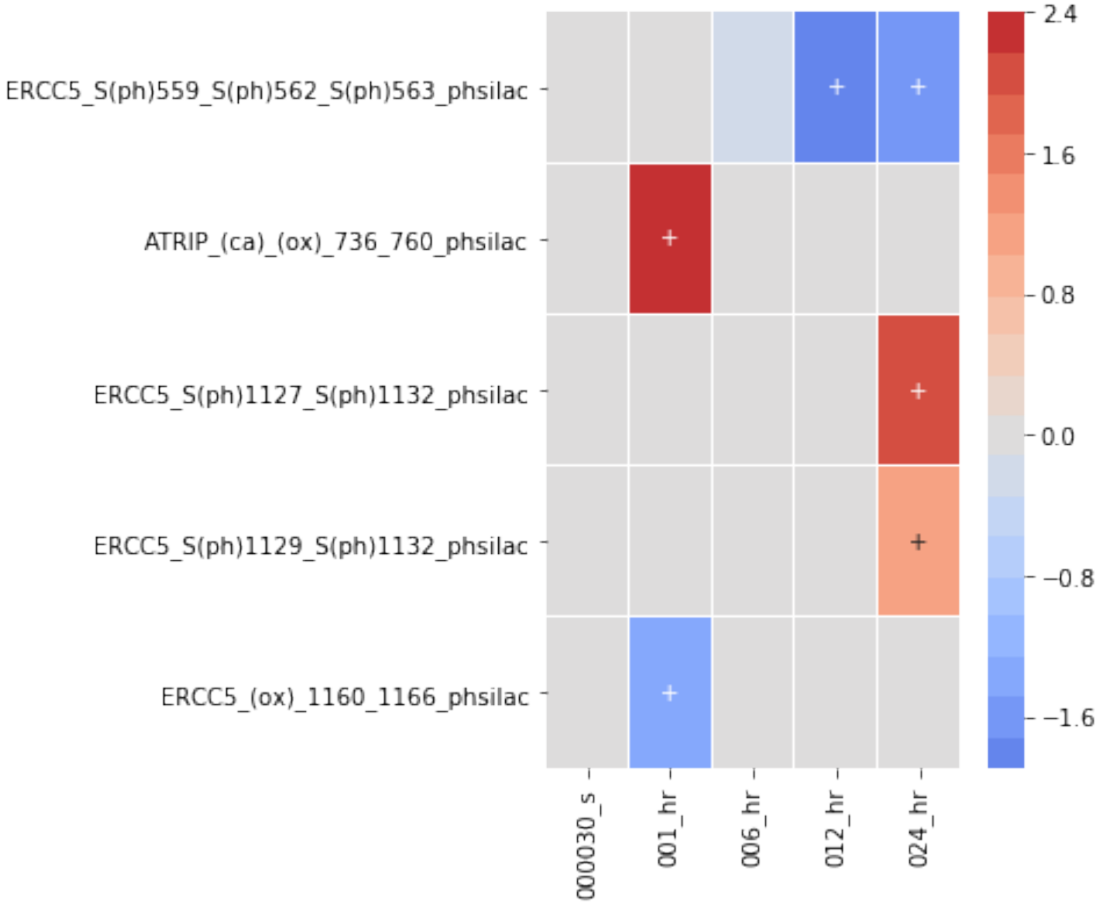

Supplement: S1-Jupyter Notebook [Data Exploration] [file 974121_file03.pdf]
