## Supplementary material for "A computational framework to explore cellular response mechanisms from multi-omics datasets": S2-Jupyter Notebook [Network Creation and Exploration]

### supplement_notebook_2_network_creation_and_exploration

December 10, 2019

#### 1 Creating a data driven network

This example shows how we create and add annotations to a data driven network.

[2]:
```
%matplotlib inline
import sys
import os
import networkx as nx
from IPython.display import display, Image
```

[3]:
```
# NBVAL_IGNORE_OUTPUT
from exp_data import exp_data
import magine.networks.utils as utils
from magine.networks.network_generator import build_network
from magine.networks.subgraphs import Subgraph
import magine.networks.exporters as exporters
import magine.networks.visualization as viz
```

```
2019-07-01 11:14:25.012 - magine - INFO - Logging started on MAGINE
2019-07-01 11:14:25.015 - magine - INFO - Log entry time offset from UTC: -7.00
hours
```

Creating list of seed species and background species for network

[4]:
```
measured = exp_data.species.id_list
sig_measured = exp_data.species.sig.id_list
print(len(measured))
print(len(sig_measured))
```

```
54750
14136
```

Now we will create the network. We pass the seed and background list to the network as well as flags turning on all of the network databases. We also trim source/sink nodes (optional). This basically cleans up dangling nodes that are not in our seed or background lists.

[5]:
```
save_name = 'bendamustine_network_w_attributes'

execute = False
```

```python
if execute:
    network = build_network(
        seed_species=sig_measured,  # seed species
        all_measured_list=measured,  # all data measured
        use_biogrid=True,  # expand with biogrid
        use_hmdb=True,  # expand with hmdb
        use_reactome=True,  # expand with reactome
        use_signor=True,  # expand with signor
        trim_source_sink=True,  # remove all source and sink nodes not measured
        save_name='Networks/bendamustine_network'
    )
    # add data to networks
    network = utils.add_data_to_graph(network, exp_data)

    # write to GML for cytoscape or other program
    nx.write_gml(network, os.path.join('Networks', save_name+'.gml'))

    # write to gpickle for fast loading in python
    nx.write_gpickle(network, os.path.join('Networks', save_name+'.p'))
else:
    network = nx.read_gpickle(os.path.join('Networks', save_name+'.p'))
```

[7]:
```python
print(len(network.nodes))
print(len(network.edges))
```

```
21130
515061
```

21130 nodes and 515061 edges are too much to manually explore. Thus, we are going to use the Subgraph Class to being to query the network.

[8]:
```python
# initialize it
net_sub = Subgraph(network)
```

[10]:
```python
help(net_sub)
```

```
Help on Subgraph in module magine.networks.subgraphs object:

class Subgraph(builtins.object)
 |  Subgraph(network, exp_data=None, pool=None)
 |
 |  Methods defined here:
 |
 |  __init__(self, network, exp_data=None, pool=None)
 |      Generates network subgraphs
 |
 |      Parameters
 |      ----------
```

```
 |      network : networkx.DiGraph
 |      exp_data : magine.data.datatypes.ExperimentalData
 |
 |  downstream_of_node(self, species_1, include_list=None, save_name=None,
draw=False)
 |      Generate network of all downstream species of provides species
 |
 |
 |      Parameters
 |      ----------
 |      species_1 : str
 |          species name
 |      save_name : str
 |          name to save gml file
 |      draw : bool
 |          create figure of graph
 |      include_list : list_like
 |          list of species that must be in path in order to consider a path
 |      Returns
 |      -------
 |      nx.DiGraph
 |
 |
 |      Examples
 |      --------
 |      >>> from networkx import DiGraph
 |      >>> from magine.networks.subgraphs import Subgraph
 |      >>> g = DiGraph()
 |      >>> g.add_edges_from([('a','b'),('b','c'), ('c', 'd'), ('a', 'd'),
('e', 'd')])
 |      >>> net_sub = Subgraph(g)
 |      >>> downstream_d = net_sub.downstream_of_node('d')
 |      >>> sorted(downstream_d.edges)
 |      []
 |      >>> downstream_c = net_sub.downstream_of_node('c')
 |      >>> sorted(downstream_c.edges)
 |      [('c', 'd')]
 |
 |  expand_neighbors(self, network=None, nodes=None, upstream=False,
downstream=False, max_dist=1, include_only=None,
add_interconnecting_edges=False)
 |      Create/expand a network based on neighbors from a list of species
 |
 |      Parameters
 |      ----------
 |      network : nx.DiGraph or None
 |          Starting network to expand nodes. If not provided, will use
 |          default network
```

```
 |      nodes : list_like
 |          List of nodes to expand
 |      upstream : bool
 |          Expand upstream nodes
 |      downstream : bool
 |          Expand downstream nodes
 |      max_dist :
 |          Max distance to explore
 |      include_only : list_like
 |          Limit network to only contain these species
 |      add_interconnecting_edges : bool
 |          Add edges connecting all nodes. Default if False, so only direct
 |          edges to neighbors will be added.
 |
 |      Returns
 |      -------
 |      nx.DiGraph
 |
 |  measured_networks_over_time(self, graph, colors, prefix)
 |      Adds color to a network over time
 |
 |      Parameters
 |      ----------
 |      graph : nx.DiGraph
 |      colors : list
 |          List of colors for time points
 |      prefix : str
 |          Prefix for image files
 |
 |
 |      Returns
 |      -------
 |
 |  measured_networks_over_time_up_down(self, graph, prefix, color_up='tomato',
color_down='lightblue')
 |      Parameters
 |      ----------
 |      graph : nx.DiGraph
 |
 |      prefix : str
 |          Prefix for image files
 |      color_up : str
 |
 |      color_down : str
 |
 |
 |      Returns
 |      -------
```

```
 |
 |  neighbors(self, node, upstream=True, downstream=True, max_dist=1,
include_only=None, start_network=None)
 |      Create network containing provided node and its neighbors.
 |
 |      Parameters
 |      ----------
 |      node : str
 |      upstream : bool
 |      downstream : bool
 |      max_dist : int
 |      include_only : list
 |      start_network : nx.DiGraph
 |
 |
 |      Returns
 |      -------
 |      nx.DiGraph
 |
 |  paths_between_list(self, species_list, single_path=False, max_length=None,
add_interconnecting_edges=False, include_only=None, pool=None, save_name=None,
draw=False, image_format='png')
 |      Returns graph containing all shortest paths between list.
 |
 |      Parameters
 |      ----------
 |
 |      species_list : list_like
 |          list of species
 |      save_name : str
 |          name to save
 |      single_path : bool
 |          use single shortest path if True, else use all shortest paths
 |      draw : bool
 |          create a dot generated figure
 |      image_format : str
 |          dot acceptable output formats, (pdf, png, etc)
 |      pool : multiprocessing.Pool
 |          If it it provided, it uses its map function to run this function.
 |      max_length : int
 |          Max length for path between any 2 species
 |      include_only : list_like
 |          List of species that must be present
 |      Returns
 |      -------
 |      graph : networkx.DiGraph
 |          graph containing paths between species list provided
 |
```

```
 |
 |      Examples
 |      --------
 |      >>> from networkx import DiGraph
 |      >>> from magine.networks.subgraphs import Subgraph
 |      >>> g = DiGraph()
 |      >>> g.add_edges_from([('a','b'),('b','c'), ('c', 'd'), ('a', 'd'), ('e',
'd')])
 |      >>> g.add_path(['g', 'h', 'c', 'i', 'j', 'k'])
 |      >>> net_sub = Subgraph(g)
 |      >>> path_a_d = net_sub.paths_between_list(['a','c','d'])
 |      >>> sorted(path_a_d.edges)
 |      [('a', 'b'), ('a', 'd'), ('b', 'c'), ('c', 'd')]
 |      >>> path_a_f = net_sub.paths_between_list(['g', 'h', 'j'], max_length=4)
 |      >>> sorted(path_a_f.edges)
 |      [('c', 'i'), ('g', 'h'), ('h', 'c'), ('i', 'j')]
 |
 |  paths_between_pair(self, node_1, node_2, bidirectional=False,
single_path=False, draw=False, image_format='png')
 |      Generates a graph based on all shortest paths between two species.
 |
 |
 |      Parameters
 |      ----------
 |      node_1 : str
 |          name of first species
 |      node_2 : str
 |          name of second species
 |      bidirectional : bool
 |          If you want to search bidirectionally
 |      single_path : bool
 |          If you only want a single shortest path
 |      draw : bool
 |          create an image of returned network
 |      image_format : str, optional
 |          If draw=True you can pass an image format. (pdf, png, svg).
 |          default=png
 |
 |      Returns
 |      -------
 |      graph : networkx.DiGraph
 |
 |
 |      Examples
 |      --------
 |      >>> from networkx import DiGraph
 |      >>> from magine.networks.subgraphs import Subgraph
 |      >>> g = DiGraph()
```

```
 |      >>> g.add_edges_from([('a','b'),('b','c'), ('c', 'd'),  ('e', 'd'),
('d', 'a')])
 |      >>> net_sub = Subgraph(g)
 |      >>> path_a_d = net_sub.paths_between_pair('a','d', False)
 |      >>> sorted(path_a_d.edges)
 |      [('a', 'b'), ('b', 'c'), ('c', 'd')]
 |      >>> path_a_d = net_sub.paths_between_pair('a','d', True)
 |      >>> sorted(path_a_d.edges)
 |      [('a', 'b'), ('b', 'c'), ('c', 'd'), ('d', 'a')]
 |
 |  paths_between_two_lists(self, list_1, list_2, single_path=False,
max_length=None, include_only=None, reverse=False,
add_interconnecting_edges=False, draw=False, save_name=None, image_format='png')
 |      Generates a graph based on all shortest paths between two species.
 |
 |
 |      Parameters
 |      ----------
 |      list_1 : list
 |          Node names
 |      list_2 : list
 |          Node names
 |      single_path : bool
 |          If you only want a single shortest path.
 |      max_length : int
 |          Maximum distance between any two species.
 |      include_only : list
 |          Species required to be in paths/
 |      reverse : bool
 |          Flag to check list_2 to list_1. Default will only look for list_1
 |          to list_2.
 |      add_interconnecting_edges : bool
 |          Add edges between species even if not between list_1 and list_2
 |          nodes.
 |      save_name : str
 |          Save of figure/network
 |      draw : bool
 |          create an image of returned network
 |      image_format : str, optional
 |          If draw=True you can pass an image format. (pdf, png, svg).
 |          default=png
 |
 |      Returns
 |      -------
 |      graph : networkx.DiGraph
 |
 |      Examples
 |      --------
```

```
 |      >>> from networkx import DiGraph
 |      >>> from magine.networks.subgraphs import Subgraph
 |      >>> g = DiGraph()
 |      >>> g.add_path(['a', 'b', 'c', 'd'])
 |      >>> g.add_path(['g', 'h', 'c', 'i'])
 |      >>> net_sub = Subgraph(g)
 |      >>> path_a_d = net_sub.paths_between_two_lists(['a','g'], ['c', 'i'],
max_length=3)
 |      >>> sorted(path_a_d.edges)
 |      [('a', 'b'), ('b', 'c'), ('g', 'h'), ('h', 'c')]
 |
 |  upstream_of_node(self, species_1, include_list=None, save_name=None,
draw=False)
 |      Generate network of all upstream species of provides species
 |
 |      Parameters
 |      ----------
 |      species_1 : str
 |          species name
 |      save_name : str
 |          name to save gml file
 |      draw : bool
 |          create figure of graph
 |      include_list : list_like
 |          Species that must be in path in order to consider a path
 |
 |      Returns
 |      -------
 |      nx.DiGraph
 |
 |
 |      Examples
 |      --------
 |      >>> from networkx import DiGraph
 |      >>> from magine.networks.subgraphs import Subgraph
 |      >>> g = DiGraph()
 |      >>> g.add_edges_from([('a','b'),('b','c'), ('c', 'd'), ('a', 'd'),
('e', 'd')])
 |      >>> net_sub = Subgraph(g)
 |      >>> upstream_d = net_sub.upstream_of_node('d')
 |      >>> sorted(upstream_d.edges())
 |      [('a', 'd'), ('b', 'c'), ('c', 'd'), ('e', 'd')]
 |      >>> upstream_c = net_sub.upstream_of_node('c')
 |      >>> sorted(upstream_c.edges())
 |      [('a', 'b'), ('b', 'c')]
 |
 |  ----------------------------------------------------------------------
 |  Data descriptors defined here:
```

```
 |
 |  __dict__
 |      dictionary for instance variables (if defined)
 |
 |  __weakref__
 |      list of weak references to the object (if defined)
```

#### 1.1 Exploring neighbors of nodes of interest

For demonstration purposes, we are starting with the protein CASP3. CASP3 is an effector caspase that is required for apoptosis. It cleaves other proteins and starts the degradation CASP3. We start our exploration at CASP3 as it is a marker for intrinsic apoptosis, which is what is generally regarded as bendamustines pathway for cell death.

First, we are going to look at all the neighbors of CASP3 that were signficantly changed in our experimental data.

[23]:
```
casp3_neighbors = net_sub.neighbors(
    'CASP3', # node of interest
    upstream=True, # include upstream nodes
    downstream=False,  # include downstream nodes
    include_only=exp_data.species.sig.id_list # limit nodes to only significant␣
 ↪changed species
)
```

[24]:
```
# one of many ways to draw the network.
# draw_cyjs is ideal for Jupyter notebooks since it allows us to move nodes,␣
 ↪apply various layouts
# Notice that if you click on an edge, it provides you with the interaction␣
 ↪type.
# If you click on a node, it provides a link to genecards.
viz.draw_cyjs(casp3_neighbors)
```

```
<IPython.core.display.HTML object>
```

Next we can continue to expand this network to explore nodes of interest.

This next function expands a single node and creates plots of the species that are connected to thht node.

[61]:
```
def show_neighbors(node, df, upstream=True, downstream=False, max_dist=1,
                   include_only=None, figsize=None, show_network=False):

    neighbors = net_sub.expand_neighbors(
        network=None,
        nodes=node,
        upstream=upstream,
        downstream=downstream,
        max_dist=max_dist,
        include_only=include_only
```

```
    )
    df_copy = df.subset(neighbors.nodes).copy()

    # remove nodes not connected to casp3
    neighbors = utils.delete_disconnected_network(neighbors)

    # moves a time point if no signficant changes
    df_copy.require_n_sig(n_sig=1, inplace=True, index='sample_id',␣
↪columns='label',)

    # removes measured species if no signficant changes
    df_copy.require_n_sig(n_sig=1, index='label', inplace=True)
    if show_network:
        # export image
        s_name =  'node_{}.png'.format(node)
        exporters.export_to_dot(neighbors, s_name, image_format='png',␣
↪engine='circo')

        # display image
        display(Image(s_name, width=400))

    # create heatmap of the neighnor nodes
    fig = df_copy.heatmap(
        rank_index=True,
        index='label',
        linewidths=0.01,
        figsize=figsize
    );


show_neighbors('CYCS',
               exp_data.species,
               upstream=True,
               downstream=False,
               max_dist=1,
               include_only=exp_data.species.sig.id_list
)
```

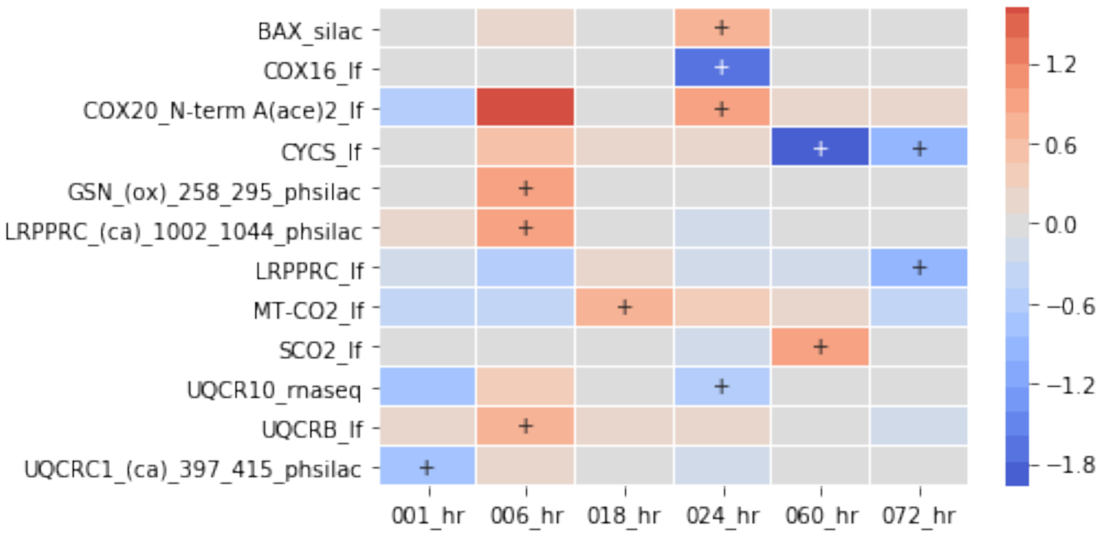


[62]:
```
expand = net_sub.expand_neighbors(casp3_neighbors,
                                  nodes='CYCS',
                                  upstream=True,
                                  include_only=exp_data.species.sig.id_list)
```

[63]:
```
viz.draw_cyjs(expand)
```

<IPython.core.display.HTML object>

[64]:
```
expand = net_sub.expand_neighbors(expand, nodes='BAX', upstream=True,␣
 ↪include_only=exp_data.species.sig.id_list)
viz.draw_cyjs(expand)
```

<IPython.core.display.HTML object>

[65]:
```
expand = net_sub.expand_neighbors(expand, nodes='BID', upstream=True,␣
 ↪include_only=exp_data.species.sig.id_list)
viz.draw_cyjs(expand)
```

<IPython.core.display.HTML object>

[67]:
```
show_neighbors('CASP8',
               exp_data.species,
               upstream=True,
               downstream=False,
               max_dist=1,
```

```
                  include_only=exp_data.species.sig.id_list,
                  figsize=(6, 12)
)
```

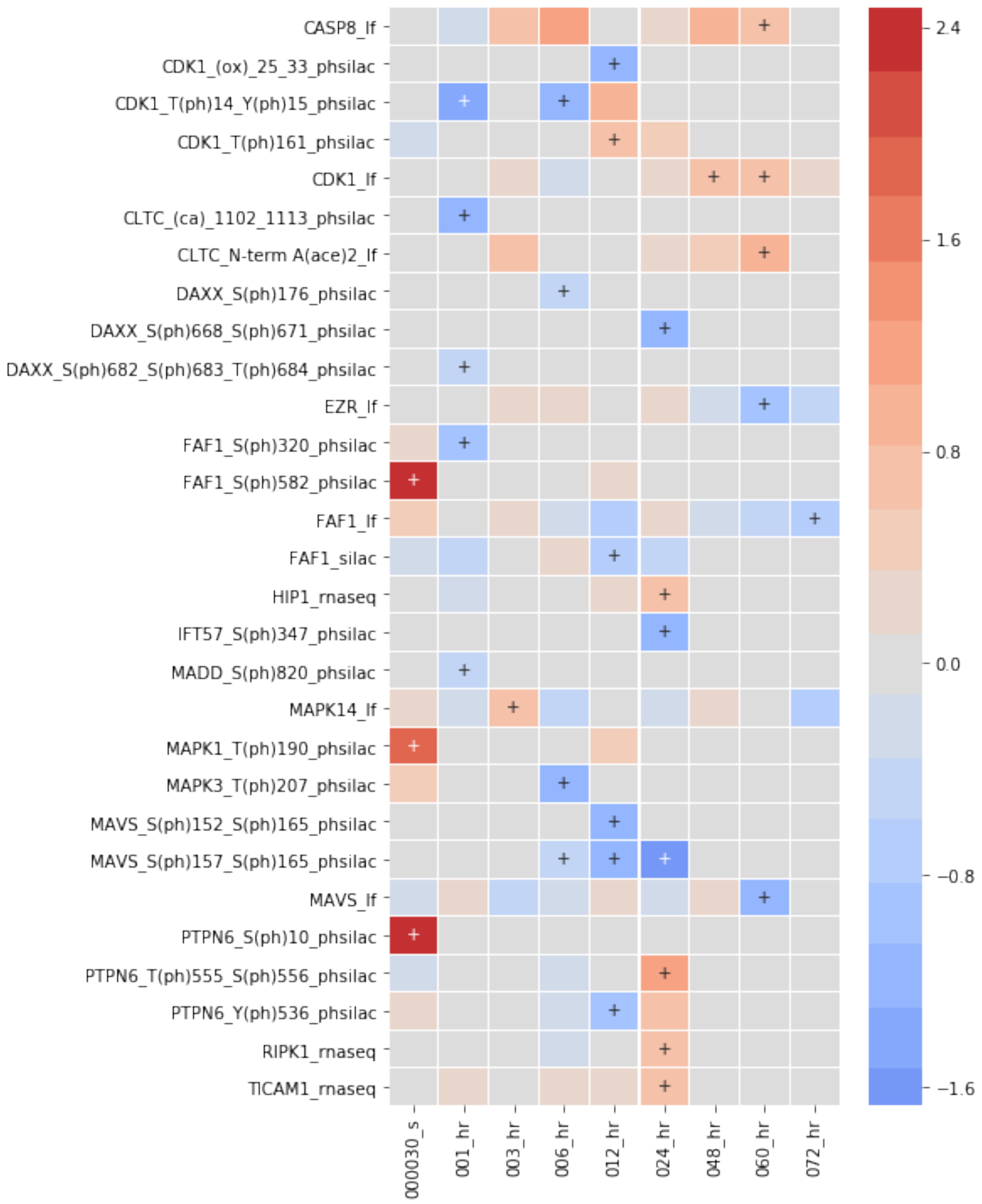


[68]:
```
expand = net_sub.expand_neighbors(expand, nodes='CASP8', upstream=True,␣
↪include_only=exp_data.species.sig.id_list)
```

```
viz.draw_cyjs(expand)
```

<IPython.core.display.HTML object>

[ ]:
```
show_neighbors(['FAF1', 'MADD', 'DAXX'],
               exp_data.species,
               upstream=True,
               downstream=False,
               max_dist=1,
               include_only=exp_data.species.require_n_sig(n_sig=1).id_list,
               figsize=(8, 6)
)
```

[79]:
```
expand = net_sub.expand_neighbors(expand,
                                  nodes=['FAF1', 'MADD', 'DAXX'],
                                  upstream=True,
                                  include_only=exp_data.species.sig.id_list)
# NBVAL_IGNORE_OUTPUT
viz.draw_cyjs(expand)
```

<IPython.core.display.HTML object>

[70]:
```
show_neighbors('TNFRSF1B',
               exp_data.species,
               upstream=True,
               downstream=False,
               max_dist=1,
               include_only=exp_data.species.require_n_sig(n_sig=1).id_list,
               figsize=(8, 6)
)
```

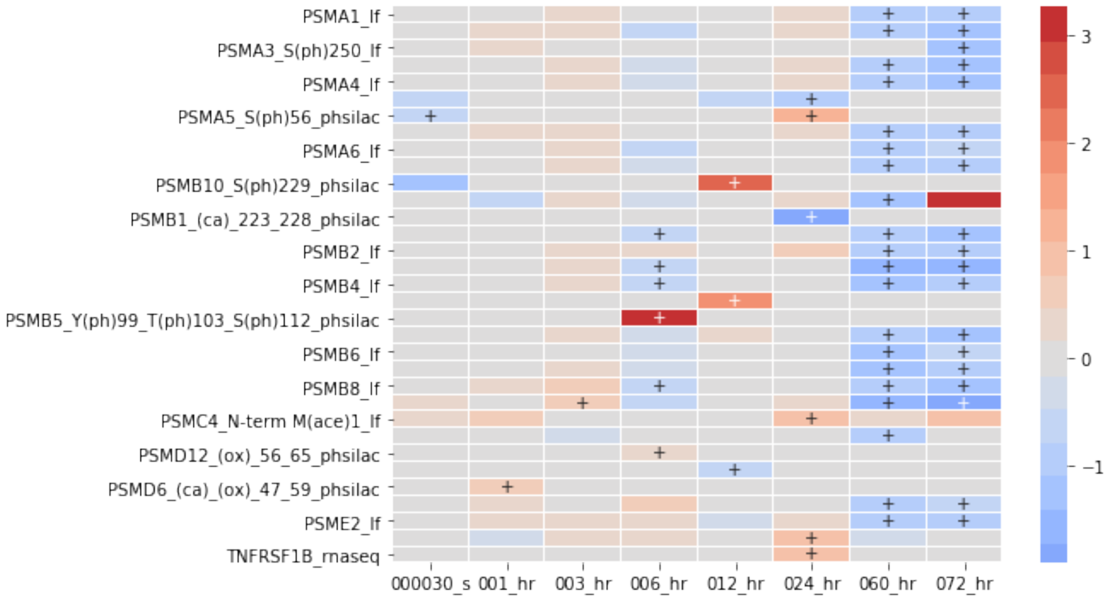


[71]:
```
show_neighbors('AURKA',
               exp_data.species,
               upstream=True,
               downstream=False,
               max_dist=1,
               include_only=exp_data.species.require_n_sig(n_sig=1).id_list,
               figsize=(8, 12),
               show_network=True
)
```

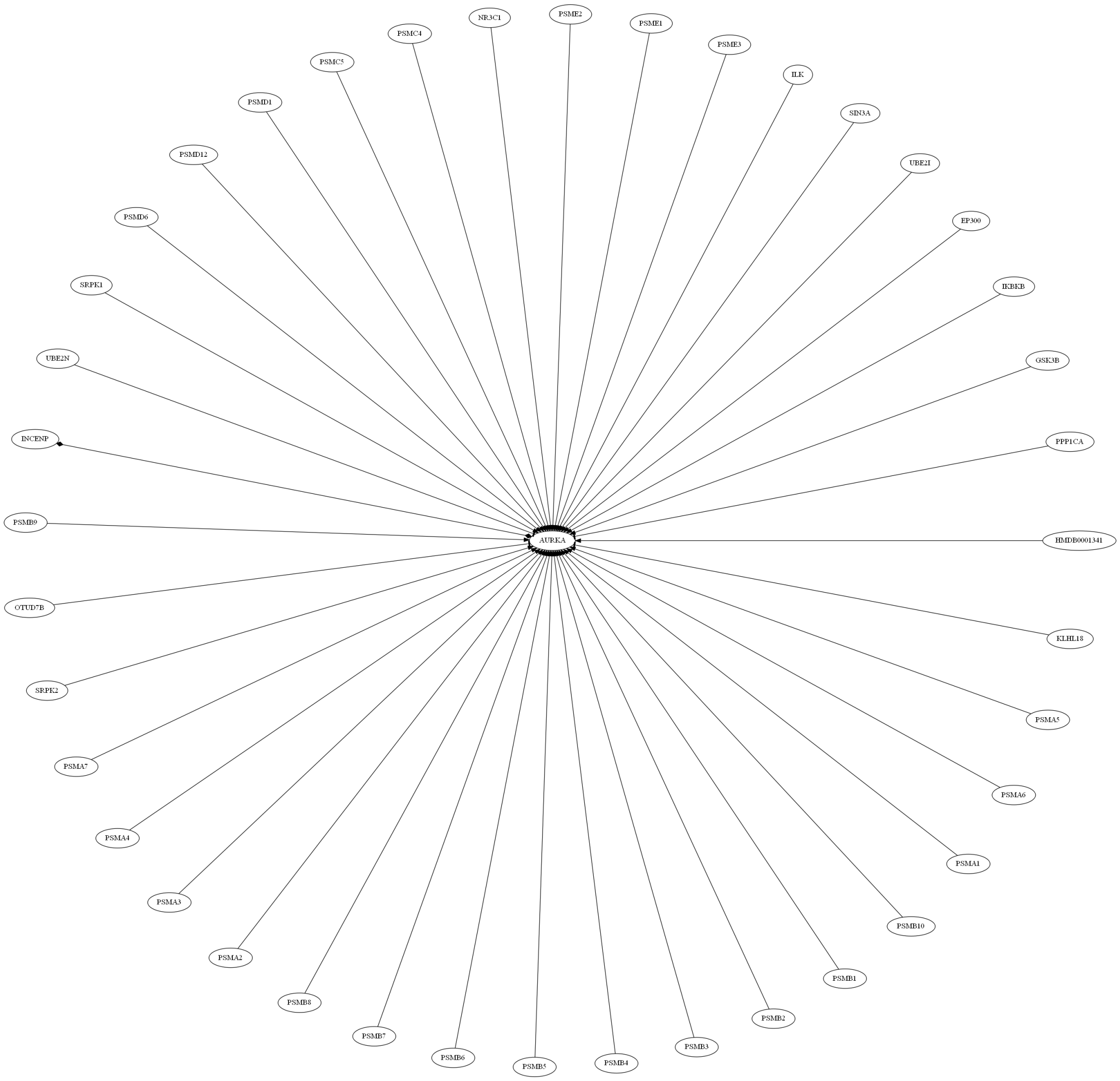
NR3C1
PSME2
PSMC4
PSME1
PSMC5
PSME3
ILK
PSMD1
SIN3A
PSMD12
UBE2I
PSMD6
EP300
SRPK1
IKBKB
UBE2N
GSK3B
INCENP
PPP1CA
PSMB9
AURKA
HMDB0001341
OTUD7B
KLHL18
SRPK2
PSMA5
PSMA7
PSMA6
PSMA4
PSMA1
PSMA3
PSMB10
PSMA2
PSMB1
PSMB8
PSMB2
PSMB7
PSMB3
PSMB6
PSMB4
PSMB5

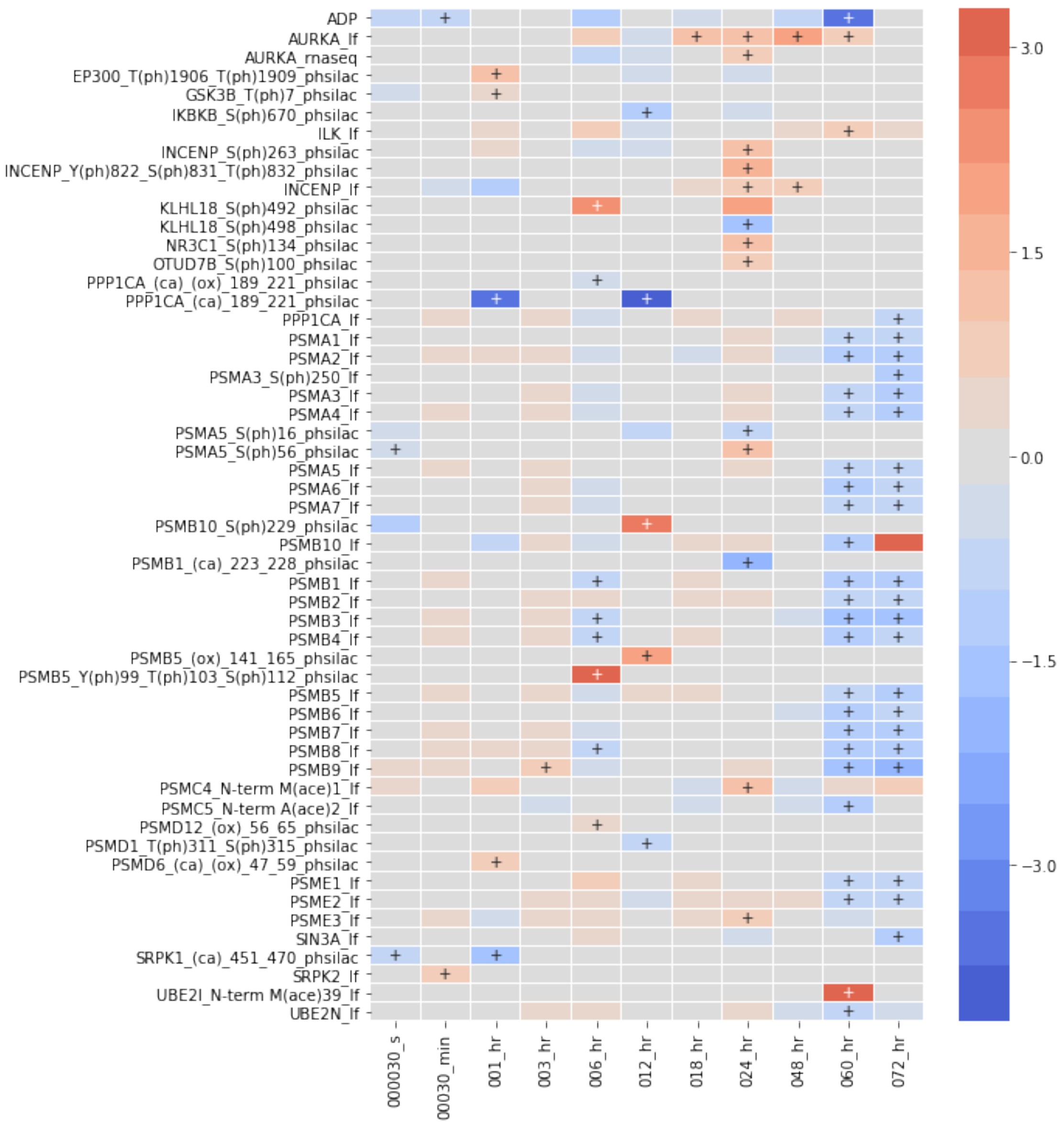


[83]:
```
expand = net_sub.expand_neighbors(expand, nodes=['AURKA'], upstream=True,␣
 ↪include_only=exp_data.species.require_n_sig(n_sig=2).id_list)
# NBVAL_IGNORE_OUTPUT
viz.draw_cyjs(expand)
```

```
<IPython.core.display.HTML object>
```

This last network has gotten a little hard to explore. We can simplify it by just looking for a single path, in this case AURKA to CYCS

[87]:
```
sub_g_2 = Subgraph(expand)
aurka_to_cycs = sub_g_2.paths_between_pair('AURKA', 'CYCS')
```

[88]:
```
viz.draw_cyjs(aurka_to_cycs)
```

```
<IPython.core.display.HTML object>
```

[ ]:
```
```
