## Supplementary material for "A computational framework to explore cellular response mechanisms from multi-omics datasets": S3-Jupyter Notebook [Enrichment Analysis]

### supplement_notebook_3_enrichment_analysis

December 10, 2019

[1]:
```
from IPython.display import display
%matplotlib inline

import pandas as pd
import numpy as np
import networkx as nx
import matplotlib.pyplot as plt
import seaborn as sns
```

[2]:
```
# load the experimental data
from exp_data import exp_data
```

##### 0.0.1 Running enrichment analysis via EnrichR

MAGINE allows users to upload lists of genes for analysis and retrieves the results in an `EnrichmentResult` Class.

[ ]:
```
from magine.enrichment.enrichr import Enrichr
e = Enrichr()
```

[8]:
```
help(e)
```

```
Help on Enrichr in module magine.enrichment.enrichr object:

class Enrichr(builtins.object)
 |  Enrichr(verbose=False)
 |
 |  Methods defined here:
 |
 |  __init__(self, verbose=False)
 |      Initialize self.  See help(type(self)) for accurate signature.
 |
 |  print_valid_libs(self)
 |      Print a list of all available libraries EnrichR has to offer.
 |
 |  run(self, list_of_genes, gene_set_lib='GO_Biological_Process_2017')
 |      Parameters
 |      ----------
```

```
 |      list_of_genes : list_like
 |          List of genes using HGNC gene names
 |      gene_set_lib : str or list
 |          Name of gene set library
 |          To print options use Enrichr.print_valid_libs
 |
 |
 |      Examples
 |      --------
 |      >>> import pandas as pd
 |      >>> pd.set_option('display.max_colwidth', 40)
 |      >>> pd.set_option('precision', 3)
 |      >>> e = Enrichr()
 |      >>> df = e.run(['BAX', 'BCL2', 'CASP3', 'CASP8'],
gene_set_lib='Reactome_2016')
 |      >>> print(df[['term_name','combined_score']].head(5))#doctest:
+NORMALIZE_WHITESPACE
 |                                        term_name  combined_score
 |          0  intrinsic pathway for apoptosis_hsa_...      48.157
 |          1  programmed cell death_hsa_r-hsa-5357801      41.516
 |          2              apoptosis_hsa_r-hsa-109581      41.403
 |          3  caspase-mediated cleavage of cytoske...      27.349
 |          4  caspase activation via extrinsic apo...      22.438
 |
 |      Returns
 |      -------
 |      df : EnrichmentResult
 |          Results from enrichR
 |
 |  run_samples(self, sample_lists, sample_ids,
gene_set_lib='GO_Biological_Process_2017', save_name=None, create_html=False,
out_dir=None, run_parallel=False, exp_data=None, pivot=False)
 |      Run enrichment analysis on a list of samples.
 |
 |      Parameters
 |      ----------
 |      sample_lists : list_like
 |          List of lists of genes for enrichment analysis
 |      sample_ids : list
 |          list of ids for the provided sample list
 |      gene_set_lib : str, list
 |          Type of gene set, refer to Enrichr.print_valid_libs
 |      save_name : str, optional
 |          if provided it will save a file as a pivoted table with
 |          the term_ids vs sample_ids
 |      create_html : bool
 |          Creates html of output with plots of species across sample
 |      out_dir : str
```

```
 |          If create_html, it will place all html plots into this directory
 |      run_parallel : bool
 |          If create_html, it will create plots using multiprocessing
 |      exp_data : magine.data.ExperimentalData
 |          Must be provided if create_html=True
 |      pivot : bool
 |
 |      Examples
 |      --------
 |      .. plot::
 |          :context: close-figs
 |
 |          >>> import pandas as pd
 |          >>> import matplotlib.pyplot as plt
 |          >>> from magine.enrichment.enrichr import Enrichr
 |          >>> pd.set_option('display.max_colwidth', 40)
 |          >>> pd.set_option('precision', 3)
 |          >>> samples = [['BAX', 'BCL2', 'CASP3', 'CASP8'], ['ATR', 'ATM',
'TP53', 'CHEK1']]
 |          >>> sample_ids = ['apoptosis', 'dna_repair']
 |          >>> e = Enrichr()
 |          >>> df = e.run_samples(samples, sample_ids,
gene_set_lib='Reactome_2016')
 |          >>> print(df[['term_name','combined_score']].head(5))#doctest:
+NORMALIZE_WHITESPACE
 |                                            term_name  combined_score
 |              0  intrinsic pathway for apoptosis_hsa_...      48.157
 |              1  programmed cell death_hsa_r-hsa-5357801      41.516
 |              2              apoptosis_hsa_r-hsa-109581      41.403
 |              3  caspase-mediated cleavage of cytoske...      27.349
 |              4  caspase activation via extrinsic apo...      22.438
 |          >>> df.filter_multi(rank=10, inplace=True)
 |          >>> df['term_name'] = df['term_name'].str.split('_').str.get(0)
 |          >>> fig = df.sig.heatmap(figsize=(6, 6), linewidths=.05)
 |
 |      Returns
 |      -------
 |      EnrichmentResult
 |
 |  ----------------------------------------------------------------------
 |  Data descriptors defined here:
 |
 |  __dict__
 |      dictionary for instance variables (if defined)
 |
 |  __weakref__
 |      list of weak references to the object (if defined)
```

[5]:
```
# from supplement_notebook_1
exp_data.label_free.heatmap(
    index='label',
    linewidths=0.01,
    cluster_row=True,
    min_sig=4,
    figsize=(4, 8)
);
```

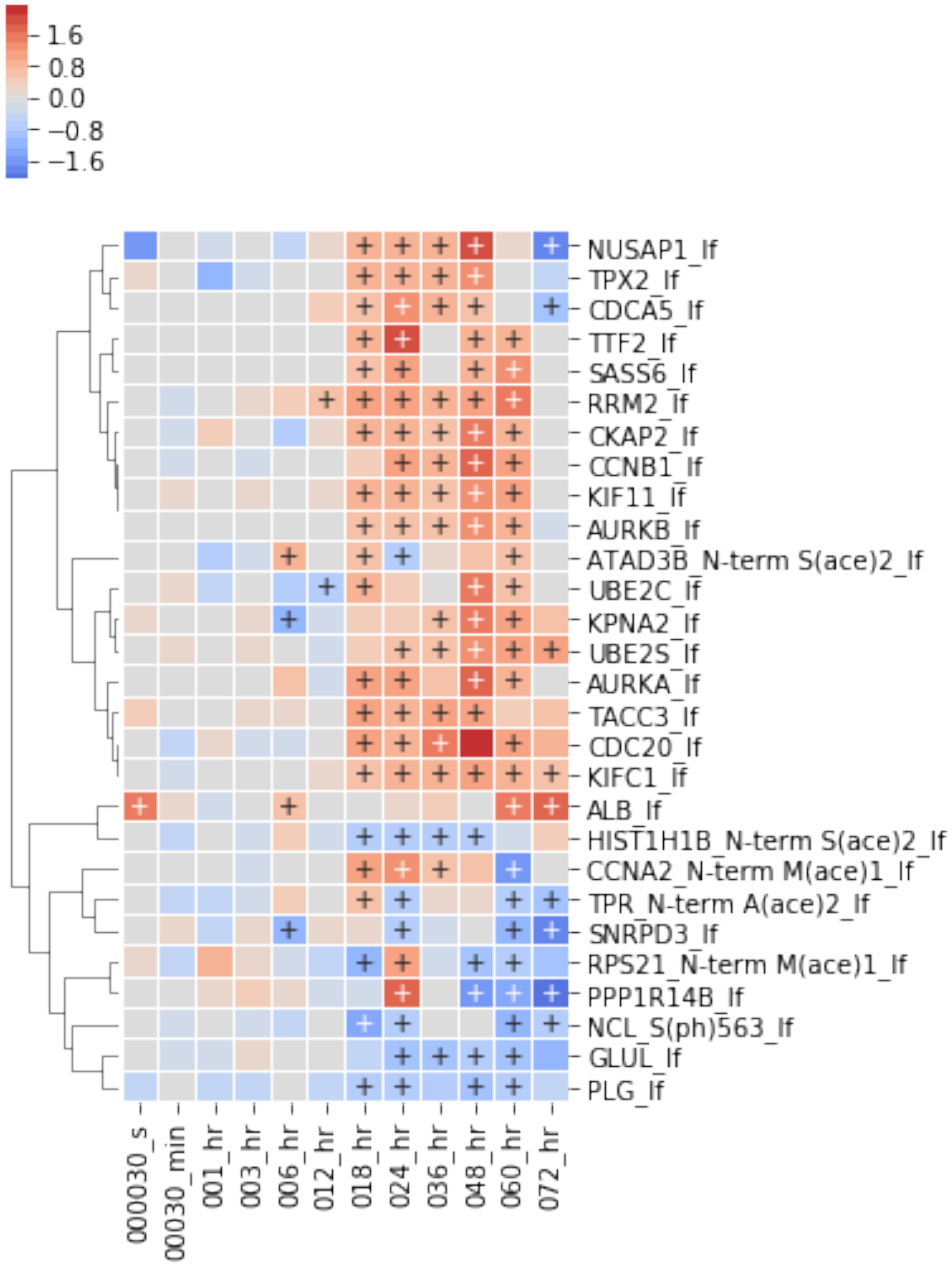

[10]:
```
df = e.run_samples(
    [exp_data.label_free.up.require_n_sig(n_sig=4).id_list,
     exp_data.label_free.down.require_n_sig(n_sig=4).id_list,],
    ['label_free_up', 'label_free_down'],
    gene_set_lib='Reactome_2016')
df.term_name = df.term_name.str.split('_').str.get(0)
```

[ ]:
```
df.heatmap(
    min_sig=1,
    linewidths=0.01,
    convert_to_log=False,
    figsize=(3, 12)
);
print(up_only.shape)
```

[24]:
```
df.head(10)
```

[24]:
```
                                          term_name  rank       p_value  \
0                               cell cycle, mitotic     1  1.545920e-09
1                                        cell cycle     2  7.603575e-09
2  apc/c-mediated degradation of cell cycle proteins     3  7.316318e-09
3                     regulation of mitotic cell cycle     4  7.316318e-09
4                resolution of sister chromatid cohesion     5  1.279403e-06
5                             mitotic prometaphase     6  1.746644e-06
6  regulation of tp53 activity through phosphoryl...     7  8.341746e-07
7                  transcriptional regulation by tp53     8  8.074179e-06
8                                   g2/m transition     9  1.178583e-05
9                          mitotic g2-g2/m phases    10  1.233244e-05

    z_score  combined_score   adj_p_value  \
0 -2.477071       50.253950  2.056073e-07
1 -2.434067       45.504030  2.528189e-07
2 -2.284682       42.799318  2.528189e-07
3 -2.274341       42.605591  2.528189e-07
4 -2.047109       27.777462  2.836010e-05
5 -1.999010       26.502501  3.318624e-05
6 -1.893455       26.502358  2.218904e-05
7 -2.238861       26.254767  1.145954e-04
8 -2.098984       23.820552  1.261703e-04
9 -2.099087       23.726560  1.261703e-04

                                          genes  n_genes            db  \
0  AURKA,AURKB,CCNA2,CCNB1,CDC20,CDCA5,RRM2,TPX2        8  Reactome_2016
1  AURKA,AURKB,CCNA2,CCNB1,CDC20,CDCA5,RRM2,TPX2        8  Reactome_2016
2                  AURKA,AURKB,CCNA2,CCNB1,CDC20        5  Reactome_2016
3                  AURKA,AURKB,CCNA2,CCNB1,CDC20        5  Reactome_2016
```
