## Supplementary material for "A computational framework to explore cellular response mechanisms from multi-omics datasets": S4-Jupyter Notebook [Annotated Gene Set Networks]

### supplement_notebook_4_agn

December 10, 2019

#### 1 Exploring enrichment analysis of HL60 response to bendamustine.

This is the second notebook for the Pino Et. Al. In this notebook we demonstrate how MAGINE can be used to explore the enrichment analysis.

[1]:
```
from IPython.display import display, Image
%matplotlib inline
import networkx as nx
import matplotlib.pyplot as plt
import pandas as pd
pd.set_option('display.precision', 2)
pd.set_option('display.max_colwidth', 50)
%load_ext autoreload
%autoreload 2
```

[2]:
```
# load magine specific tools
from magine.plotting.wordcloud_tools import create_wordcloud
from magine.plotting.heatmaps import heatmap_by_terms
from magine.plotting.venn_diagram_maker import create_venn3, create_venn2

from magine.enrichment import load_enrichment_csv
from magine.networks import visualization as vis
from magine.networks import utils, exporters

from magine.networks.annotated_set import create_subnetwork
from magine.networks.subgraphs import Subgraph
```

#### 2 Exploring enrichment output

##### 2.1 Loading data and networks

[3]:
```
from exp_data import exp_data
```

**Load enrichment array.** This `bendamustine_enrichment.csv.gz` was created by `run_enrichment.py` script. If it doesn't exist, run that file to generate the results. Due to the number of samples, we run this outside a Jupyter notebook as it can take quite a bit of time.

[4]:
```
enrichment_array = load_enrichment_csv('Data/bendamustine_enrichment.csv.gz',␣
↪index_col=0)

enrichment_array['significant'] = False
enrichment_array.loc[(enrichment_array['adj_p_value'] <= 0.05) &
                     (enrichment_array['combined_score'] > 0.0),
                     'significant'] = True
# Remove terms that are not signficant in at least one time point/sample/
↪category
enrichment_array.require_n_sig(
    columns='sample_id',
    index='term_name',
    n_sig=1,
    inplace=True
)
```

[5]:
```
enrichment_array[['term_name', 'db', 'category']].nunique()
```

[5]:
```
term_name    20758
db              52
category        15
dtype: int64
```

[6]:
```
display(enrichment_array.head(5))
```

```
           term_name  rank  combined_score  adj_p_value  \
0     vinblastine-up     1           34.53     2.83e-05
1        mitotane-up     2           31.90     3.64e-05
2  dideoxycytidine-dn     3           30.15     1.38e-04
3         busulfan-dn     4           29.18     1.01e-04
4        mitotane-up     5           28.76     1.39e-04

                                               genes  n_genes sample_id  \
0  ADSS,APOA1,APOE,BSG,BTF3,CANX,COPA,DNAJC9,EI24...       24  000030_s
1  ABI1,ACLY,ALDH3A2,BTF3,CD97,COL4A3BP,COPA,GIT2...       23  000030_s
2  ACADVL,ACLY,ALDH3A2,APOA1,APOA2,APOE,BSG,CANX,...       23  000030_s
3  ABI1,BTF3,CANX,COPA,DNAJC9,FBXW5,FLNA,HMHA1,HP...       22  000030_s
4  ACLY,ADSS,AKAP8L,BTF3,CANX,CD97,COL4A3BP,COPA,...       21  000030_s

          category          db  significant
0  proteomics_both  DrugMatrix         True
1  proteomics_both  DrugMatrix         True
2  proteomics_both  DrugMatrix         True
3  proteomics_both  DrugMatrix         True
4  proteomics_both  DrugMatrix         True
```

[7]:
```
# clean up printing by selecting fewer columns
cols = ['term_name', 'rank', 'combined_score', 'adj_p_value',
```

```
        'n_genes', 'sample_id', 'category']
```

[8]:
```
display(enrichment_array[cols].head(5))
```

```
           term_name  rank  combined_score  adj_p_value  n_genes sample_id  \
0      vinblastine-up     1           34.53     2.83e-05       24  000030_s
1         mitotane-up     2           31.90     3.64e-05       23  000030_s
2  dideoxycytidine-dn     3           30.15     1.38e-04       23  000030_s
3         busulfan-dn     4           29.18     1.01e-04       22  000030_s
4         mitotane-up     5           28.76     1.39e-04       21  000030_s

          category
0  proteomics_both
1  proteomics_both
2  proteomics_both
3  proteomics_both
4  proteomics_both
```

**Load network** Load in the network and initialize Subgraph. We will use this later to construct networks from queries.

[9]:
```
network = nx.read_gpickle('Networks/bendamustine_network_w_attributes.p')
net_sub = Subgraph(network)
```

#### 3 Single database exploration

Here we will focus on the Reactome enrichment.

[10]:
```
reactome_only = enrichment_array.filter_multi(
    db='Reactome_2016', # Only reactome db
)
# This just cleans up the term name
display(reactome_only['term_name'].head(5))
reactome_only['term_name'] = reactome_only['term_name'].str.split('_').str.
 ↪get(0)
display(reactome_only['term_name'].head(5))
```

```
80551    processing of capped intron-containing pre-mrn...
80552                        gene expression_hsa_r-hsa-74160
80553                          mrna splicing_hsa_r-hsa-72172
80554        mrna splicing - major pathway_hsa_r-hsa-72163
80555    transport of mature transcript to cytoplasm_hs...
Name: term_name, dtype: object


80551    processing of capped intron-containing pre-mrna
80552                                    gene expression
```

```
80553                                              mrna splicing
80554                               mrna splicing - major pathway
80555          transport of mature transcript to cytoplasm
Name: term_name, dtype: object
```

[11]:
```python
# we can use a word cloud to view what terms are enriched
word_cloud = create_wordcloud(reactome_only.sig)
word_cloud.plot();
```

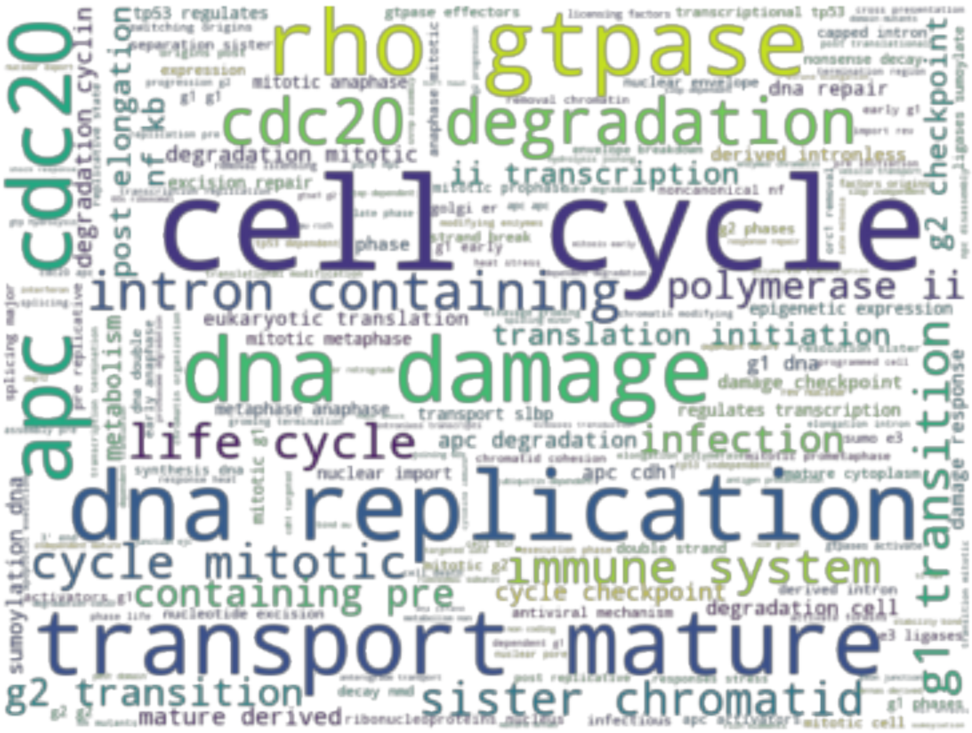


[12]:
```python
word_cloud.data.head(20)
```

[12]:

| | words | counts |
|---|---|---|
| 181 | cell cycle | 325 |
| 208 | dna replication | 160 |
| 249 | rho gtpase | 142 |
| 243 | dna damage | 139 |
| 179 | transport mature | 138 |
| 586 | apc cdc20 | 132 |
| 587 | cdc20 degradation | 122 |
| 175 | intron containing | 121 |
| 353 | immune system | 98 |
| 199 | life cycle | 97 |
| 228 | cycle mitotic | 95 |

```
381          g1 transition      94
355        sister chromatid      86
176          containing pre      80
236           polymerase ii      79
400           g2 transition      79
5                 infection      77
429  translation initiation      77
237        ii transcription      75
261          post elongation      73
```

[ ]:

```

```

#### 3.1 Phospho-SILAC enrichment

##### 3.1.1 Filtering enrichment output

[13]:

```
# subset the data to only look at ph-silac data.
# Later on we will look at label-free, then both.

ph_silac = reactome_only.filter_multi(
    category=['ph_silac_up', 'ph_silac_down'],
)

# arbitrary value that can be easily changed.
ph_silac.require_n_sig(n_sig=3, inplace=True)

not_useful = [
    'gene expression', 'translation',
    'immune system',
    'disease', 'diseases of signal transduction',
    'infectious disease',
    'influenza infection', 'influenza life cycle',
    'influenza viral rna transcription and replication',
]
ph_silac = ph_silac.loc[~ph_silac['term_name'].isin(not_useful)]
ph_silac_copy = ph_silac.copy()

ph_silac.remove_redundant(
    threshold=.5,
    level='dataframe',
    sort_by='combined_score',
    inplace=True
)
```

```
Number of rows went from 159 to 29
```

[14]:
```
ph_silac.heatmap(
    convert_to_log=False,
    cluster_by_set=False,
    cluster_row=False,
    values='combined_score',
    columns=['category', 'sample_id'],
    annotate_sig=True,
    div_colors=True,
    linewidths=.005,
    figsize=(5,12)
);
plt.savefig("ph_silac_enriched.png", dpi=300, bbox_inches='tight')
```

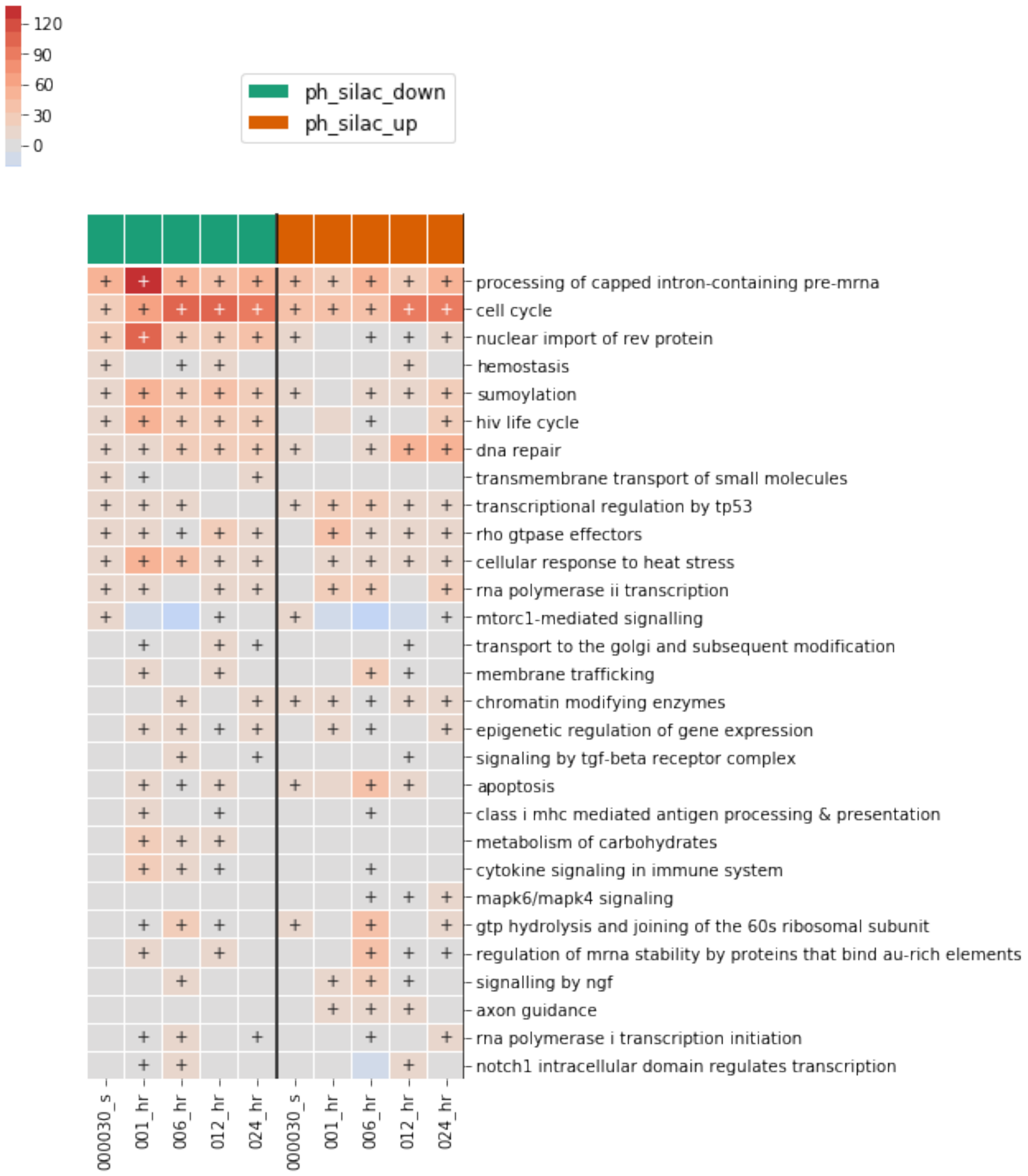


[15]:

```
# This figure shows the process of discovering "too general/not useful term".
# Translation involves various genes that make up individual process
ph_silac_copy.show_terms_below("translation").heatmap(
    convert_to_log=False,
    cluster_by_set=False,
    cluster_row=False,
    values='combined_score',
    columns=['category', 'sample_id'],
    annotate_sig=True,
```

```
    div_colors=True,
    linewidths=.005,
    figsize=(5, 12)
);
```

```
Number of rows went from 159 to 36
```

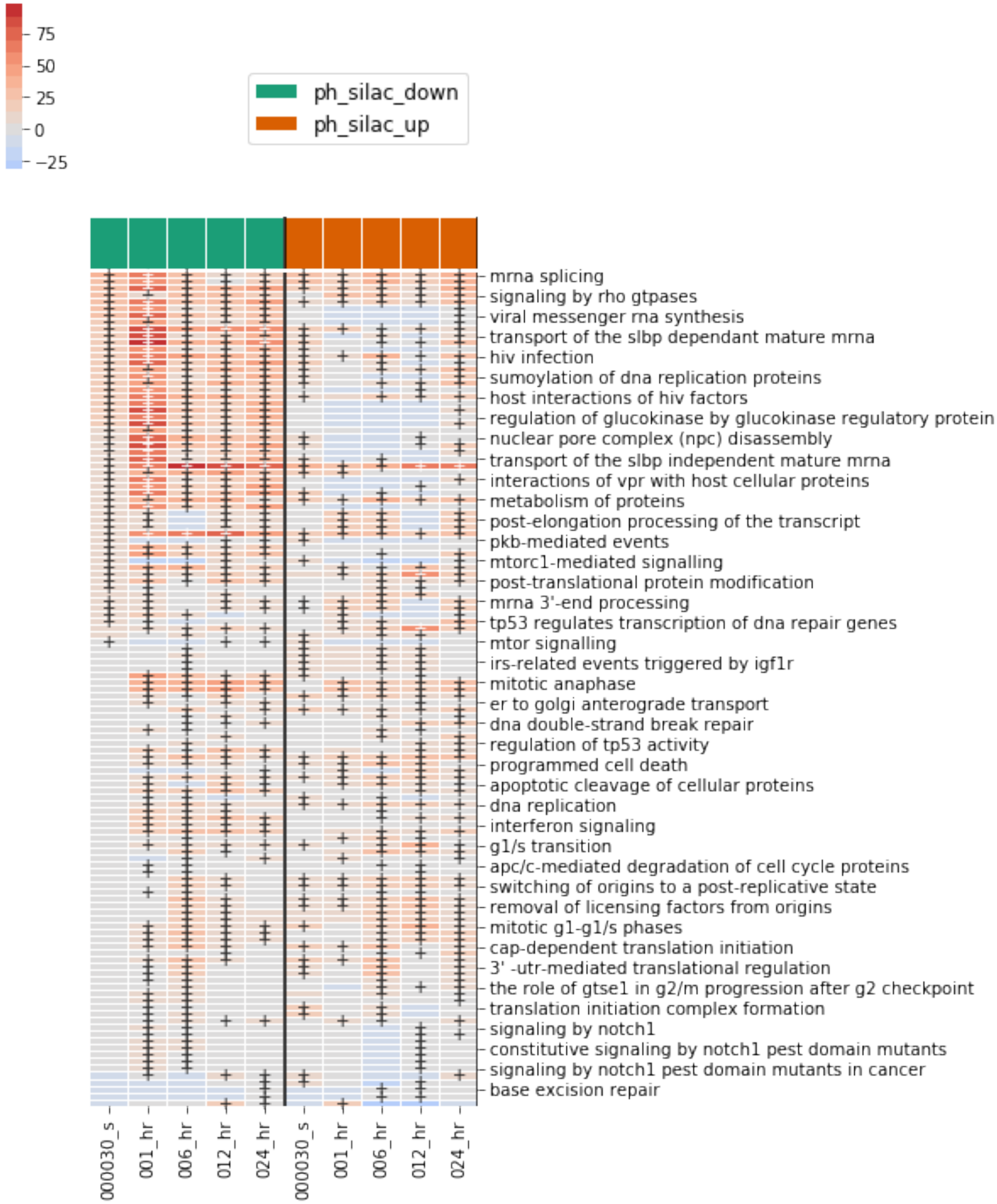

[16]:

```
# We can use the same thought process to find more specific terms
ph_silac_copy.show_terms_below("cell cycle", remove_subset=False, threshold=.5).
↪require_n_sig(n_sig=1).heatmap(
    convert_to_log=False,
    cluster_by_set=False,
    cluster_row=False,
    values='combined_score',
    columns=['category', 'sample_id'],
    annotate_sig=True,
    div_colors=True,
    linewidths=.005,
    figsize=(5, 12)
);
plt.savefig("below_cell_cycle.png", bbox_inches='tight', dpi=300)
```

```
Number of rows went from 159 to 29
```

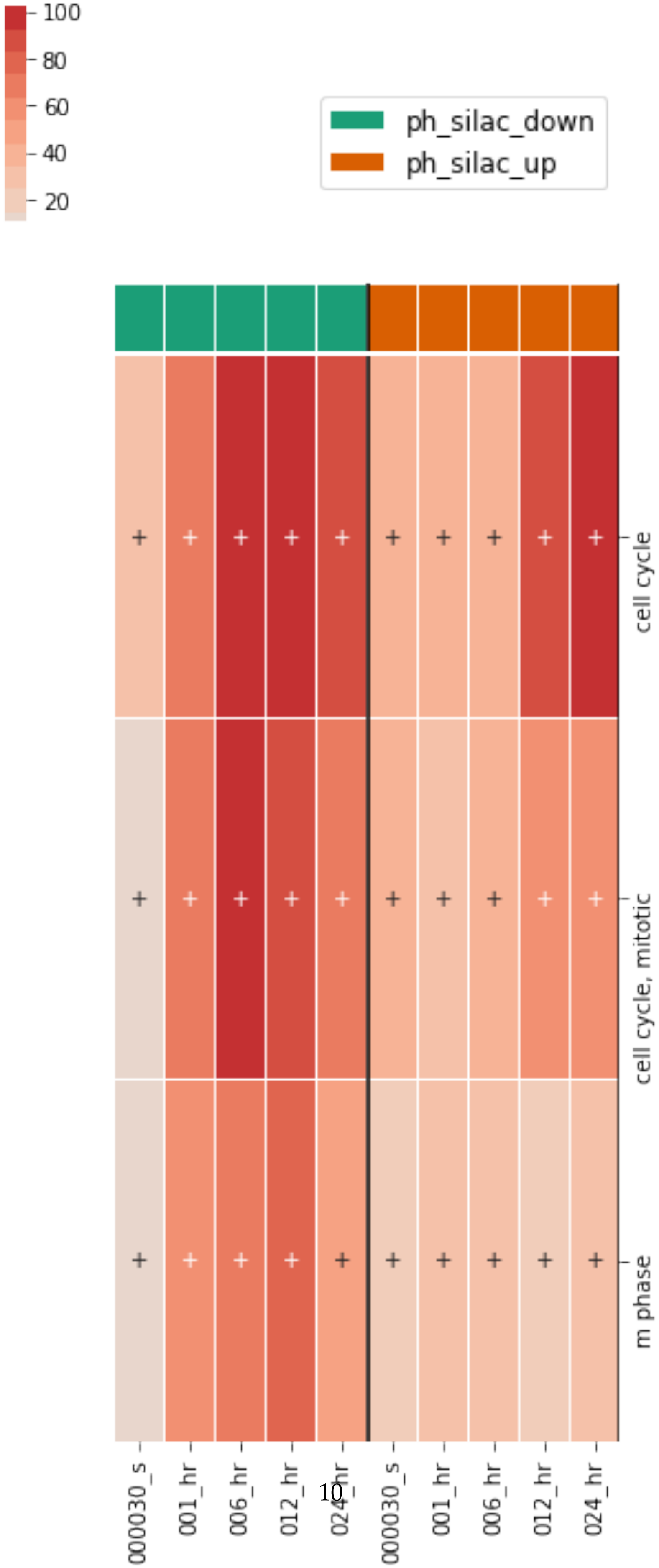
100
80
60
40
20
ph_silac_down
ph_silac_up
cell cycle
cell cycle, mitotic
m phase
000030_s
001_hr
006_hr
012_hr
024_hr
000030_s
001_hr
006_hr
012_hr
024_hr

##### 3.1.2 Creating an annotated set network.

[17]:
```
term_net, mol_net = create_subnetwork(
    ph_silac,
    network=network,
    save_name='ph_silac_ags',
    use_cytoscape=False,
    use_fdr=True, use_threshold=True, min_edges=20
)
```

```
Creating ontology network
```

[18]:
```
vis.draw_cyjs(term_net, default_color='white', layout='concentric',␣
 ↪spacingFactor=2.8)
```

```
<IPython.core.display.HTML object>
```

##### 3.1.3 Known mechanisms of bendamustine

Next, we will demonstrate how to explore known mechanisms. Bendamustine is known to cause dna damage, cell cycle arrest, and apoptosis, so we will start there.

[19]:
```
selected_terms = ['dna repair', 'cell cycle', 'apoptosis']

# extract out lists of genes for each term
g_sets = [ph_silac.sig.term_to_genes(i) for i in selected_terms]

# visualize the overlap
create_venn3(*g_sets+selected_terms, save_name='venn_canonical');
```

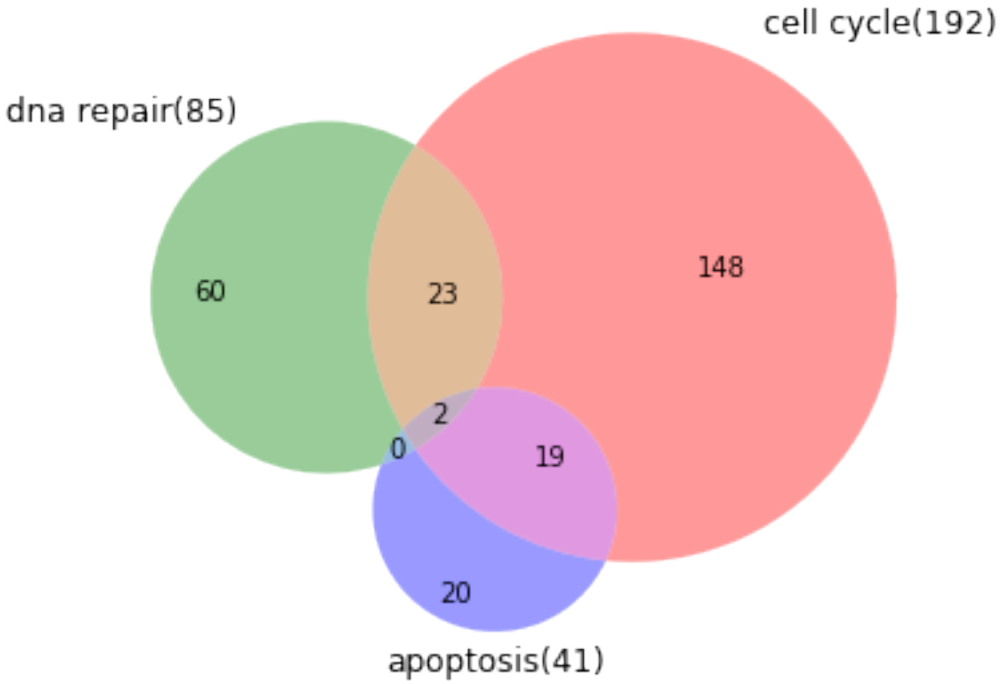


[20]:
```
def print_numbers(term_name):
    genes = reactome_only.sig.term_to_genes(term_name)
    n_sig = len(genes)
    if n_sig:
        n_sig_ptms = len(exp_data.subset(genes).sig.label_list)
        print("{} : {} : {}".format(term_name, n_sig, n_sig_ptms))
print_numbers('cell cycle')
print_numbers('dna repair')
print_numbers('apoptosis')
```

```
cell cycle : 247 : 600
dna repair : 110 : 232
apoptosis : 72 : 138
```

[21]:
```
# though we focused on ph-silac originally, we can see if the terms of interest␣
↪are in other datasets
subset = reactome_only.loc[reactome_only.term_name.isin(selected_terms)].copy()

subset.filter_multi(
    category=['ph_silac_down', 'ph_silac_up',
              'label_free_down', 'label_free_up',
              'silac_up', 'silac_down',
              'rna_up', 'rna_down'],
    inplace=True,
```

```
)

# remove time points and samples that don't contain a significant term
subset.require_n_sig(
    n_sig=1,
    index=['category', 'sample_id'],
    columns='term_name',
    inplace=True
)
subset.require_n_sig(
    n_sig=1,
    index=['sample_id', 'category'],
    columns='term_name',
    inplace=True
)

fig = subset.heatmap(
    convert_to_log=False,
    cluster_by_set=False,
    annotate_sig=True,
    columns=['category', 'sample_id'],
    div_colors=True,
    linewidths=.005,
    figsize=(12, 16)
);
```

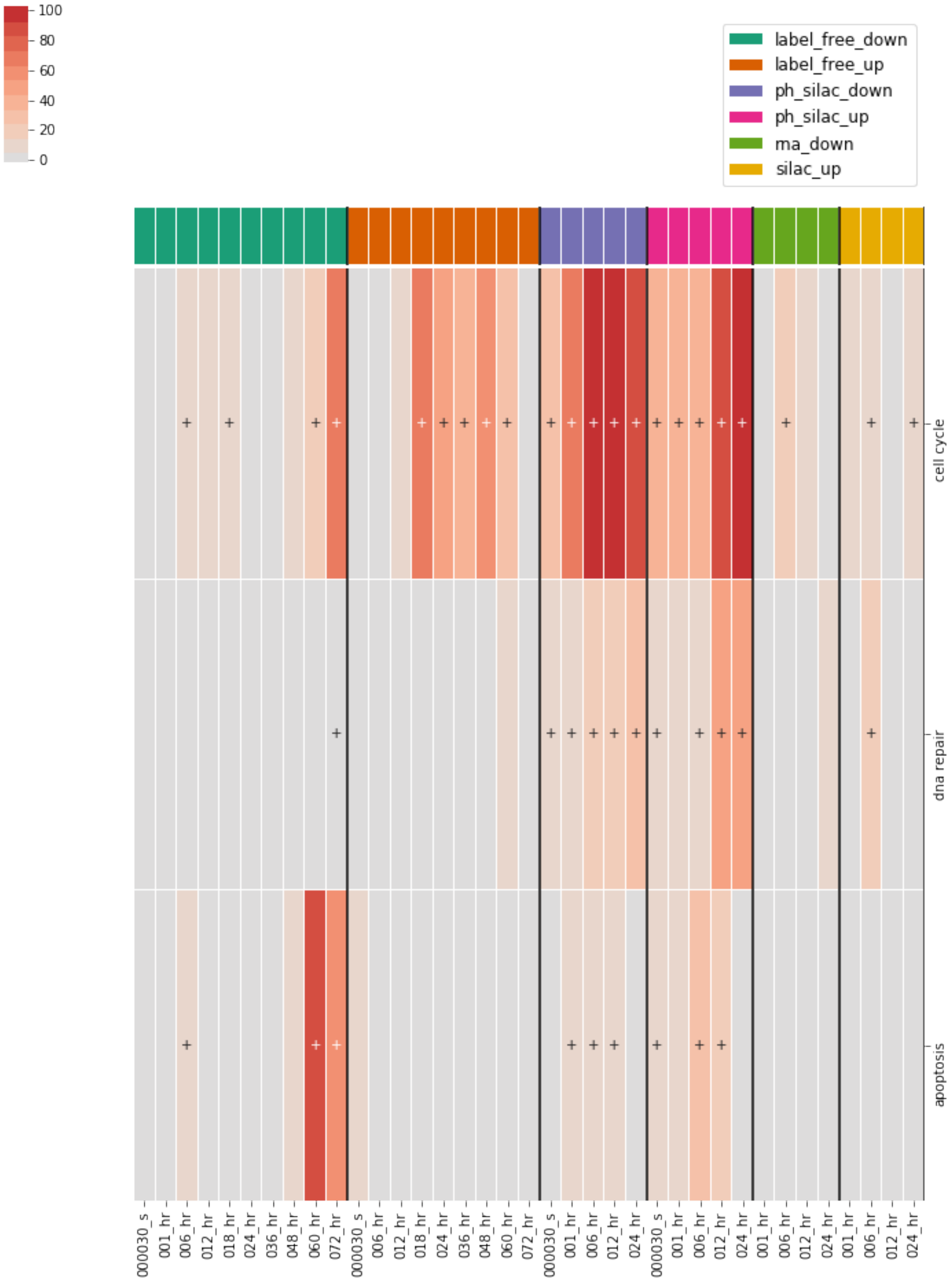


[22]:
```
# subset the data to only include these terms
subset = reactome_only.sig.loc[reactome_only.sig.term_name.
 ↪isin(selected_terms)].copy()
```

```
term_net, mol_net = create_subnetwork(
    subset,
    network=network,
    save_name='apop_dna_cell_cycle',
    use_cytoscape=False,
    use_fdr=True,
    use_threshold=True,
    min_edges=10
)
print(len(mol_net.nodes))
print(len(mol_net.edges))
```

```
Creating ontology network
311
2406
```

[23]:
```
nx.set_node_attributes(term_net, 'white', 'color')
vis.draw_cyjs(term_net, layout='concentric', spacingFactor=2.8)
```

```
<IPython.core.display.HTML object>
```

##### DNA damage response

[24]:
```
# extract dna repair genes
dna_repair_genes = reactome_only.sig.term_to_genes('dna repair')


exp_data.ph_silac.heatmap(
    dna_repair_genes,
    convert_to_log=True,
    subset_index='identifier',
    index='label',
    cluster_row=True,
    rank_index=True,
    min_sig=2,
    num_colors=13,
    linewidths=0.01
);


exp_data.label_free.heatmap(
    dna_repair_genes,
    convert_to_log=True,
    subset_index='identifier',
    index='label',
```

```
    cluster_row=True,
    rank_index=True,
    min_sig=2,
    num_colors=13,
    linewidths=0.01
);


exp_data.rna_seq.heatmap(
    dna_repair_genes,
    convert_to_log=True,
    subset_index='identifier',
    index='label',
    cluster_row=True,
    rank_index=True,
    min_sig=2,
    num_colors=13,
    linewidths=0.01
);
```

```
E:\PycharmProjects\PycharmProjects\Magine\magine\plotting\heatmaps.py:64:
UserWarning:

Empty array after filtering.
```

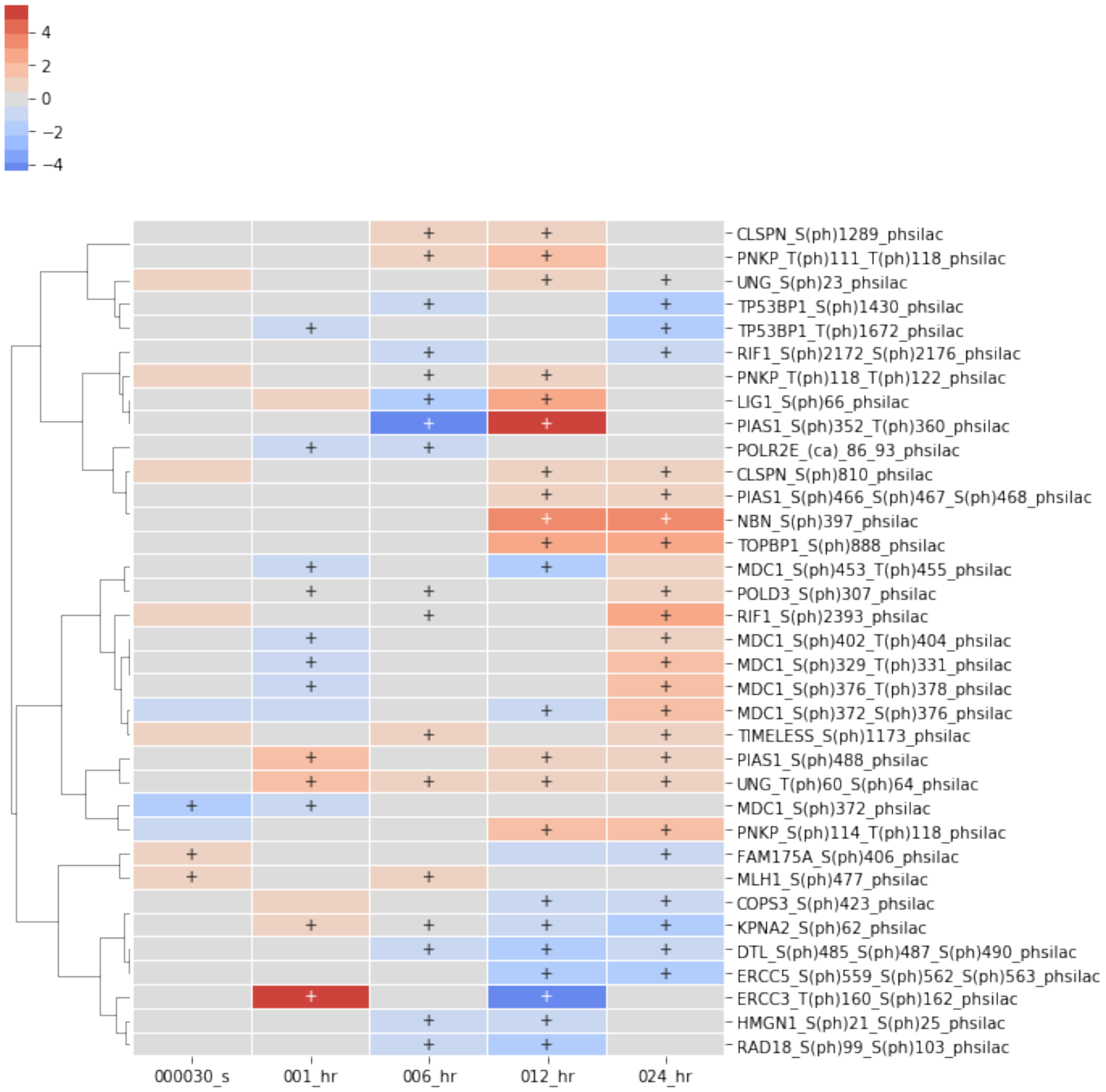

4
2
0
−2
−4
CLSPN_S(ph)1289_phsilac
PNKP_T(ph)111_T(ph)118_phsilac
UNG_S(ph)23_phsilac
TP53BP1_S(ph)1430_phsilac
TP53BP1_T(ph)1672_phsilac
RIF1_S(ph)2172_S(ph)2176_phsilac
PNKP_T(ph)118_T(ph)122_phsilac
LIG1_S(ph)66_phsilac
PIAS1_S(ph)352_T(ph)360_phsilac
POLR2E_(ca)_86_93_phsilac
CLSPN_S(ph)810_phsilac
PIAS1_S(ph)466_S(ph)467_S(ph)468_phsilac
NBN_S(ph)397_phsilac
TOPBP1_S(ph)888_phsilac
MDC1_S(ph)453_T(ph)455_phsilac
POLD3_S(ph)307_phsilac
RIF1_S(ph)2393_phsilac
MDC1_S(ph)402_T(ph)404_phsilac
MDC1_S(ph)329_T(ph)331_phsilac
MDC1_S(ph)376_T(ph)378_phsilac
MDC1_S(ph)372_S(ph)376_phsilac
TIMELESS_S(ph)1173_phsilac
PIAS1_S(ph)488_phsilac
UNG_T(ph)60_S(ph)64_phsilac
MDC1_S(ph)372_phsilac
PNKP_S(ph)114_T(ph)118_phsilac
FAM175A_S(ph)406_phsilac
MLH1_S(ph)477_phsilac
COPS3_S(ph)423_phsilac
KPNA2_S(ph)62_phsilac
DTL_S(ph)485_S(ph)487_S(ph)490_phsilac
ERCC5_S(ph)559_S(ph)562_S(ph)563_phsilac
ERCC3_T(ph)160_S(ph)162_phsilac
HMGN1_S(ph)21_S(ph)25_phsilac
RAD18_S(ph)99_S(ph)103_phsilac
000030_s
001_hr
006_hr
012_hr
024_hr

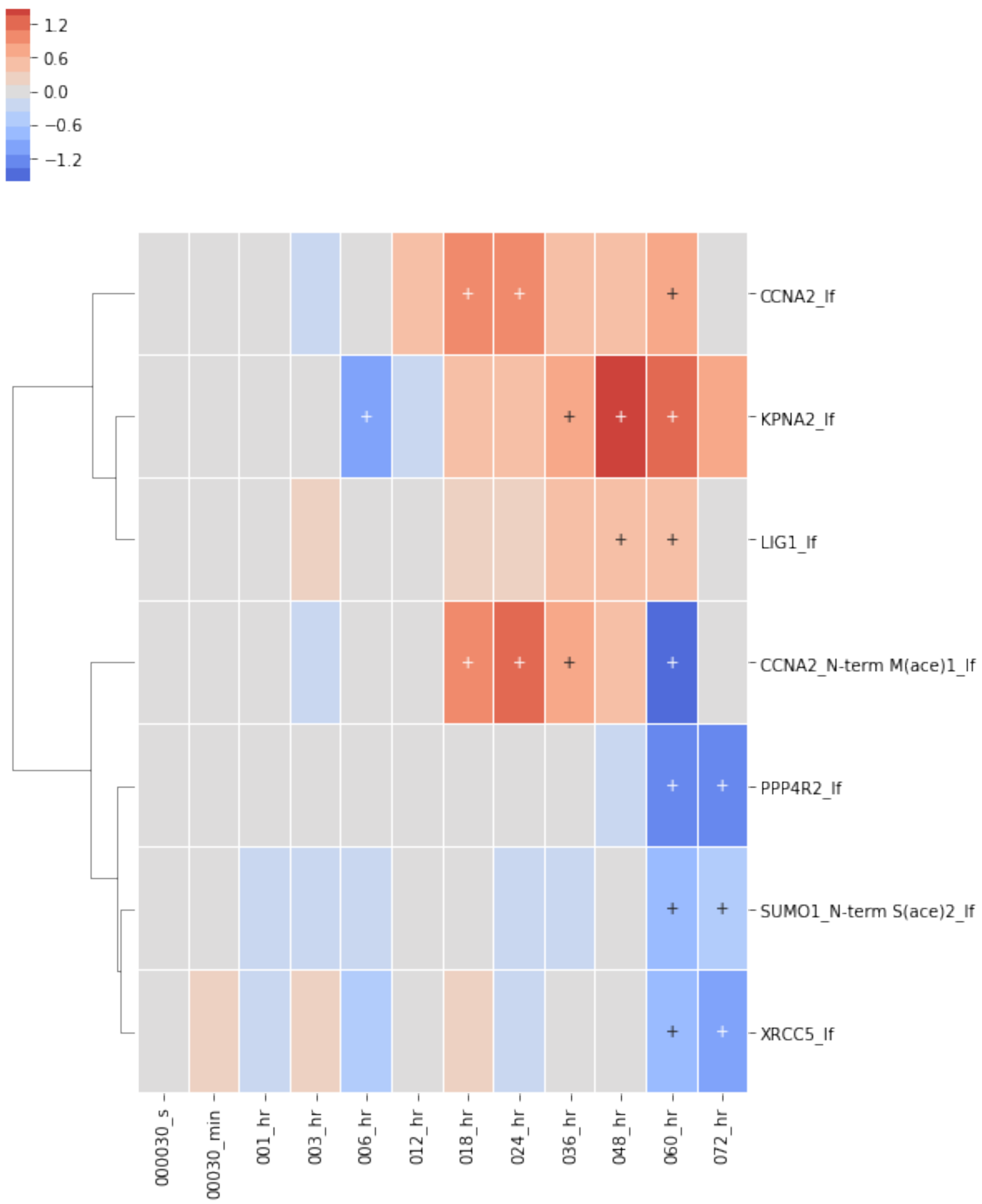


[25]:
```
# look at only first time point
first_tp = exp_data.species.subset(dna_repair_genes, sample_ids=['000030_s']).
 ↪sig

first_tp.heatmap(figsize=(3,6), index='label', linewidths=0.01);
```

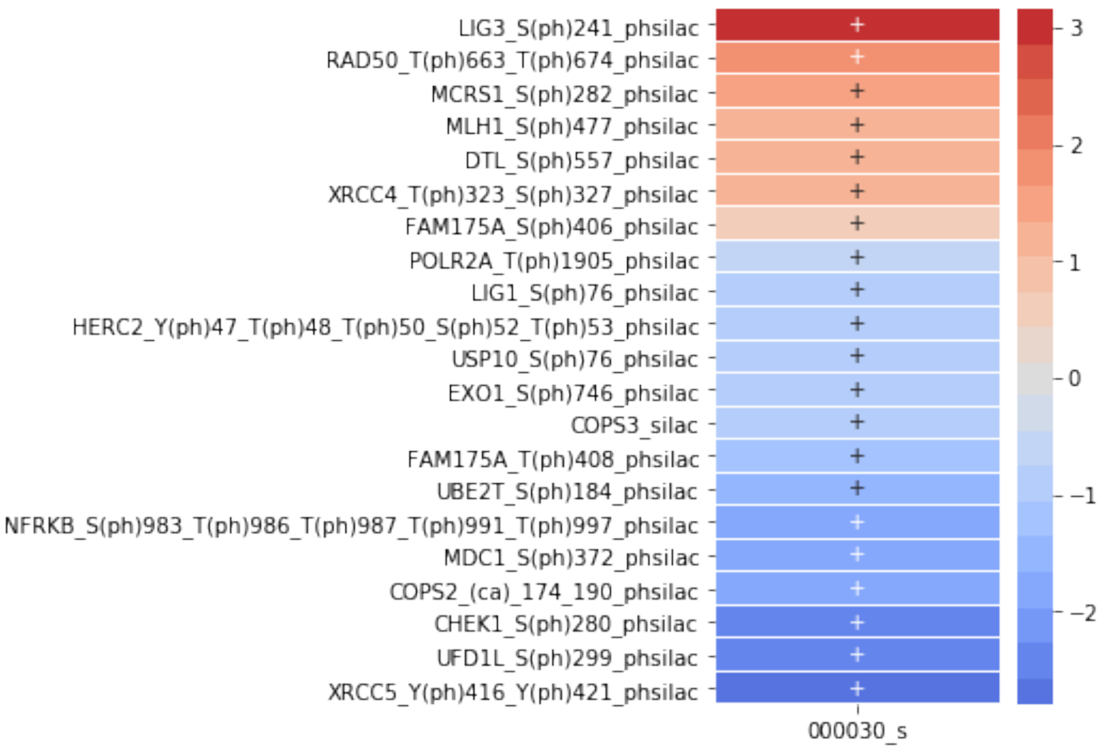


[26]:
```
# Create a network based on the first time point, include dna damage nodes
dna_repair_subnet = net_sub.expand_neighbors(
    nodes=first_tp.id_list,
    upstream=False,
    downstream=True,
    add_interconnecting_edges=True,
    max_dist=1,
    include_only=dna_repair_genes
)
# removes disconnected nodes in network
dna_repair_subnet = utils.delete_disconnected_network(dna_repair_subnet)

colors = dict()
for i in dna_repair_subnet.nodes:
    if i in first_tp.id_list:
        colors[i] = 'lightgreen'
    else:
        colors[i] = 'lightblue'
nx.set_node_attributes(dna_repair_subnet, colors, 'color')
```

```
Node not in graph
```

[27]:
```
vis.draw_cyjs(dna_repair_subnet, layout='cose-bilkent', spacingFactor=2.)
```

```
<IPython.core.display.HTML object>
```

[28]:
```
vis.draw_cyjs(dna_repair_subnet, layout='concentric', spacingFactor=1.)
```

```
<IPython.core.display.HTML object>
```

**Cell cycle**

[29]:
```
cell_cycle_genes = reactome_only.sig.term_to_genes('cell cycle')


exp_data.ph_silac.heatmap(
    cell_cycle_genes,
    subset_index='identifier',
    index='label',
    cluster_row=True,
    rank_index=True,
    min_sig=2,
    num_colors=13,
    linewidths=0.01,
    figsize=(6, 20),
    y_tick_labels=True
);

exp_data.label_free.heatmap(
    cell_cycle_genes,
    subset_index='identifier',
    index='label',
    cluster_row=True,
    rank_index=True,
    min_sig=2,
    num_colors=13,
    linewidths=0.01,
    figsize=(6,16)
);

exp_data.rna_seq.heatmap(
    cell_cycle_genes,
    subset_index='identifier',
    index='label',
    cluster_row=True,
    rank_index=True,
    min_sig=1,
    num_colors=13,
    linewidths=0.01,
    figsize=(6,10)
```

```
);
```

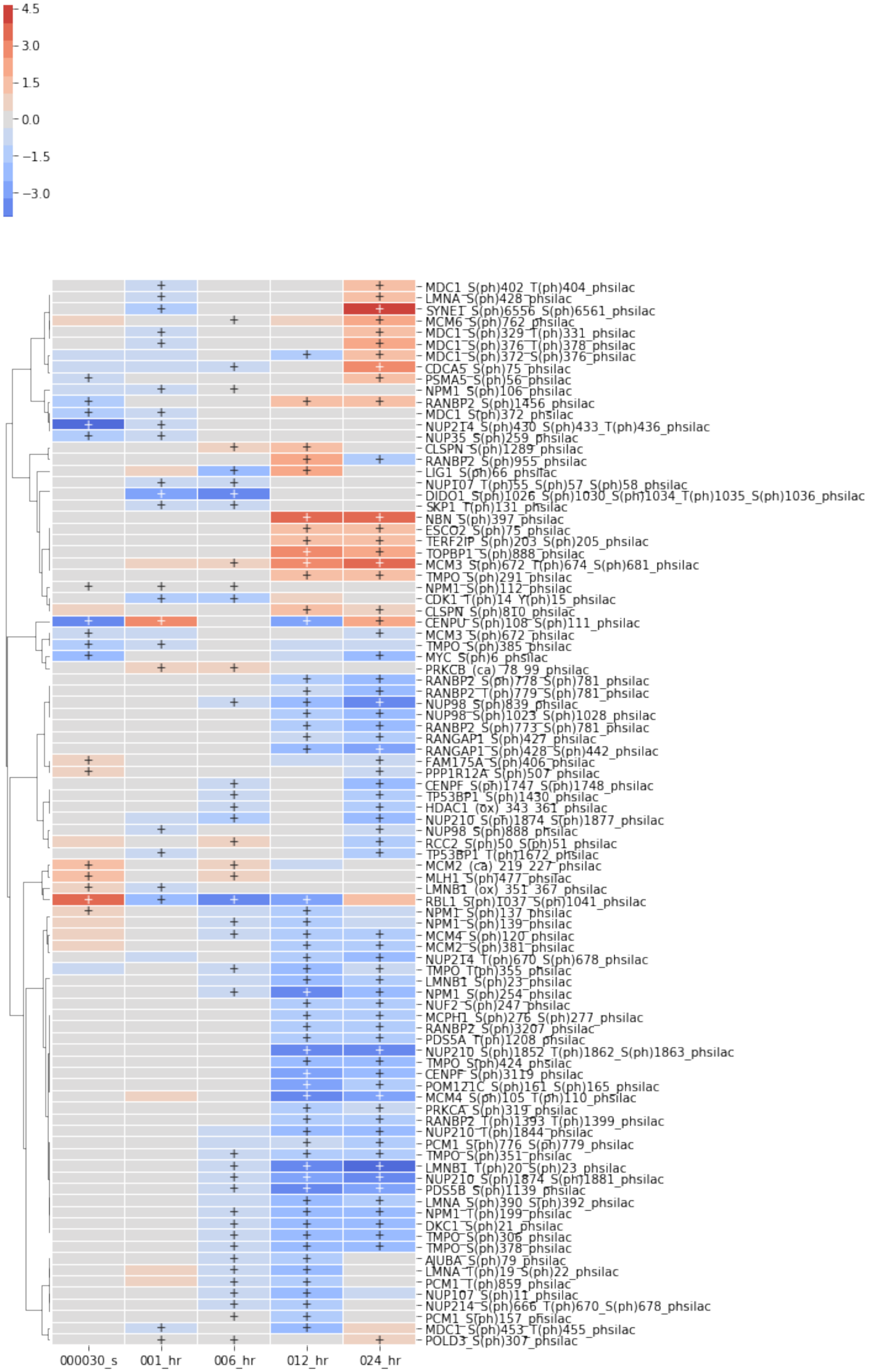
4.5
3.0
1.5
0.0
−1.5
−3.0
MDC1_S(ph)402_T(ph)404_phsilac
LMNA_S(ph)428_phsilac
SYNE1_S(ph)6556_S(ph)6561_phsilac
MCM6_S(ph)762_phsilac
MDC1_S(ph)329_T(ph)331_phsilac
MDC1_S(ph)376_T(ph)378_phsilac
MDC1_S(ph)372_S(ph)376_phsilac
CDCA5_S(ph)75_phsilac
PSMA5_S(ph)56_phsilac
NPM1_S(ph)106_phsilac
RANBP2_S(ph)1456_phsilac
MDC1_S(ph)372_phsilac
NUP214_S(ph)430_S(ph)433_T(ph)436_phsilac
NUP35_S(ph)259_phsilac
CLSPN_S(ph)1289_phsilac
RANBP2_S(ph)955_phsilac
LIG1_S(ph)66_phsilac
NUP107_T(ph)55_S(ph)57_S(ph)58_phsilac
DIDO1_S(ph)1026_S(ph)1030_S(ph)1034_T(ph)1035_S(ph)1036_phsilac
SKP1_T(ph)131_phsilac
NBN_S(ph)397_phsilac
ESCO2_S(ph)75_phsilac
TERF2IP_S(ph)203_S(ph)205_phsilac
TOPBP1_S(ph)888_phsilac
MCM3_S(ph)672_T(ph)674_S(ph)681_phsilac
TMPO_S(ph)291_phsilac
NPM1_S(ph)112_phsilac
CDK1_T(ph)14_Y(ph)15_phsilac
CLSPN_S(ph)810_phsilac
CENPU_S(ph)108_S(ph)111_phsilac
MCM3_S(ph)672_phsilac
TMPO_S(ph)385_phsilac
MYC_S(ph)6_phsilac
PRKCB_(ca)_78_99_phsilac
RANBP2_S(ph)778_S(ph)781_phsilac
RANBP2_T(ph)779_S(ph)781_phsilac
NUP98_S(ph)839_phsilac
NUP98_S(ph)1023_S(ph)1028_phsilac
RANBP2_S(ph)773_S(ph)781_phsilac
RANGAP1_S(ph)427_phsilac
RANGAP1_S(ph)428_S(ph)442_phsilac
FAM175A_S(ph)406_phsilac
PPP1R12A_S(ph)507_phsilac
CENPF_S(ph)1747_S(ph)1748_phsilac
TP53BP1_S(ph)1430_phsilac
HDAC1_(ox)_343_361_phsilac
NUP210_S(ph)1874_S(ph)1877_phsilac
NUP98_S(ph)888_phsilac
RCC2_S(ph)50_S(ph)51_phsilac
TP53BP1_T(ph)1672_phsilac
MCM2_(ca)_219_227_phsilac
MLH1_S(ph)477_phsilac
LMNB1_(ox)_351_367_phsilac
RBL1_S(ph)1037_S(ph)1041_phsilac
NPM1_S(ph)137_phsilac
NPM1_S(ph)139_phsilac
MCM4_S(ph)120_phsilac
MCM2_S(ph)381_phsilac
NUP214_T(ph)670_S(ph)678_phsilac
TMPO_T(ph)355_phsilac
LMNB1_S(ph)23_phsilac
NPM1_S(ph)254_phsilac
NUF2_S(ph)247_phsilac
MCPH1_S(ph)276_S(ph)277_phsilac
RANBP2_S(ph)3207_phsilac
PDS5A_T(ph)1208_phsilac
NUP210_S(ph)1852_T(ph)1862_S(ph)1863_phsilac
TMPO_S(ph)424_phsilac
CENPF_S(ph)3119_phsilac
POM121C_S(ph)161_S(ph)165_phsilac
MCM4_S(ph)105_T(ph)110_phsilac
PRKCA_S(ph)319_phsilac
RANBP2_T(ph)1393_T(ph)1399_phsilac
NUP210_T(ph)1844_phsilac
PCM1_S(ph)776_S(ph)779_phsilac
TMPO_S(ph)351_phsilac
LMNB1_T(ph)20_S(ph)23_phsilac
NUP210_S(ph)1874_S(ph)1881_phsilac
PDS5B_S(ph)1139_phsilac
LMNA_S(ph)390_S(ph)392_phsilac
NPM1_T(ph)199_phsilac
DKC1_S(ph)21_phsilac
TMPO_S(ph)306_phsilac
TMPO_S(ph)378_phsilac
AJUBA_S(ph)79_phsilac
LMNA_T(ph)19_S(ph)22_phsilac
PCM1_T(ph)859_phsilac
NUP107_S(ph)11_phsilac
NUP214_S(ph)666_T(ph)670_S(ph)678_phsilac
PCM1_S(ph)157_phsilac
MDC1_S(ph)453_T(ph)455_phsilac
POLD3_S(ph)307_phsilac
000030_s
001_hr
006_hr
012_hr
024_hr

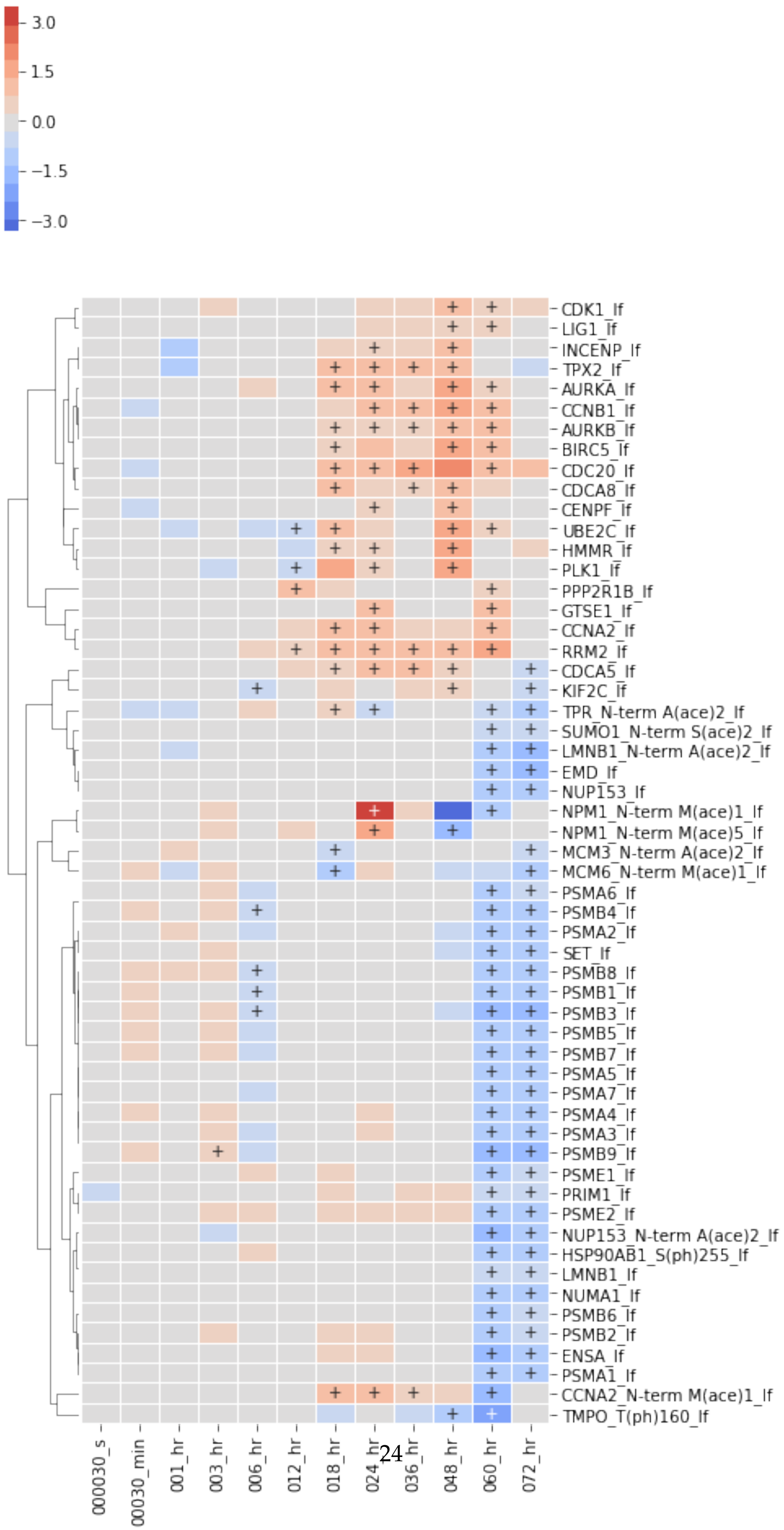
3.0
1.5
0.0
−1.5
−3.0
CDK1_lf
LIG1_lf
INCENP_lf
TPX2_lf
AURKA_lf
CCNB1_lf
AURKB_lf
BIRC5_lf
CDC20_lf
CDCA8_lf
CENPF_lf
UBE2C_lf
HMMR_lf
PLK1_lf
PPP2R1B_lf
GTSE1_lf
CCNA2_lf
RRM2_lf
CDCA5_lf
KIF2C_lf
TPR_N-term A(ace)2_lf
SUMO1_N-term S(ace)2_lf
LMNB1_N-term A(ace)2_lf
EMD_lf
NUP153_lf
NPM1_N-term M(ace)1_lf
NPM1_N-term M(ace)5_lf
MCM3_N-term A(ace)2_lf
MCM6_N-term M(ace)1_lf
PSMA6_lf
PSMB4_lf
PSMA2_lf
SET_lf
PSMB8_lf
PSMB1_lf
PSMB3_lf
PSMB5_lf
PSMB7_lf
PSMA5_lf
PSMA7_lf
PSMA4_lf
PSMA3_lf
PSMB9_lf
PSME1_lf
PRIM1_lf
PSME2_lf
NUP153_N-term A(ace)2_lf
HSP90AB1_S(ph)255_lf
LMNB1_lf
NUMA1_lf
PSMB6_lf
PSMB2_lf
ENSA_lf
PSMA1_lf
CCNA2_N-term M(ace)1_lf
TMPO_T(ph)160_lf
000030_s
00030_min
001_hr
003_hr
006_hr
012_hr
018_hr
024_hr
036_hr
048_hr
060_hr
072_hr

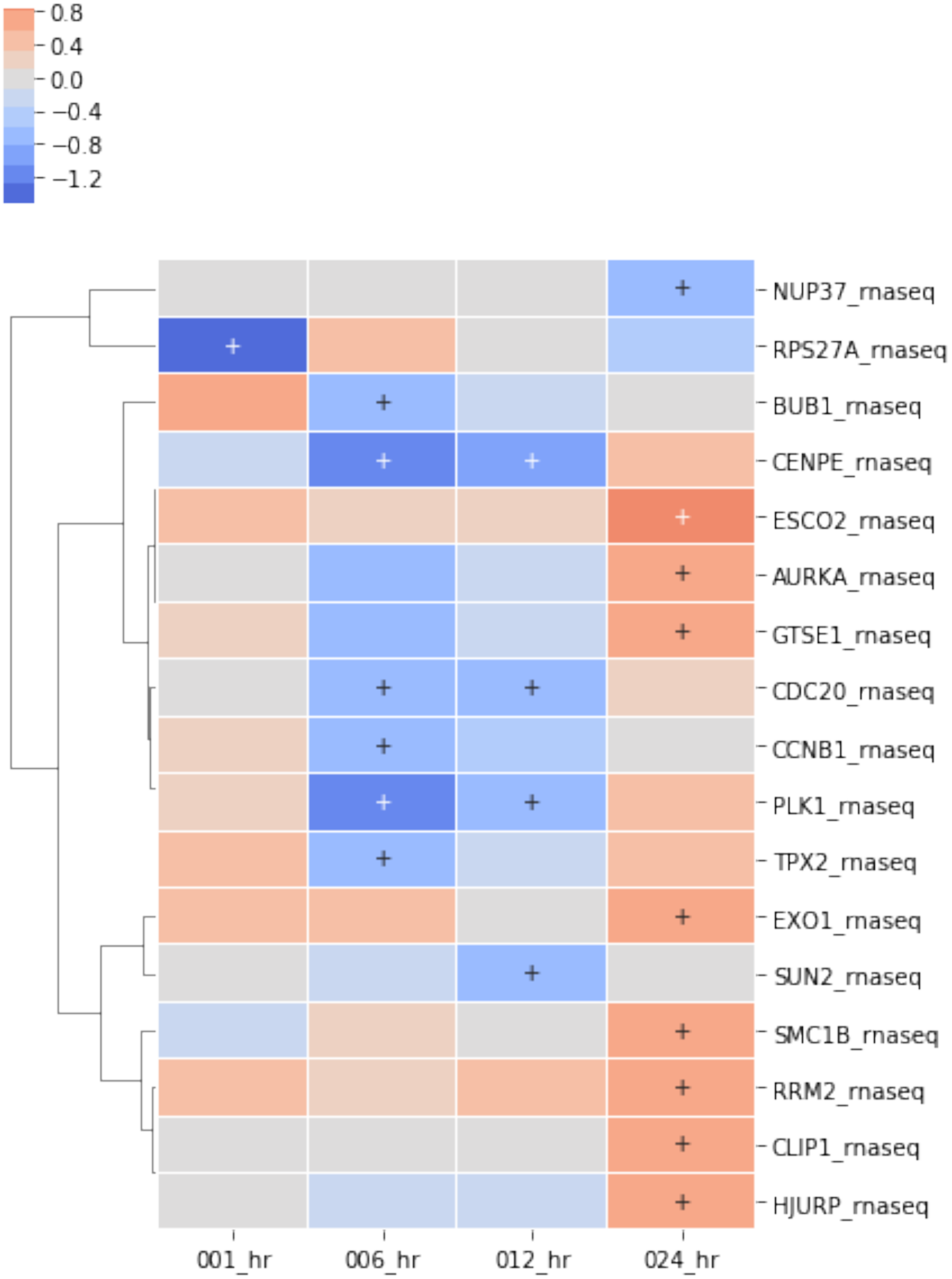
0.8
0.4
0.0
−0.4
−0.8
−1.2
NUP37_rnaseq
RPS27A_rnaseq
BUB1_rnaseq
CENPE_rnaseq
ESCO2_rnaseq
AURKA_rnaseq
GTSE1_rnaseq
CDC20_rnaseq
CCNB1_rnaseq
PLK1_rnaseq
TPX2_rnaseq
EXO1_rnaseq
SUN2_rnaseq
SMC1B_rnaseq
RRM2_rnaseq
CLIP1_rnaseq
HJURP_rnaseq
001_hr
006_hr
012_hr
024_hr

[30]:
```
apoptosis_genes = reactome_only.sig.term_to_genes('apoptosis')

exp_data.ph_silac.heatmap(
    apoptosis_genes,
    subset_index='identifier',
    index='label',
    cluster_row=False,
    rank_index=True,
    min_sig=1,
    num_colors=13,
    linewidths=0.01,
    figsize=(6,16)
);


exp_data.label_free.heatmap(
    apoptosis_genes,
    subset_index='identifier',
    index='label',
    cluster_row=False,
    rank_index=True,
    min_sig=1,
    num_colors=13,
    linewidths=0.01,
    figsize=(6,16)
);
exp_data.rna_seq.heatmap(
    apoptosis_genes,
    subset_index='identifier',
    index='label',
    cluster_row=True,
    rank_index=True,
    min_sig=1,
    num_colors=13,
    linewidths=0.01,
    figsize=(6,10)
);
```

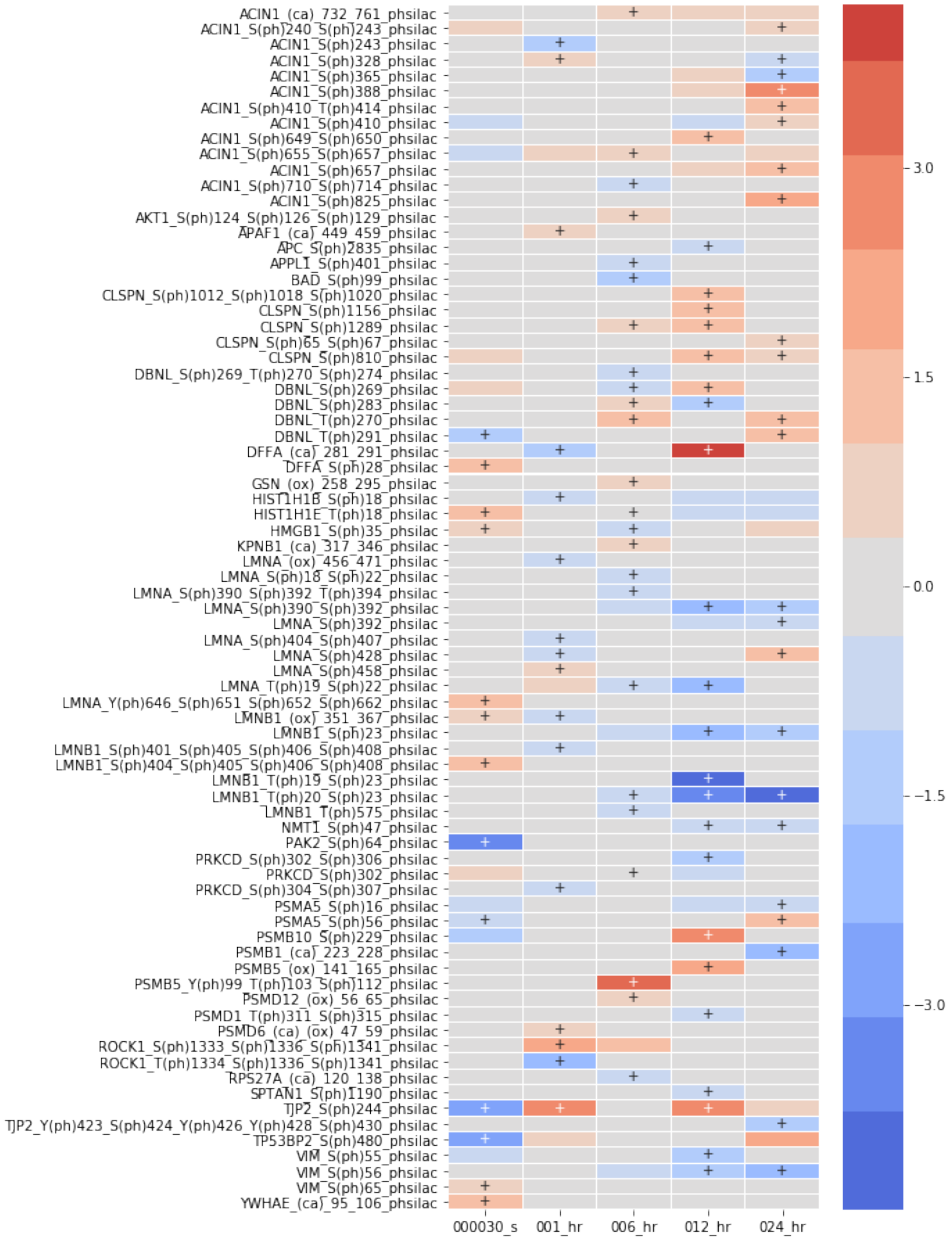
ACIN1_(ca)_732_761_phsilac
ACIN1_S(ph)240_S(ph)243_phsilac
ACIN1_S(ph)243_phsilac
ACIN1_S(ph)328_phsilac
ACIN1_S(ph)365_phsilac
ACIN1_S(ph)388_phsilac
ACIN1_S(ph)410_T(ph)414_phsilac
ACIN1_S(ph)410_phsilac
ACIN1_S(ph)649_S(ph)650_phsilac
ACIN1_S(ph)655_S(ph)657_phsilac
ACIN1_S(ph)657_phsilac
ACIN1_S(ph)710_S(ph)714_phsilac
ACIN1_S(ph)825_phsilac
AKT1_S(ph)124_S(ph)126_S(ph)129_phsilac
APAF1_(ca)_449_459_phsilac
APC_S(ph)2835_phsilac
APPL1_S(ph)401_phsilac
BAD_S(ph)99_phsilac
CLSPN_S(ph)1012_S(ph)1018_S(ph)1020_phsilac
CLSPN_S(ph)1156_phsilac
CLSPN_S(ph)1289_phsilac
CLSPN_S(ph)65_S(ph)67_phsilac
CLSPN_S(ph)810_phsilac
DBNL_S(ph)269_T(ph)270_S(ph)274_phsilac
DBNL_S(ph)269_phsilac
DBNL_S(ph)283_phsilac
DBNL_T(ph)270_phsilac
DBNL_T(ph)291_phsilac
DFFA_(ca)_281_291_phsilac
DFFA_S(ph)28_phsilac
GSN_(ox)_258_295_phsilac
HIST1H1B_S(ph)18_phsilac
HIST1H1E_T(ph)18_phsilac
HMGB1_S(ph)35_phsilac
KPNB1_(ca)_317_346_phsilac
LMNA_(ox)_456_471_phsilac
LMNA_S(ph)18_S(ph)22_phsilac
LMNA_S(ph)390_S(ph)392_T(ph)394_phsilac
LMNA_S(ph)390_S(ph)392_phsilac
LMNA_S(ph)392_phsilac
LMNA_S(ph)404_S(ph)407_phsilac
LMNA_S(ph)428_phsilac
LMNA_S(ph)458_phsilac
LMNA_T(ph)19_S(ph)22_phsilac
LMNA_Y(ph)646_S(ph)651_S(ph)652_S(ph)662_phsilac
LMNB1_(ox)_351_367_phsilac
LMNB1_S(ph)23_phsilac
LMNB1_S(ph)401_S(ph)405_S(ph)406_S(ph)408_phsilac
LMNB1_S(ph)404_S(ph)405_S(ph)406_S(ph)408_phsilac
LMNB1_T(ph)19_S(ph)23_phsilac
LMNB1_T(ph)20_S(ph)23_phsilac
LMNB1_T(ph)575_phsilac
NMT1_S(ph)47_phsilac
PAK2_S(ph)64_phsilac
PRKCD_S(ph)302_S(ph)306_phsilac
PRKCD_S(ph)302_phsilac
PRKCD_S(ph)304_S(ph)307_phsilac
PSMA5_S(ph)16_phsilac
PSMA5_S(ph)56_phsilac
PSMB10_S(ph)229_phsilac
PSMB1_(ca)_223_228_phsilac
PSMB5_(ox)_141_165_phsilac
PSMB5_Y(ph)99_T(ph)103_S(ph)112_phsilac
PSMD12_(ox)_56_65_phsilac
PSMD1_T(ph)311_S(ph)315_phsilac
PSMD6_(ca)_(ox)_47_59_phsilac
ROCK1_S(ph)1333_S(ph)1336_S(ph)1341_phsilac
ROCK1_T(ph)1334_S(ph)1336_S(ph)1341_phsilac
RPS27A_(ca)_120_138_phsilac
SPTAN1_S(ph)1190_phsilac
TJP2_S(ph)244_phsilac
TJP2_Y(ph)423_S(ph)424_Y(ph)426_Y(ph)428_S(ph)430_phsilac
TP53BP2_S(ph)480_phsilac
VIM_S(ph)55_phsilac
VIM_S(ph)56_phsilac
VIM_S(ph)65_phsilac
YWHAE_(ca)_95_106_phsilac
000030_s
001_hr
006_hr
012_hr
024_hr
3.0
1.5
0.0
−1.5
−3.0

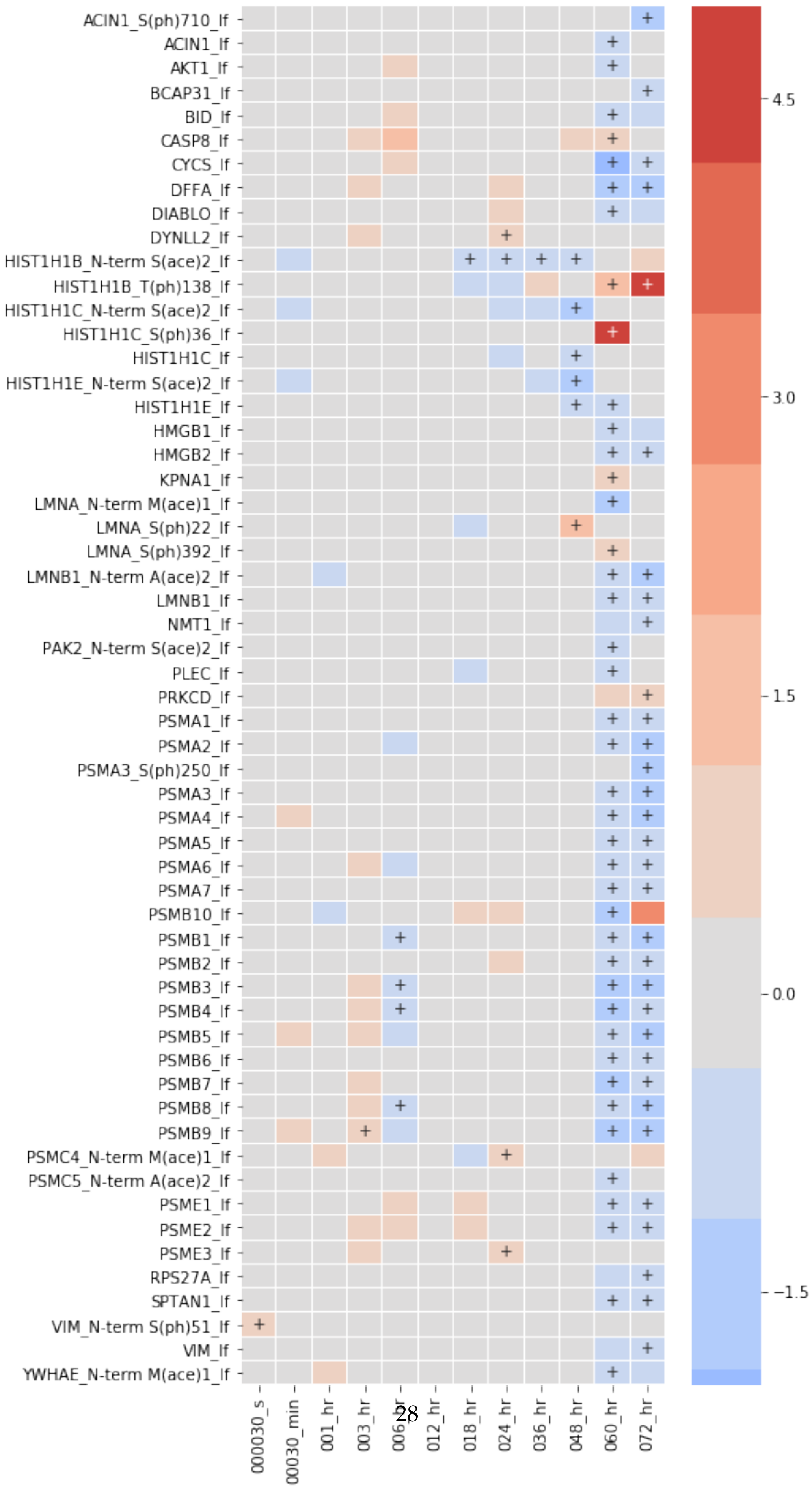
ACIN1_S(ph)710_lf
ACIN1_lf
AKT1_lf
BCAP31_lf
BID_lf
CASP8_lf
CYCS_lf
DFFA_lf
DIABLO_lf
DYNLL2_lf
HIST1H1B_N-term S(ace)2_lf
HIST1H1B_T(ph)138_lf
HIST1H1C_N-term S(ace)2_lf
HIST1H1C_S(ph)36_lf
HIST1H1C_lf
HIST1H1E_N-term S(ace)2_lf
HIST1H1E_lf
HMGB1_lf
HMGB2_lf
KPNA1_lf
LMNA_N-term M(ace)1_lf
LMNA_S(ph)22_lf
LMNA_S(ph)392_lf
LMNB1_N-term A(ace)2_lf
LMNB1_lf
NMT1_lf
PAK2_N-term S(ace)2_lf
PLEC_lf
PRKCD_lf
PSMA1_lf
PSMA2_lf
PSMA3_S(ph)250_lf
PSMA3_lf
PSMA4_lf
PSMA5_lf
PSMA6_lf
PSMA7_lf
PSMB10_lf
PSMB1_lf
PSMB2_lf
PSMB3_lf
PSMB4_lf
PSMB5_lf
PSMB6_lf
PSMB7_lf
PSMB8_lf
PSMB9_lf
PSMC4_N-term M(ace)1_lf
PSMC5_N-term A(ace)2_lf
PSME1_lf
PSME2_lf
PSME3_lf
RPS27A_lf
SPTAN1_lf
VIM_N-term S(ph)51_lf
VIM_lf
YWHAE_N-term M(ace)1_lf
000030_s
00030_min
001_hr
003_hr
006_hr
012_hr
018_hr
024_hr
036_hr
048_hr
060_hr
072_hr
4.5
3.0
1.5
0.0
−1.5

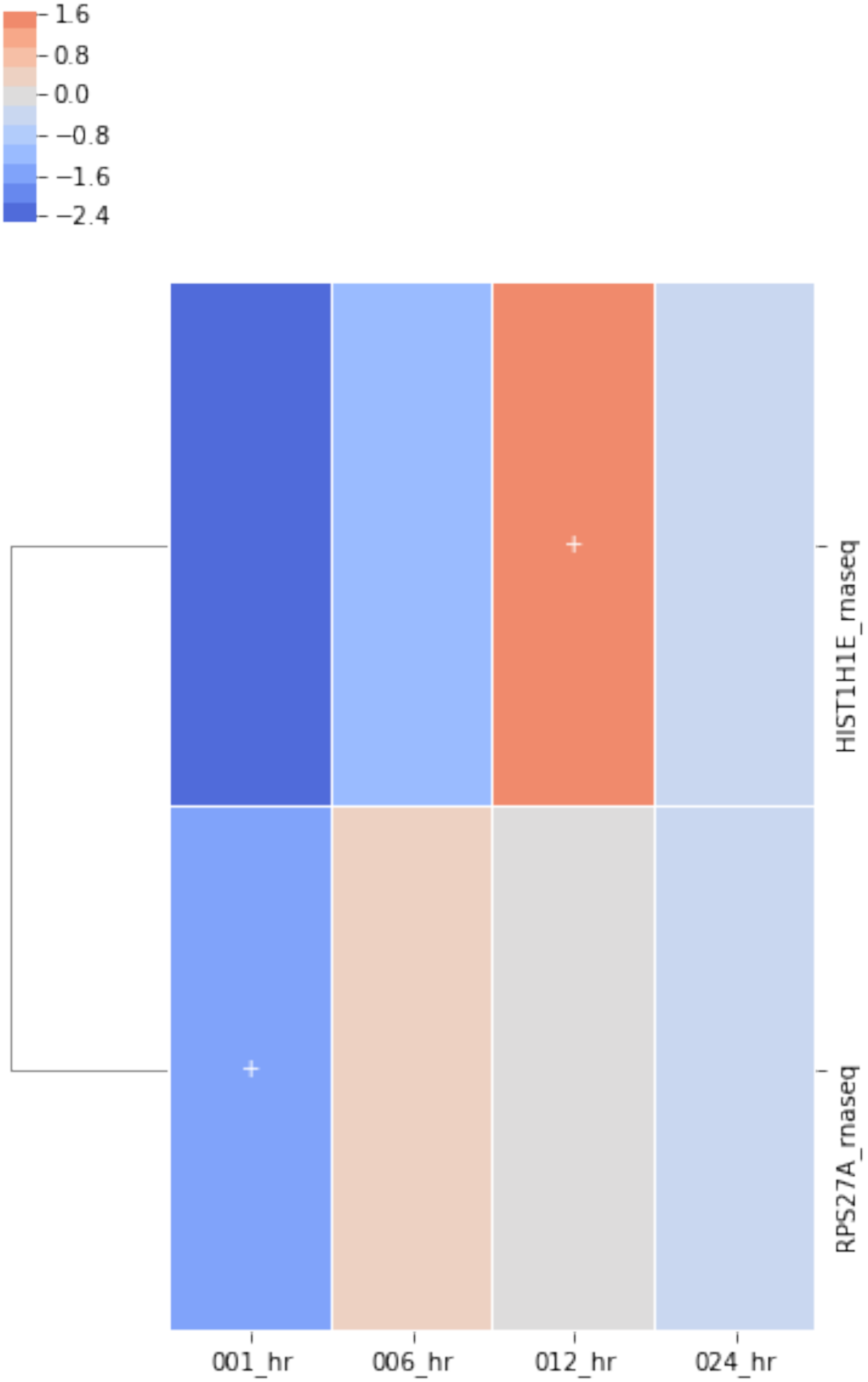

1.6
0.8
0.0
−0.8
−1.6
−2.4
HIST1H1E_rnaseq
RPS27A_rnaseq
001_hr
006_hr
012_hr
024_hr

[31]:
```
print("DNA repair")
for i in exp_data.exp_methods:
    print("\t", i, len(exp_data[i].subset(dna_repair_genes).sig.id_list),
          len(exp_data[i].subset(dna_repair_genes).sig.label_list))
print("Cell cycle")
for i in exp_data.exp_methods:
    print("\t", i, len(exp_data[i].subset(cell_cycle_genes).sig.id_list),
          len(exp_data[i].subset(cell_cycle_genes).sig.label_list))
print("Apoptosis")
for i in exp_data.exp_methods:
    print("\t", i, len(exp_data[i].subset(apoptosis_genes).sig.id_list),
          len(exp_data[i].subset(apoptosis_genes).sig.label_list))
```

```
DNA repair
         silac 7 7
         ph_silac 80 173
         HILIC 0 0
         C18 0 0
         label_free 43 47
         rna_seq 5 5
Cell cycle
         silac 10 10
         ph_silac 153 444
         HILIC 0 0
         C18 0 0
         label_free 111 129
         rna_seq 17 17
Apoptosis
         silac 2 2
         ph_silac 33 77
         HILIC 0 0
         C18 0 0
         label_free 47 57
         rna_seq 2 2
```

[32]:
```
# create plots grouping together data for each platform

genes_in_labels = utils.create_dict_from_node_attributes(mol_net, 'termName')

# phosph-silac
heatmap_by_terms(
    exp_data.ph_silac,
    convert_to_log=True,
    index='label',
    term_labels=list(genes_in_labels.keys()),
    term_sets=list(genes_in_labels.values()),
    div_colors=True,
```

```
    linewidths=0.01,
    min_sig=3,
    annotate_sig=True,
    cluster_col=False,
    cluster_row=False,
    y_tick_labels=True,
    figsize=(6, 12)
);

# label free
heatmap_by_terms(
    exp_data.label_free,
    convert_to_log=True,
    index='label',
    term_labels=list(genes_in_labels.keys()),
    term_sets=list(genes_in_labels.values()),
    div_colors=True,
    linewidths=0.01,
    min_sig=3,
    annotate_sig=True,
    cluster_col=False,
    cluster_row=False,
    y_tick_labels=True,
    figsize=(8, 12)
);

# silac
heatmap_by_terms(
    exp_data.silac,
    convert_to_log=True,
    index='label',
    term_labels=list(genes_in_labels.keys()),
    term_sets=list(genes_in_labels.values()),
    div_colors=True,
    linewidths=0.01,
    min_sig=2,
    annotate_sig=True,
    cluster_col=False,
    cluster_row=False,
    y_tick_labels=True,
    figsize=(4, 4)
);



# rna
heatmap_by_terms(
```

```
    exp_data.rna_seq,
    convert_to_log=True,
    index='label',
    term_labels=list(genes_in_labels.keys()),
    term_sets=list(genes_in_labels.values()),
    div_colors=True,
    linewidths=0.01,
    min_sig=1,
    annotate_sig=True,
    cluster_col=False,
    cluster_row=False,
    y_tick_labels=True,
    figsize=(5, 10)
);
```

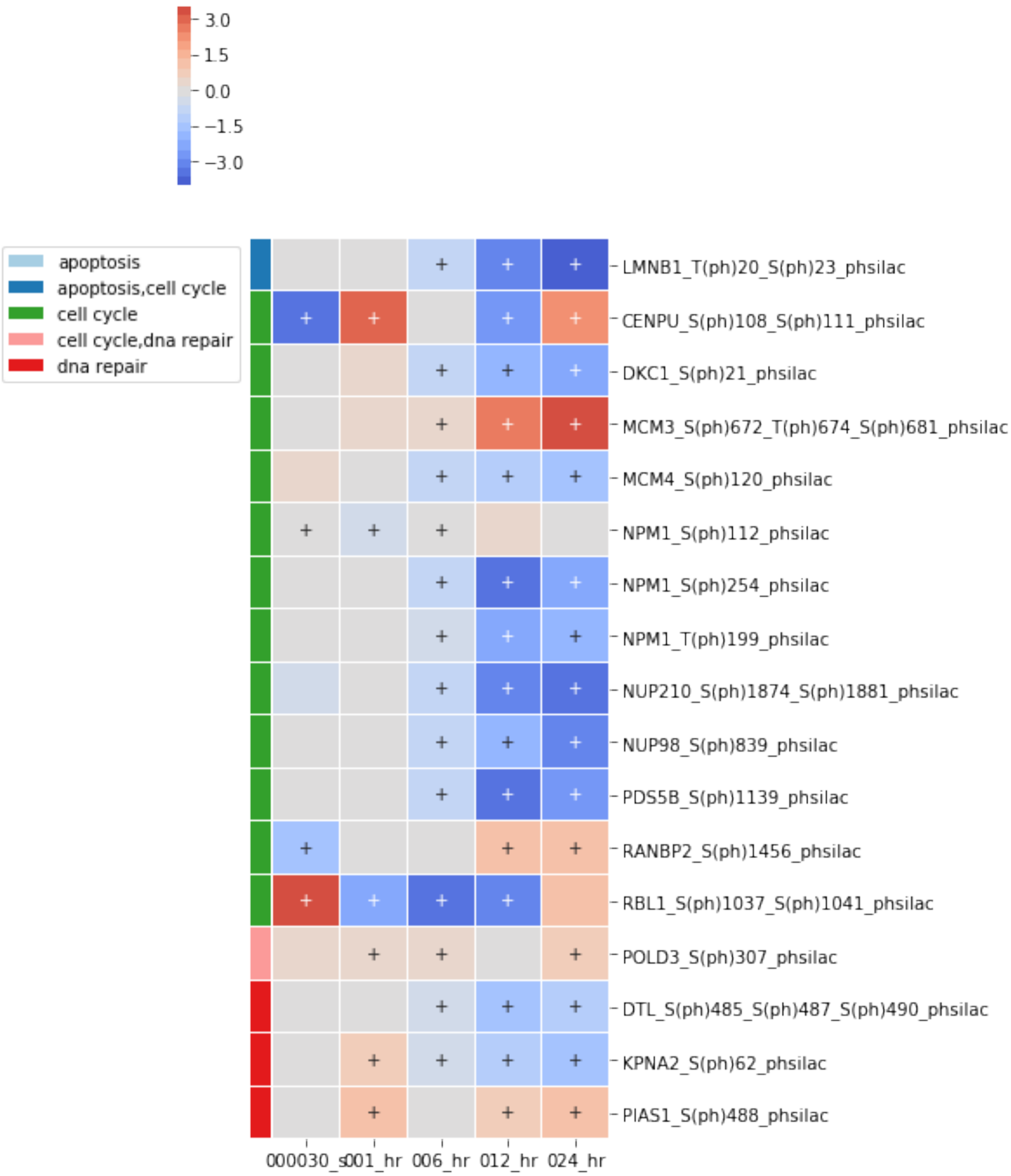

3.0
1.5
0.0
−1.5
−3.0
apoptosis
apoptosis,cell cycle
cell cycle
cell cycle,dna repair
dna repair
LMNB1_T(ph)20_S(ph)23_phsilac
CENPU_S(ph)108_S(ph)111_phsilac
DKC1_S(ph)21_phsilac
MCM3_S(ph)672_T(ph)674_S(ph)681_phsilac
MCM4_S(ph)120_phsilac
NPM1_S(ph)112_phsilac
NPM1_S(ph)254_phsilac
NPM1_T(ph)199_phsilac
NUP210_S(ph)1874_S(ph)1881_phsilac
NUP98_S(ph)839_phsilac
PDS5B_S(ph)1139_phsilac
RANBP2_S(ph)1456_phsilac
RBL1_S(ph)1037_S(ph)1041_phsilac
POLD3_S(ph)307_phsilac
DTL_S(ph)485_S(ph)487_S(ph)490_phsilac
KPNA2_S(ph)62_phsilac
PIAS1_S(ph)488_phsilac
000030_s001_hr 006_hr 012_hr 024_hr

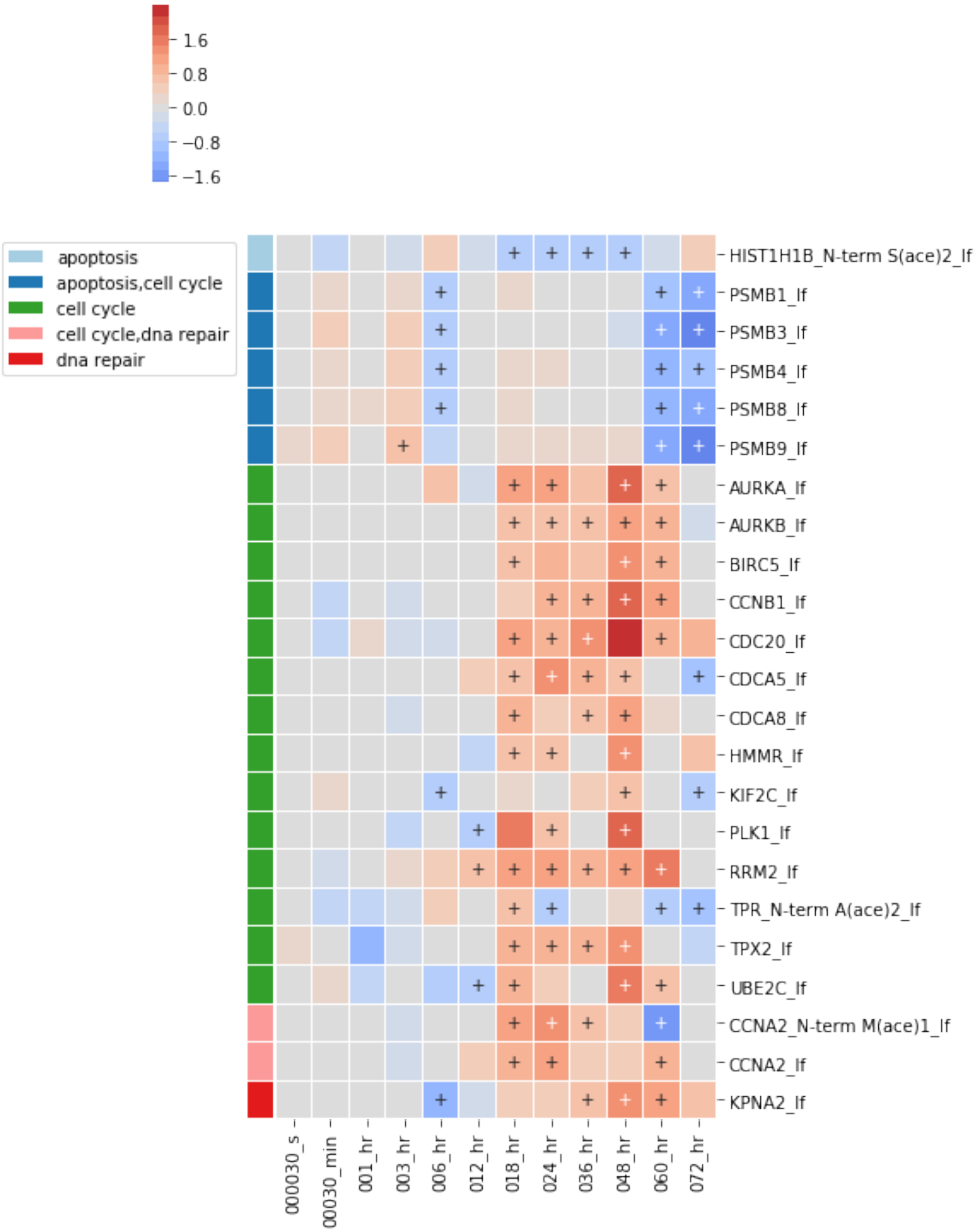
1.6
0.8
0.0
−0.8
−1.6
apoptosis
apoptosis,cell cycle
cell cycle
cell cycle,dna repair
dna repair
HIST1H1B_N-term S(ace)2_lf
PSMB1_lf
PSMB3_lf
PSMB4_lf
PSMB8_lf
PSMB9_lf
AURKA_lf
AURKB_lf
BIRC5_lf
CCNB1_lf
CDC20_lf
CDCA5_lf
CDCA8_lf
HMMR_lf
KIF2C_lf
PLK1_lf
RRM2_lf
TPR_N-term A(ace)2_lf
TPX2_lf
UBE2C_lf
CCNA2_N-term M(ace)1_lf
CCNA2_lf
KPNA2_lf
000030_s
00030_min
001_hr
003_hr
006_hr
012_hr
018_hr
024_hr
036_hr
048_hr
060_hr
072_hr

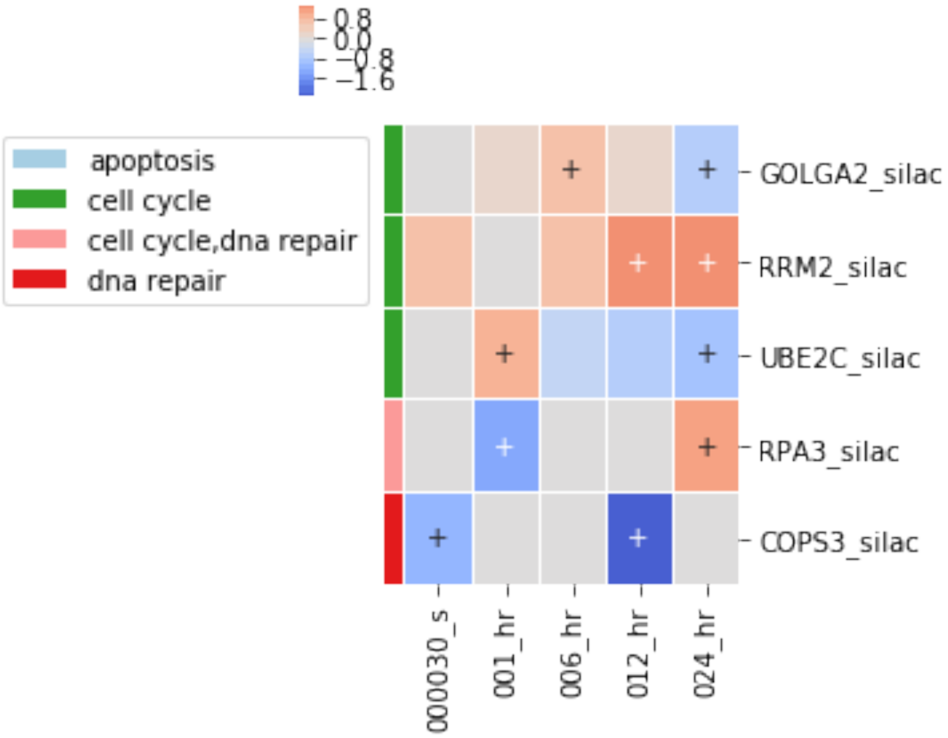
0.8
0.0
−0.8
−1.6
apoptosis
cell cycle
cell cycle,dna repair
dna repair
GOLGA2_silac
RRM2_silac
UBE2C_silac
RPA3_silac
COPS3_silac
000030_s
001_hr
006_hr
012_hr
024_hr

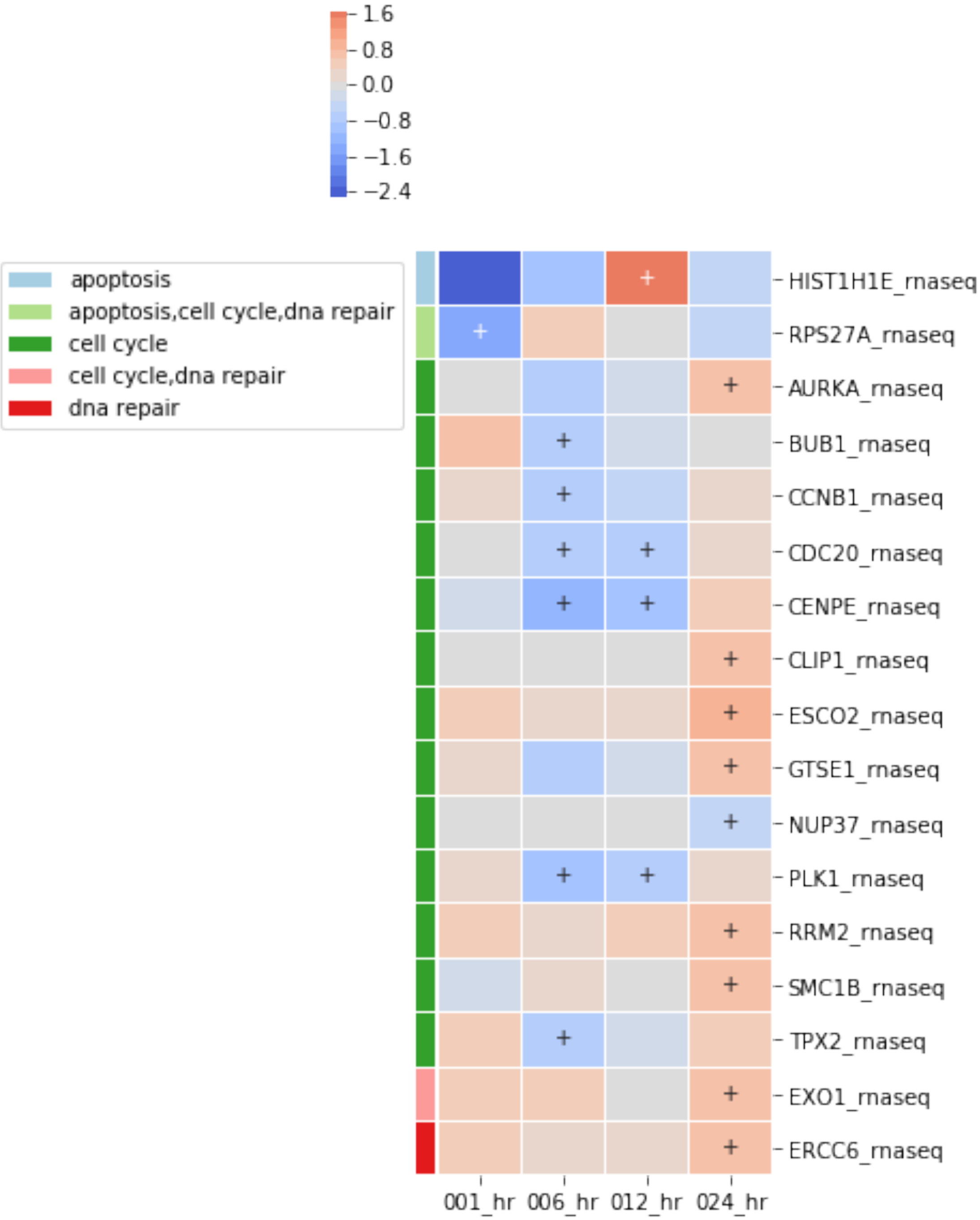


##### 3.1.4 Lets look at only up-regulated ph-silac

[33]:
```
ph_silac_up = reactome_only.filter_multi(category=['ph_silac_up'])

not_useful = [
```

```
    'gene expression', 'translation',
    'immune system',
    'disease', 'diseases of signal transduction',
    'infectious disease',
    'influenza infection', 'influenza life cycle',
    'influenza viral rna transcription and replication',
]

ph_silac_up = ph_silac_up.loc[~ph_silac_up['term_name'].isin(not_useful)]

print("Number of terms before filtering base on minimum time points : {}\
      ".format(len(ph_silac_up.term_name.unique())))

ph_silac_up.require_n_sig(
    index='term_name',
    columns='sample_id',
    n_sig=3,
    inplace=True
)
ph_silac_up_copy = ph_silac_up.copy()
print("Number of terms after filtering base on minimum time points : {}\
      ".format(len(ph_silac_up.term_name.unique())))

fig = ph_silac_up_copy.heatmap(figsize=(4, 24));
fig.savefig('ph_silac.png', dpi=300, bbox_inches='tight')

ph_silac_up.remove_redundant(
    threshold=.7,
    level='sample',
    inplace=True,
    sort_by='combined_score'
)

ph_silac_up.remove_redundant(
    threshold=.7,
    level='dataframe',
    inplace=True,
    sort_by='combined_score'
)
print("Number of genes before : {}".format(len(ph_silac_up_copy.
 ↪all_genes_from_df())))
print("Number of genes after : {}".format(len(ph_silac_up.all_genes_from_df())))
fig = ph_silac_up.heatmap(
    convert_to_log=False,
    cluster_by_set=False,
    cluster_row=False,
    values='combined_score',
```

```
    annotate_sig=True,
    div_colors=True,
    linewidths=.005,
    figsize=(4,8)
);
fig.savefig('ph_silac_slimmed.png', dpi=300, bbox_inches='tight')
```

```
Number of terms before filtering base on minimum time points : 560
Number of terms after filtering base on minimum time points : 83
Number of rows went from 83 to 30
Number of rows went from 30 to 17
Number of genes before : 570
Number of genes after : 550
```

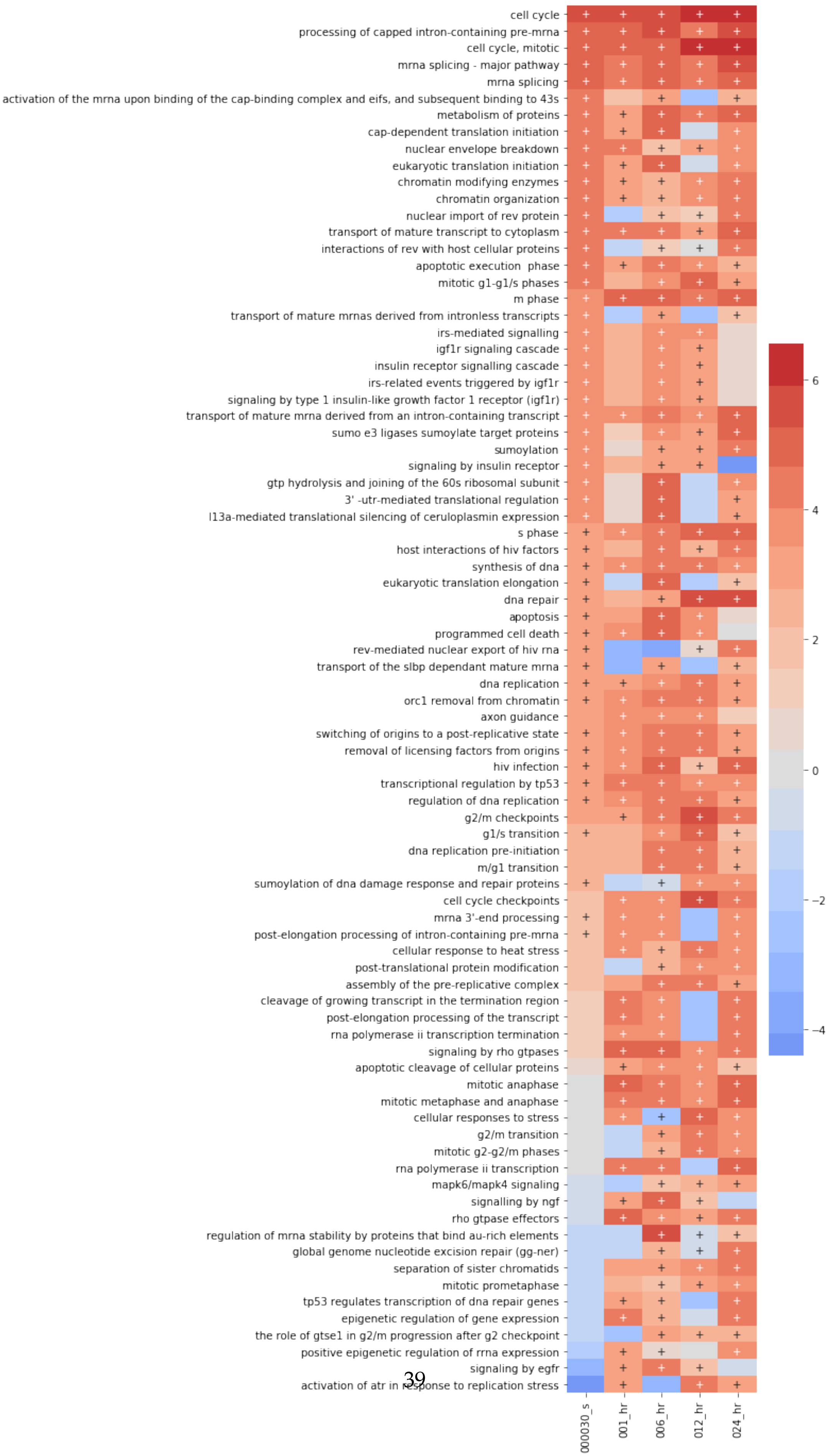
cell cycle
processing of capped intron-containing pre-mrna
cell cycle, mitotic
mrna splicing - major pathway
mrna splicing
activation of the mrna upon binding of the cap-binding complex and eifs, and subsequent binding to 43s
metabolism of proteins
cap-dependent translation initiation
nuclear envelope breakdown
eukaryotic translation initiation
chromatin modifying enzymes
chromatin organization
nuclear import of rev protein
transport of mature transcript to cytoplasm
interactions of rev with host cellular proteins
apoptotic execution phase
mitotic g1-g1/s phases
m phase
transport of mature mrnas derived from intronless transcripts
irs-mediated signalling
igf1r signaling cascade
insulin receptor signalling cascade
irs-related events triggered by igf1r
signaling by type 1 insulin-like growth factor 1 receptor (igf1r)
transport of mature mrna derived from an intron-containing transcript
sumo e3 ligases sumoylate target proteins
sumoylation
signaling by insulin receptor
gtp hydrolysis and joining of the 60s ribosomal subunit
3' -utr-mediated translational regulation
l13a-mediated translational silencing of ceruloplasmin expression
s phase
host interactions of hiv factors
synthesis of dna
eukaryotic translation elongation
dna repair
apoptosis
programmed cell death
rev-mediated nuclear export of hiv rna
transport of the slbp dependant mature mrna
dna replication
orc1 removal from chromatin
axon guidance
switching of origins to a post-replicative state
removal of licensing factors from origins
hiv infection
transcriptional regulation by tp53
regulation of dna replication
g2/m checkpoints
g1/s transition
dna replication pre-initiation
m/g1 transition
sumoylation of dna damage response and repair proteins
cell cycle checkpoints
mrna 3'-end processing
post-elongation processing of intron-containing pre-mrna
cellular response to heat stress
post-translational protein modification
assembly of the pre-replicative complex
cleavage of growing transcript in the termination region
post-elongation processing of the transcript
rna polymerase ii transcription termination
signaling by rho gtpases
apoptotic cleavage of cellular proteins
mitotic anaphase
mitotic metaphase and anaphase
cellular responses to stress
g2/m transition
mitotic g2-g2/m phases
rna polymerase ii transcription
mapk6/mapk4 signaling
signalling by ngf
rho gtpase effectors
regulation of mrna stability by proteins that bind au-rich elements
global genome nucleotide excision repair (gg-ner)
separation of sister chromatids
mitotic prometaphase
tp53 regulates transcription of dna repair genes
epigenetic regulation of gene expression
the role of gtse1 in g2/m progression after g2 checkpoint
positive epigenetic regulation of rrna expression
signaling by egfr
activation of atr in response to replication stress
000030_s
001_hr
006_hr
012_hr
024_hr
6
4
2
0
−2
−4

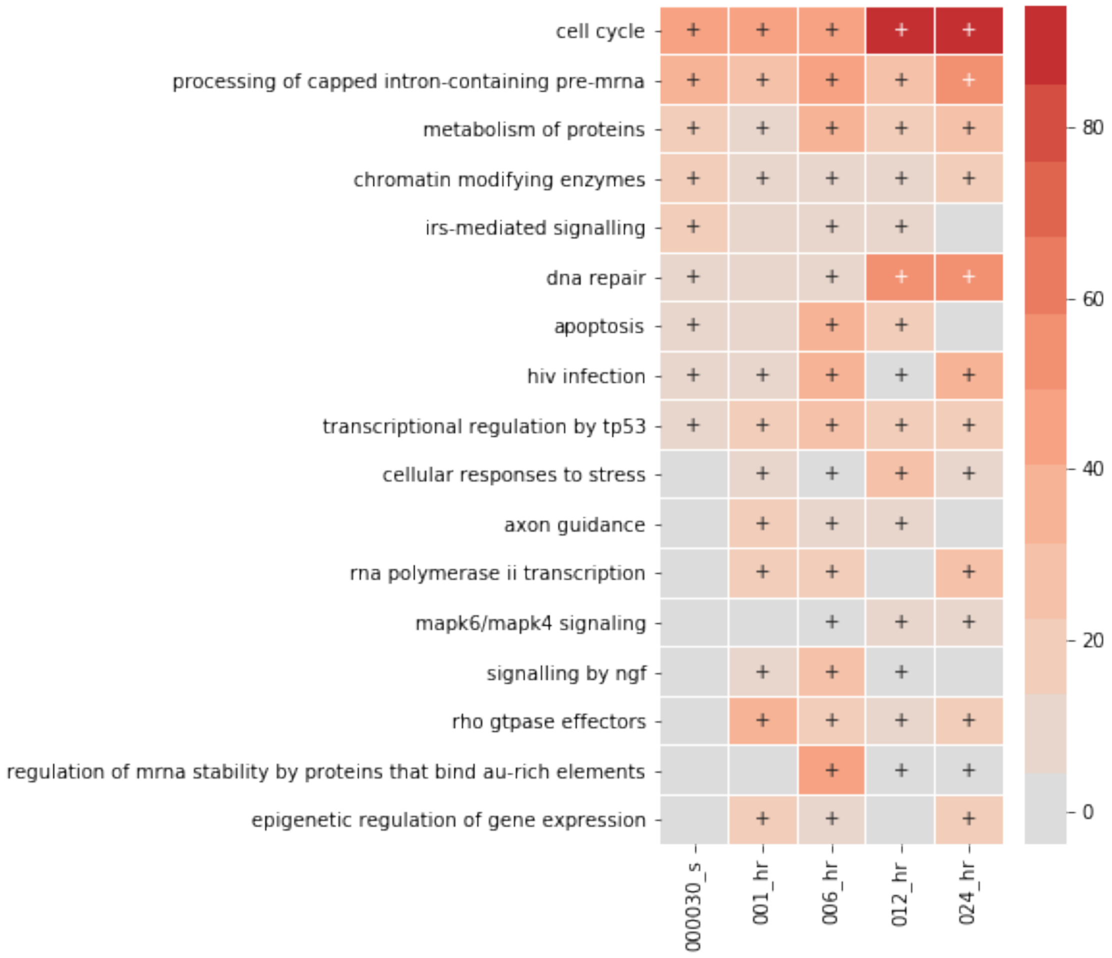


[34]:
```
# note that we can still get back terms that were compressed
ph_silac_up_copy.show_terms_below(
    'cell cycle',
    threshold=.7,
    remove_subset=True
).heatmap(
    figsize=(3, 12),
    cluster_by_set=False,
    linewidths=0.01,
);
```

```
Number of rows went from 83 to 17
```

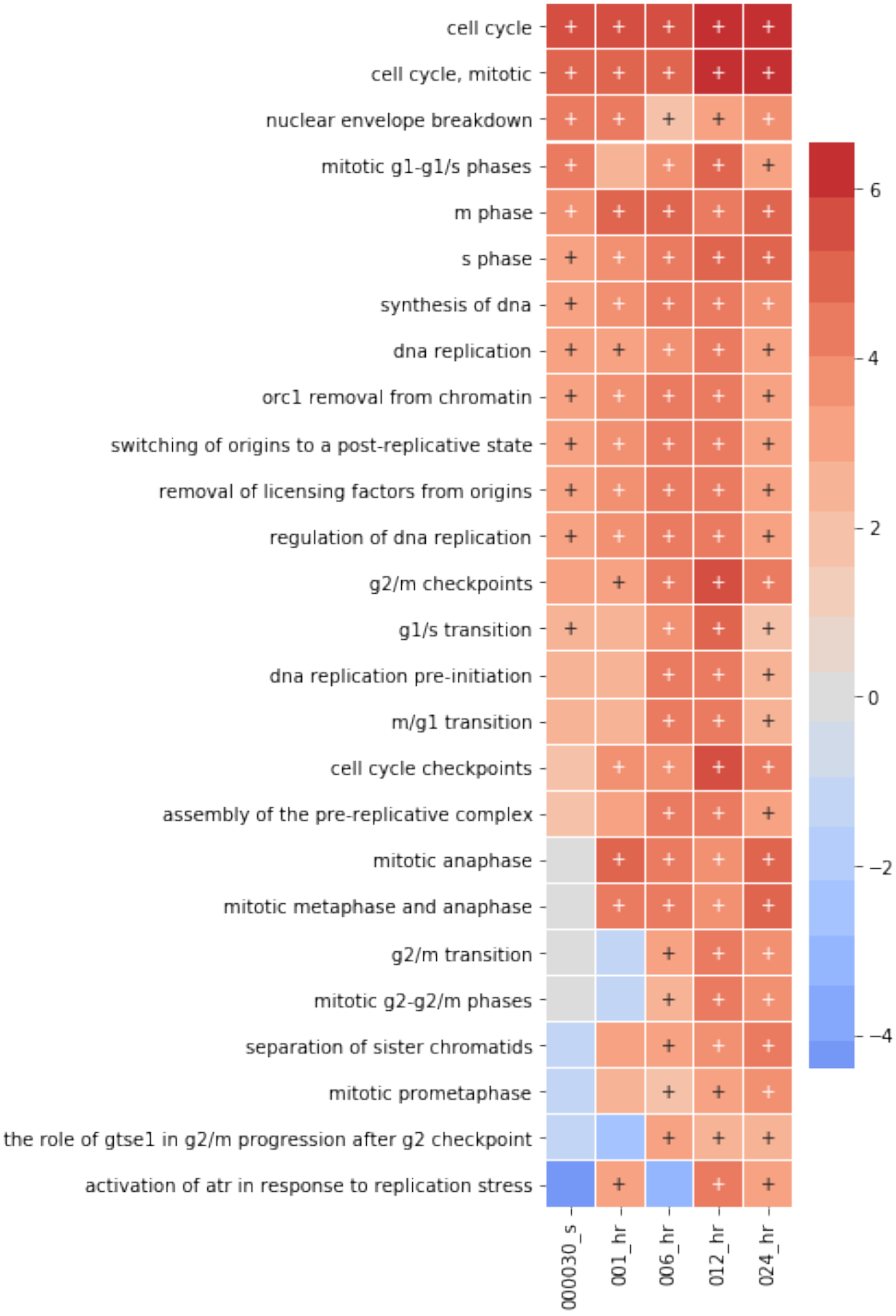
cell cycle
cell cycle, mitotic
nuclear envelope breakdown
mitotic g1-g1/s phases
m phase
s phase
synthesis of dna
dna replication
orc1 removal from chromatin
switching of origins to a post-replicative state
removal of licensing factors from origins
regulation of dna replication
g2/m checkpoints
g1/s transition
dna replication pre-initiation
m/g1 transition
cell cycle checkpoints
assembly of the pre-replicative complex
mitotic anaphase
mitotic metaphase and anaphase
g2/m transition
mitotic g2-g2/m phases
separation of sister chromatids
mitotic prometaphase
the role of gtse1 in g2/m progression after g2 checkpoint
activation of atr in response to replication stress
000030_s
001_hr
006_hr
012_hr
024_hr
6
4
2
0
−2
−4

[35]:
```
term_net, mol_net = create_subnetwork(
    ph_silac_up,
    network=network,
    save_name='ph_silac_ASN',
    use_cytoscape=False,
    use_fdr=True, use_threshold=True, min_edges=25
)
for i in ph_silac_up.term_name.unique():
    if i not in term_net.nodes:
        print(i)
```

```
Creating ontology network
```

[36]:
```
nx.set_node_attributes(term_net, 'white', 'color')
vis.draw_cyjs(term_net, layout='cose-bilkent', spacingFactor=1.4)
```

```
<IPython.core.display.HTML object>
```

##### 3.1.5 Exploring terms outside canonical

Since enrichment analysis is performed over all terms in a gene set, some of the terms might not make sense in certain context. This could be due to genes being classified in multiple sets, bad assignment, or possible new biology. The time it takes to explore each term in understanding `why` it is enriched can be time consuming, which is perhaps why people disregard terms frequently. We decided to use MAGINE to explore one of these terms. `HIV infection`

[37]:
```
reactome_only.sig.show_terms_below(
    'hiv infection',
    level='dataframe',
    threshold=.25,
    remove_subset=False
).heatmap(
    figsize=(6,16),
    linewidths=0.05,# cluster_by_set=True,
);
```

```
Number of rows went from 581 to 73
```

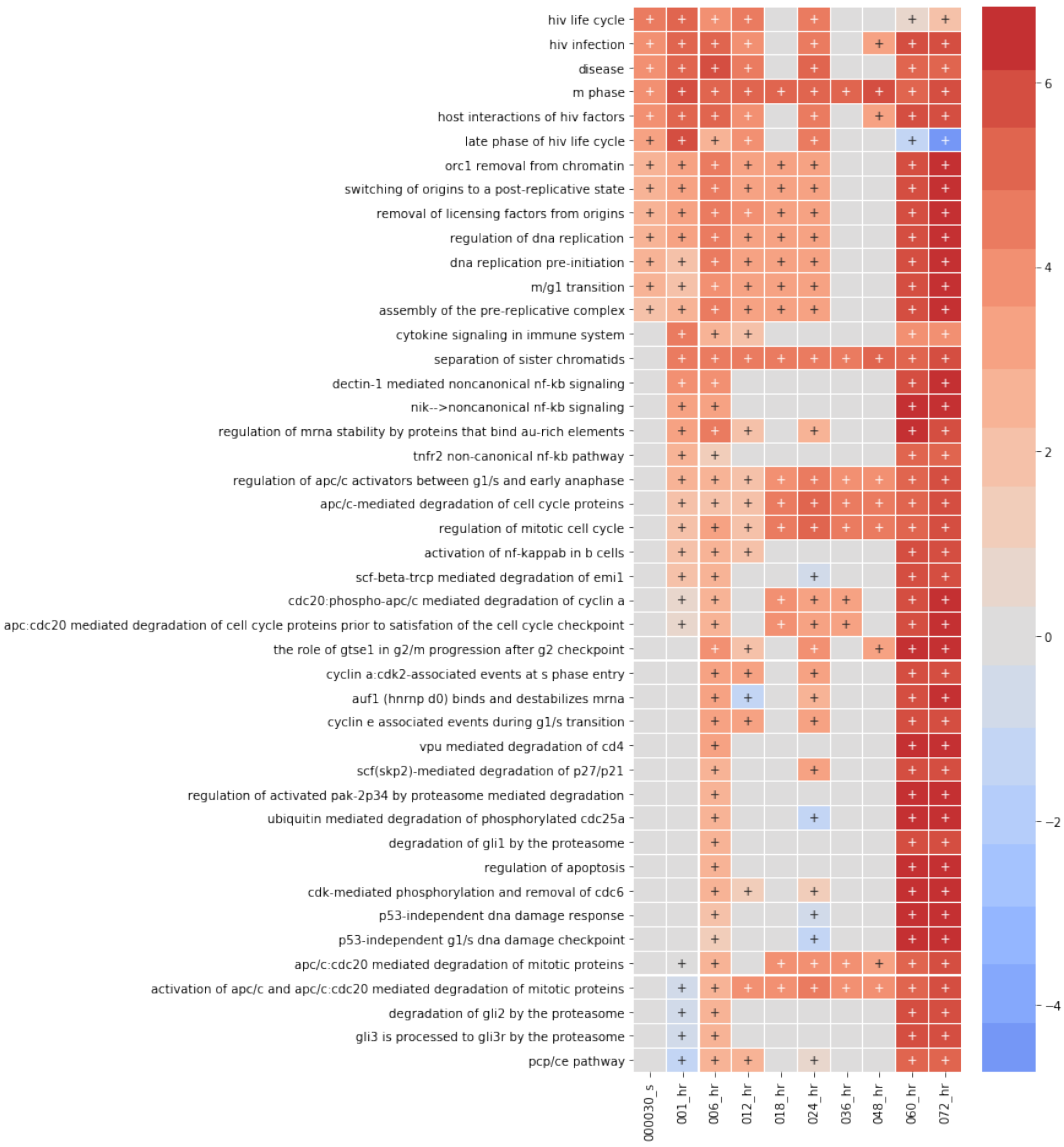


[38]:
```python
reactome_only.sig.find_similar_terms('hiv infection', level='dataframe',␣
↪remove_subset=False )
```

[38]:

| | term_name | similarity_score |
|---|---|---|
| 37 | host interactions of hiv factors | 0.70 |
| 21 | infectious disease | 0.68 |
| 17 | hiv life cycle | 0.57 |
| 46 | late phase of hiv life cycle | 0.49 |
| 27 | disease | 0.46 |
| 282 | pcp/ce pathway | 0.30 |
| 112 | cytokine signaling in immune system | 0.29 |
| 61 | m phase | 0.28 |

| | | |
|---|---|---|
| 167 | scf-beta-trcp mediated degradation of emi1 | 0.28 |
| 234 | gli3 is processed to gli3r by the proteasome | 0.27 |
| 233 | degradation of gli2 by the proteasome | 0.27 |
| 214 | activation of nf-kappab in b cells | 0.27 |
| 121 | nik-->noncanonical nf-kb signaling | 0.27 |
| 122 | dectin-1 mediated noncanonical nf-kb signaling | 0.27 |
| 134 | tnfr2 non-canonical nf-kb pathway | 0.27 |
| 223 | cyclin e associated events during g1/s transition | 0.27 |
| 254 | cdk-mediated phosphorylation and removal of cdc6 | 0.26 |
| 240 | scf(skp2)-mediated degradation of p27/p21 | 0.26 |
| 207 | cyclin a:cdk2-associated events at s phase entry | 0.26 |
| 274 | auf1 (hnrnp d0) binds and destabilizes mrna | 0.26 |
| 215 | vpu mediated degradation of cd4 | 0.26 |
| 194 | assembly of the pre-replicative complex | 0.26 |
| 270 | ubiquitin mediated degradation of phosphorylat... | 0.26 |
| 268 | p53-independent g1/s dna damage checkpoint | 0.26 |
| 265 | p53-independent dna damage response | 0.26 |
| 224 | cdc20:phospho-apc/c mediated degradation of cy... | 0.26 |
| 227 | apc:cdc20 mediated degradation of cell cycle p... | 0.26 |
| 129 | regulation of mrna stability by proteins that ... | 0.26 |
| 228 | apc/c:cdc20 mediated degradation of mitotic pr... | 0.26 |
| 192 | switching of origins to a post-replicative state | 0.26 |
| .. | ... | ... |
| 389 | mitochondrial translation initiation | 0.00 |
| 418 | clearance of nuclear envelope membranes from c... | 0.00 |
| 390 | mitochondrial translation termination | 0.00 |
| 153 | copi-mediated anterograde transport | 0.00 |
| 155 | apoptotic cleavage of cellular proteins | 0.00 |
| 394 | trna aminoacylation | 0.00 |
| 395 | cytosolic trna aminoacylation | 0.00 |
| 396 | rna polymerase iii transcription initiation fr... | 0.00 |
| 397 | rna polymerase iii transcription initiation fr... | 0.00 |
| 398 | deadenylation-dependent mrna decay | 0.00 |
| 399 | map2k and mapk activation | 0.00 |
| 400 | rna polymerase iii transcription initiation | 0.00 |
| 401 | transcriptional activation of mitochondrial bi... | 0.00 |
| 402 | rna polymerase iii abortive and retractive ini... | 0.00 |
| 403 | rna polymerase iii transcription | 0.00 |
| 404 | hdr through mmej (alt-nhej) | 0.00 |
| 405 | semet incorporation into proteins | 0.00 |
| 406 | tp53 regulates transcription of genes involved... | 0.00 |
| 407 | hdl-mediated lipid transport | 0.00 |
| 408 | growth hormone receptor signaling | 0.00 |
| 409 | pyrimidine biosynthesis | 0.00 |
| 410 | pyrimidine metabolism | 0.00 |
| 411 | initiation of nuclear envelope reformation | 0.00 |
| 412 | nuclear envelope reassembly | 0.00 |

```
413   purine ribonucleoside monophosphate biosynthesis              0.00
414  fatty acid, triacylglycerol, and ketone body m...              0.00
415                        signaling by robo receptor              0.00
416                        rho gtpases activate rocks              0.00
417                                  purine metabolism              0.00
579         synthesis of pyrophosphates in the cytosol              0.00

[580 rows x 2 columns]
```

Since HIV infection hijacks dna repair and regulates cell cycle, lets check to see if that explains why HIV infection is enriched.

[39]:
```python
hits = [
    'cell cycle',
    'dna repair',
    'hiv infection'
]
subset = reactome_only.sig.loc[reactome_only.sig.term_name.isin(hits)].copy()

g_sets = [reactome_only.sig.term_to_genes(i) for i in hits]
create_venn3(*g_sets+hits);
plt.savefig('venn_hiv.png', dpi=450)
```

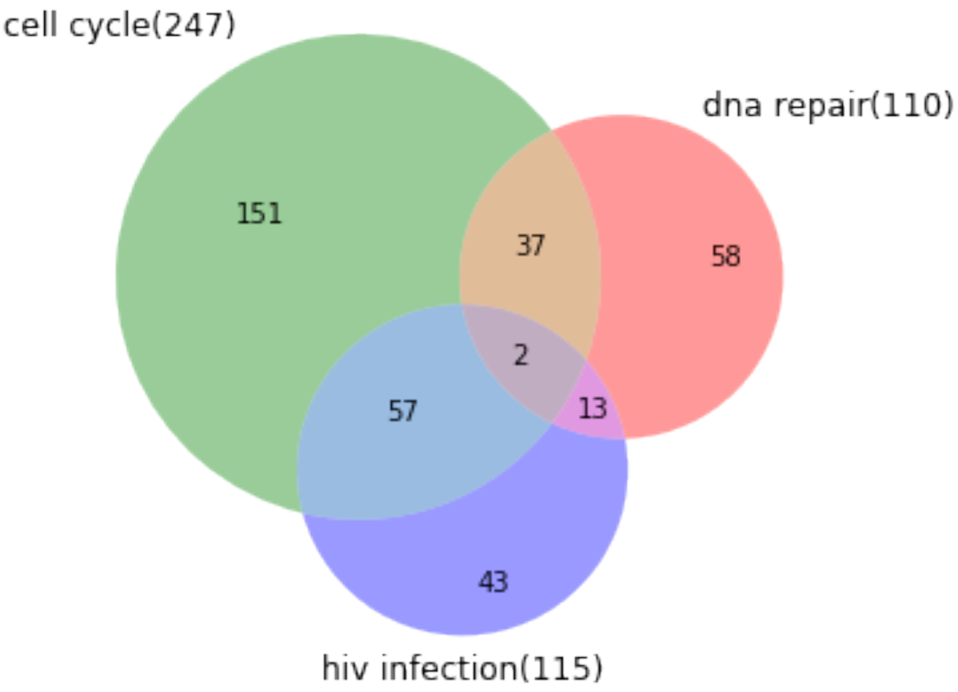


[40]:
```python
hiv_only= g_sets[-1]
hiv_only.difference_update(g_sets[0]) # remove dna repair genes
```

```python
hiv_only.difference_update(g_sets[1]) # removes cell cycle genes

exp_data.label_free.heatmap(
    hiv_only,
    cluster_row=True,
    subset_index='identifier',
    index='label',
    min_sig=1,
    linewidths=0.01
);
exp_data.ph_silac.heatmap(
    hiv_only,
    subset_index='identifier',
    index='label',
    cluster_row=True,
    rank_index=True,
    min_sig=1,
    figsize=(6, 16),
    linewidths=0.01
);
```

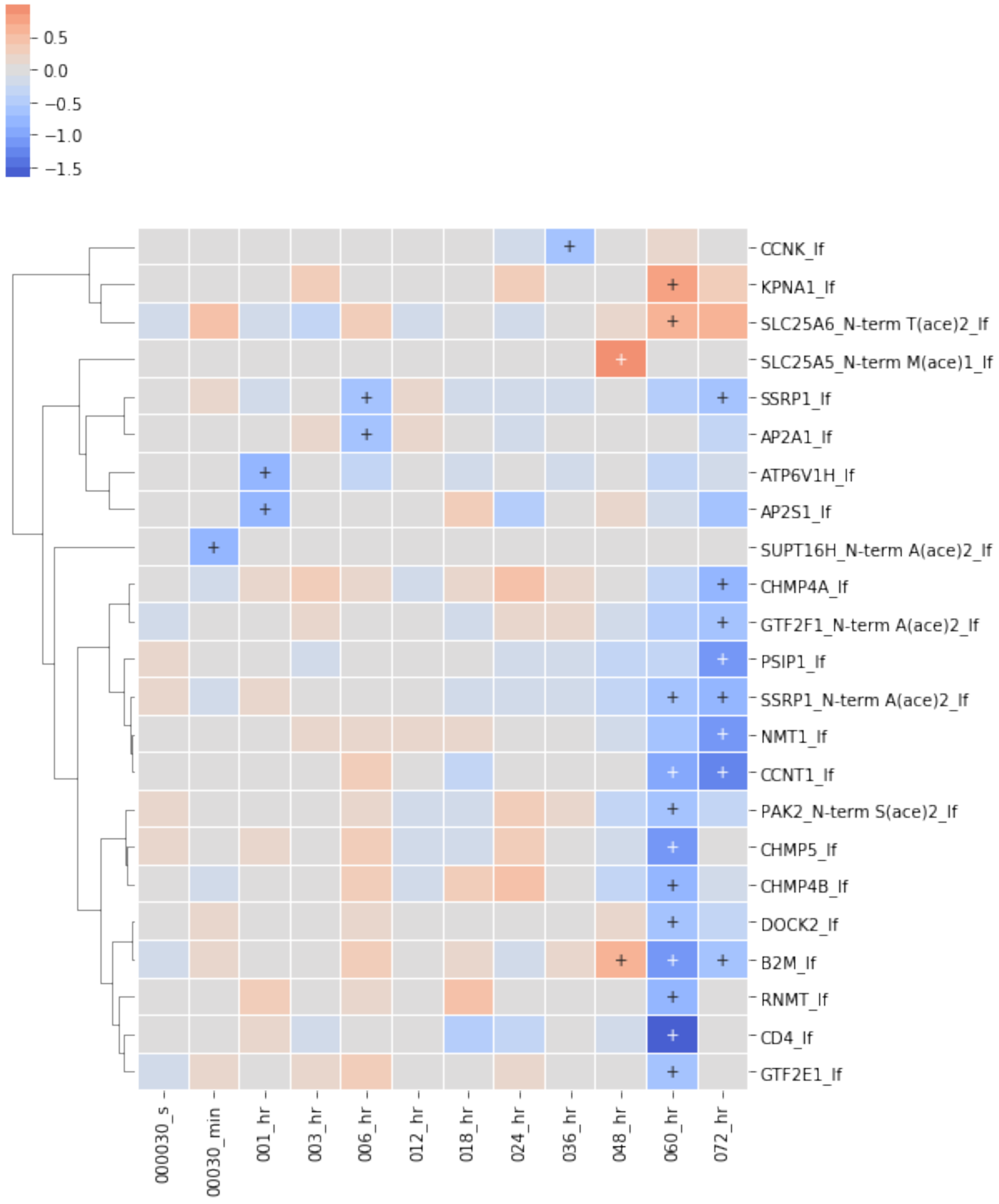

0.5
0.0
−0.5
−1.0
−1.5
CCNK_lf
KPNA1_lf
SLC25A6_N-term T(ace)2_lf
SLC25A5_N-term M(ace)1_lf
SSRP1_lf
AP2A1_lf
ATP6V1H_lf
AP2S1_lf
SUPT16H_N-term A(ace)2_lf
CHMP4A_lf
GTF2F1_N-term A(ace)2_lf
PSIP1_lf
SSRP1_N-term A(ace)2_lf
NMT1_lf
CCNT1_lf
PAK2_N-term S(ace)2_lf
CHMP5_lf
CHMP4B_lf
DOCK2_lf
B2M_lf
RNMT_lf
CD4_lf
GTF2E1_lf
000030_s
00030_min
001_hr
003_hr
006_hr
012_hr
018_hr
024_hr
036_hr
048_hr
060_hr
072_hr

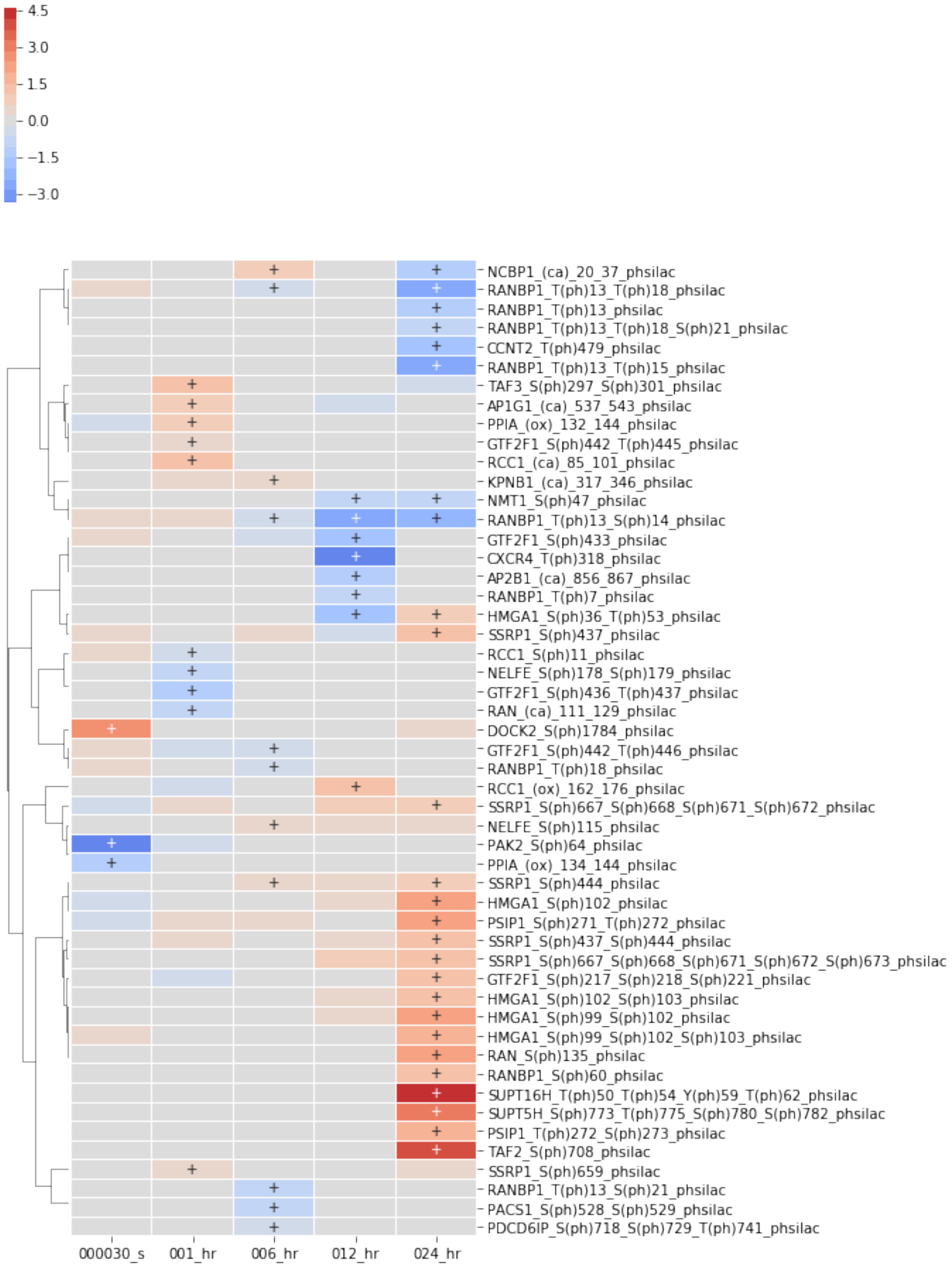


[41]:
```
term_net, mol_net = create_subnetwork(
    subset,
    network=network,
```

```
    save_name='hiv',
    use_cytoscape=False,
    use_fdr=True, use_threshold=True, min_edges=10
)
print(len(mol_net.edges))
print(len(mol_net.nodes))
```

```
Creating ontology network
2475
316
```

[42]:
```
vis.draw_cyjs(term_net)
```

```
<IPython.core.display.HTML object>
```

[43]:
```
term_to_gene = subset.term_to_genes_dict()

heatmap_by_terms(
    exp_data.label_free,
    convert_to_log=False,
    index='label',
    term_labels=list(term_to_gene.keys()),
    term_sets=list(term_to_gene.values()),
    div_colors=True,
    linewidths=0.01,
    min_sig=2,
    annotate_sig=True,
    cluster_col=False,
    figsize=(6,12),
    y_tick_labels=True
);
heatmap_by_terms(
    exp_data.ph_silac,
    convert_to_log=False,
    index='label',
    term_labels=list(term_to_gene.keys()),
    term_sets=list(term_to_gene.values()),
    div_colors=True,
    linewidths=0.01,
    min_sig=2,
    annotate_sig=True,
    cluster_col=False,
    figsize=(6,12),
    y_tick_labels=True
);
```

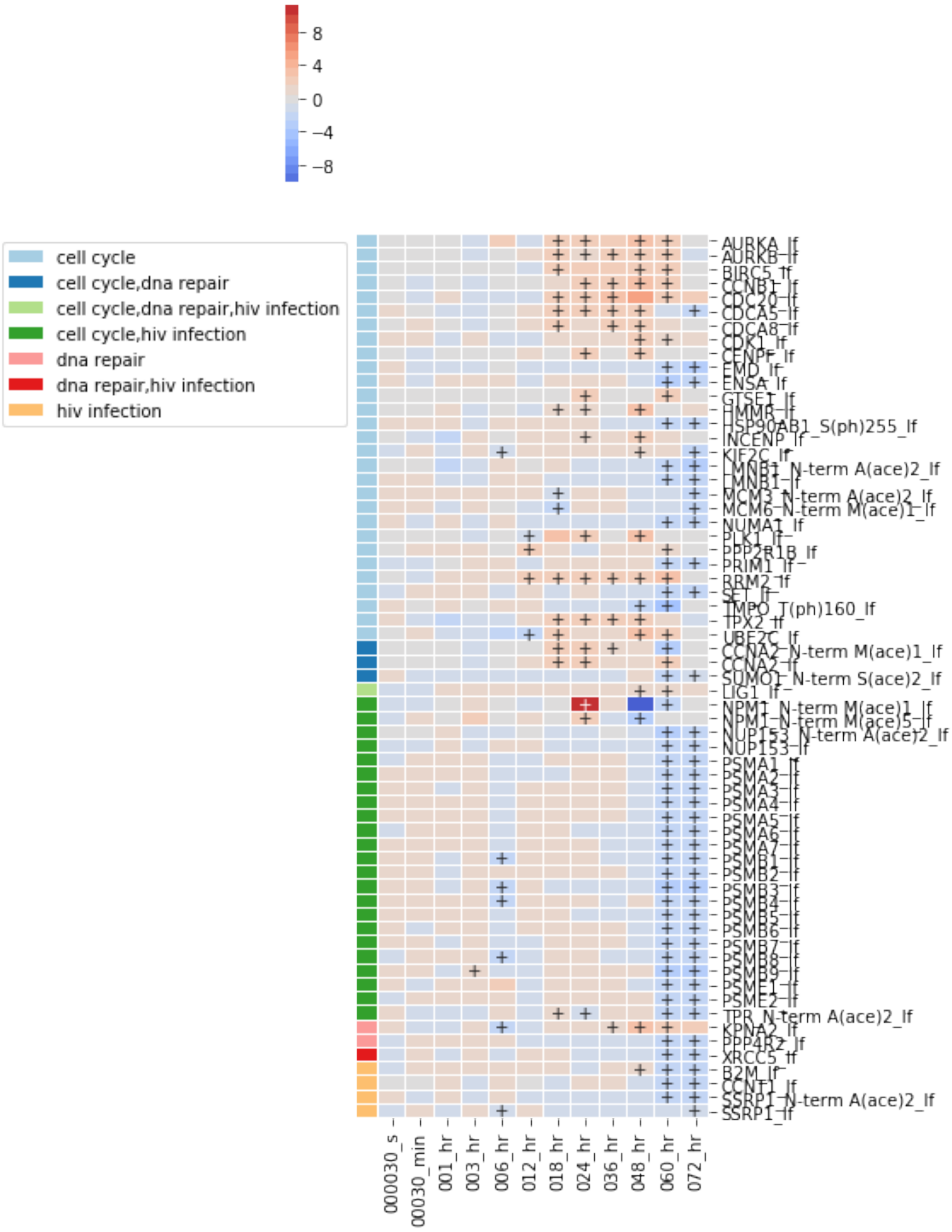
8
4
0
−4
−8
cell cycle
cell cycle,dna repair
cell cycle,dna repair,hiv infection
cell cycle,hiv infection
dna repair
dna repair,hiv infection
hiv infection
AURKA_lf
AURKB_lf
BIRC5_lf
CCNB1_lf
CDC20_lf
CDCA5_lf
CDCA8_lf
CDK1_lf
CENPF_lf
EMD_lf
ENSA_lf
GTSE1_lf
HMMR_lf
HSP90AB1_S(ph)255_lf
INCENP_lf
KIF2C_lf
LMNB1_N-term A(ace)2_lf
LMNB1_lf
MCM3_N-term A(ace)2_lf
MCM6_N-term M(ace)1_lf
NUMA1_lf
PLK1_lf
PPP2R1B_lf
PRIM1_lf
RRM2_lf
SET_lf
TMPO_T(ph)160_lf
TPX2_lf
UBE2C_lf
CCNA2_N-term M(ace)1_lf
CCNA2_lf
SUMO1_N-term S(ace)2_lf
LIG1_lf
NPM1_N-term M(ace)1_lf
NPM1_N-term M(ace)5_lf
NUP153_N-term A(ace)2_lf
NUP153_lf
PSMA1_lf
PSMA2_lf
PSMA3_lf
PSMA4_lf
PSMA5_lf
PSMA6_lf
PSMA7_lf
PSMB1_lf
PSMB2_lf
PSMB3_lf
PSMB4_lf
PSMB5_lf
PSMB6_lf
PSMB7_lf
PSMB8_lf
PSMB9_lf
PSME1_lf
PSME2_lf
TPR_N-term A(ace)2_lf
KPNA2_lf
PPP4R2_lf
XRCC5_lf
B2M_lf
CCNT1_lf
SSRP1_N-term A(ace)2_lf
SSRP1_lf
000030_s
00030_min
001_hr
003_hr
006_hr
012_hr
018_hr
024_hr
036_hr
048_hr
060_hr
072_hr

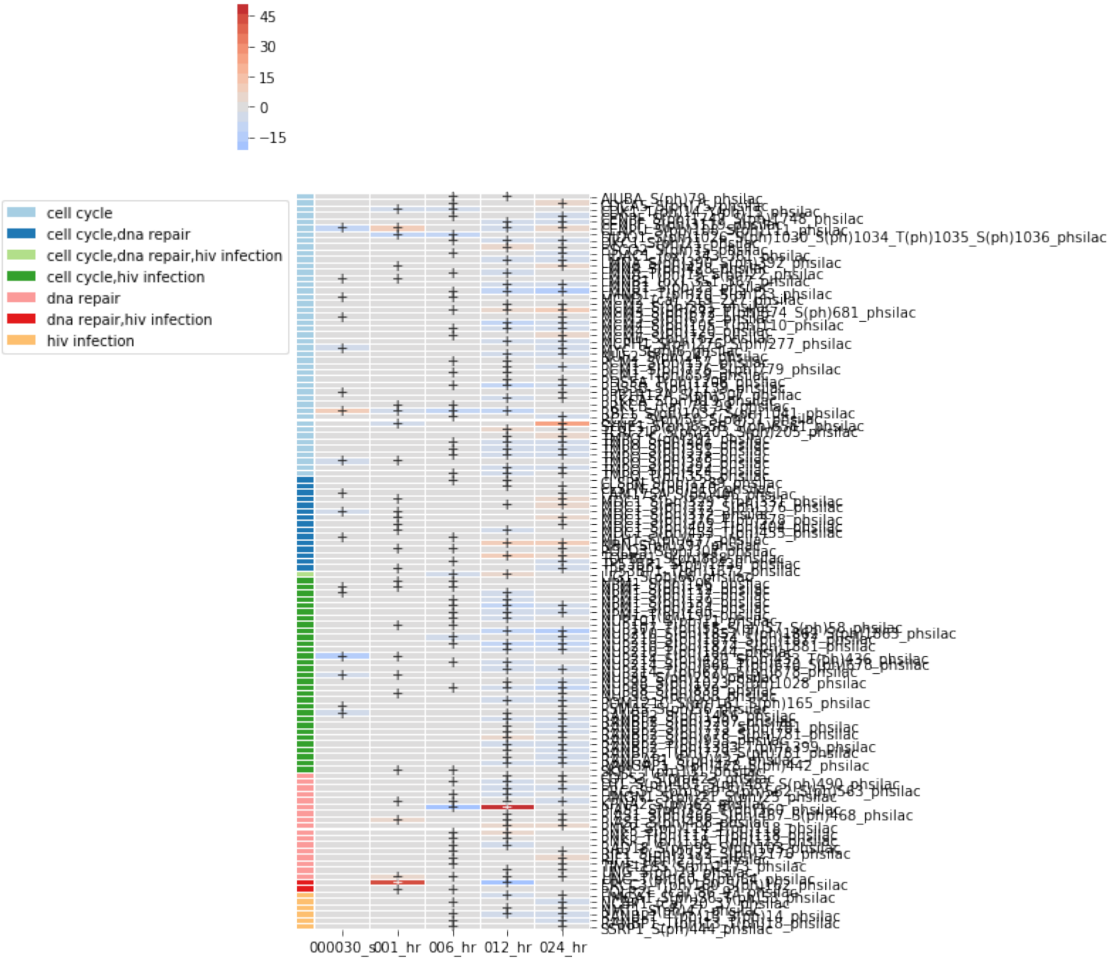


##### 3.1.6 Exploring a subset of terms based on a keyword

[44]:
```
# Filter by terms
damage_terms = enrichment_array.sig.filter_based_on_words(['damage'])
print(len(damage_terms.all_genes_from_df()))
damage_terms.head(10)
damage_terms_all = damage_terms.filter_multi(category='ph_silac_both')
first_damage = damage_terms_all.filter_multi(sample_id='000030_s')
display(first_damage[cols].head(20))

fig = first_damage.heatmap(
    convert_to_log=False,
    cluster_by_set=False,
    annotate_sig=True,
    div_colors=True,
    linewidths=.01,
    num_colors=21,
```

```
    figsize=(3,5)
);
fig.savefig('dna_damage_terms_30s.png', dpi=300, bbox_inches='tight')

damage_terms.require_n_sig(inplace=True, columns='sample_id', n_sig=2)
damage_terms.remove_redundant(inplace=True, threshold=.5, level='sample')
fig = damage_terms.heatmap(
    convert_to_log=False,
    cluster_by_set=False,
    annotate_sig=True,
    div_colors=True,
    linewidths=.01,
    num_colors=21,
    figsize=(4,8)
);
fig.savefig('dna_damage_terms_all.png', bbox_inches='tight')

dna_gene_df = exp_data.subset(first_damage.all_genes_from_df())
dna_gene_df = dna_gene_df.loc[dna_gene_df.sample_id.isin(['000030_s'])]

dna_gene_df.sig.heatmap(
    index='label',
    rank_index=False,
    convert_to_log=True,
    annotate_sig=True,
    div_colors=True,
    linewidths=.01,
    num_colors=21,
    figsize=(2,22)
);
```

```
478

                                               term_name  rank  \
3504789        dna damage induced protein phosphorylation    12
3504932  intrinsic apoptotic signaling pathway in respo...   155
3504975  histone h4 acetylation involved in response to...   198
3504976           cellular response to dna damage stimulus   199
3504977  histone h3-k56 acetylation in response to dna ...   200
3504979  regulation of transcription from rna polymeras...   202
3504989      signal transduction in response to dna damage   212
3505010                             dna damage checkpoint    233
3505029     telomere maintenance in response to dna damage   252
3505202  dna damage response, signal transduction by p5...   425
3505281  negative regulation of mitotic dna damage chec...   504
3505631  regulation of telomere maintenance in response...   854
3511095                               damaged dna binding    74
3516776  sumoylation of dna damage response and repair ...    36
```

```
3516900  dna damage/telomere stress induced senescence_...   160

        combined_score  adj_p_value  n_genes sample_id      category
3504789          73.18     5.42e-05       36  000030_s  ph_silac_both
3504932          29.22     2.92e-03       18  000030_s  ph_silac_both
3504975          25.87     5.85e-03       17  000030_s  ph_silac_both
3504976          25.87     4.08e-03       16  000030_s  ph_silac_both
3504977          25.85     4.08e-03       16  000030_s  ph_silac_both
3504979          25.76     4.26e-03       16  000030_s  ph_silac_both
3504989          24.76     5.45e-03       16  000030_s  ph_silac_both
3505010          23.40     8.45e-03       16  000030_s  ph_silac_both
3505029          21.49     1.27e-02       16  000030_s  ph_silac_both
3505202          12.66     3.12e-02        9  000030_s  ph_silac_both
3505281          11.45     2.22e-02        6  000030_s  ph_silac_both
3505631           0.10     4.32e-02        4  000030_s  ph_silac_both
3511095          17.49     9.51e-03       29  000030_s  ph_silac_both
3516776          21.40     6.71e-05       12  000030_s  ph_silac_both
3516900           0.86     4.80e-02        6  000030_s  ph_silac_both


Number of rows went from 34 to 23
```

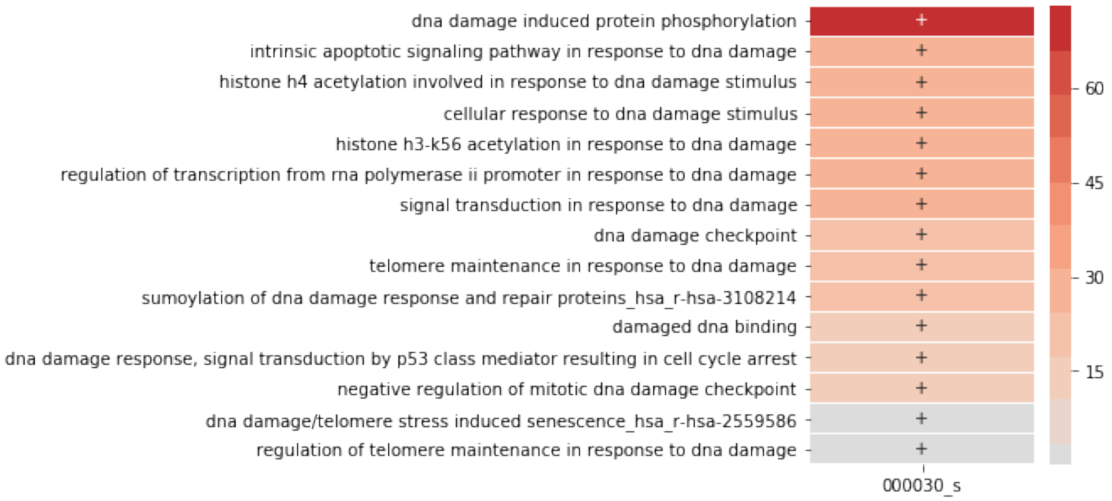

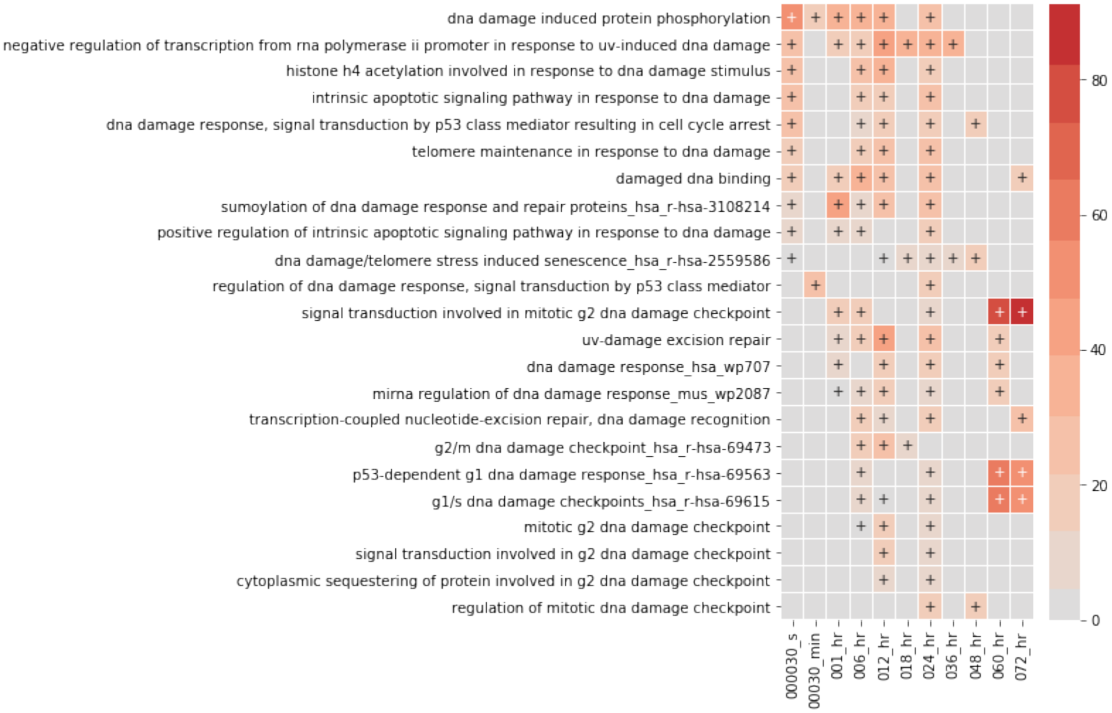
dna damage induced protein phosphorylation
negative regulation of transcription from rna polymerase ii promoter in response to uv-induced dna damage
histone h4 acetylation involved in response to dna damage stimulus
intrinsic apoptotic signaling pathway in response to dna damage
dna damage response, signal transduction by p53 class mediator resulting in cell cycle arrest
telomere maintenance in response to dna damage
damaged dna binding
sumoylation of dna damage response and repair proteins_hsa_r-hsa-3108214
positive regulation of intrinsic apoptotic signaling pathway in response to dna damage
dna damage/telomere stress induced senescence_hsa_r-hsa-2559586
regulation of dna damage response, signal transduction by p53 class mediator
signal transduction involved in mitotic g2 dna damage checkpoint
uv-damage excision repair
dna damage response_hsa_wp707
mirna regulation of dna damage response_mus_wp2087
transcription-coupled nucleotide-excision repair, dna damage recognition
g2/m dna damage checkpoint_hsa_r-hsa-69473
p53-dependent g1 dna damage response_hsa_r-hsa-69563
g1/s dna damage checkpoints_hsa_r-hsa-69615
mitotic g2 dna damage checkpoint
signal transduction involved in g2 dna damage checkpoint
cytoplasmic sequestering of protein involved in g2 dna damage checkpoint
regulation of mitotic dna damage checkpoint
000030_s
00030_min
001_hr
006_hr
012_hr
018_hr
024_hr
036_hr
048_hr
060_hr
072_hr
80
60
40
20
0

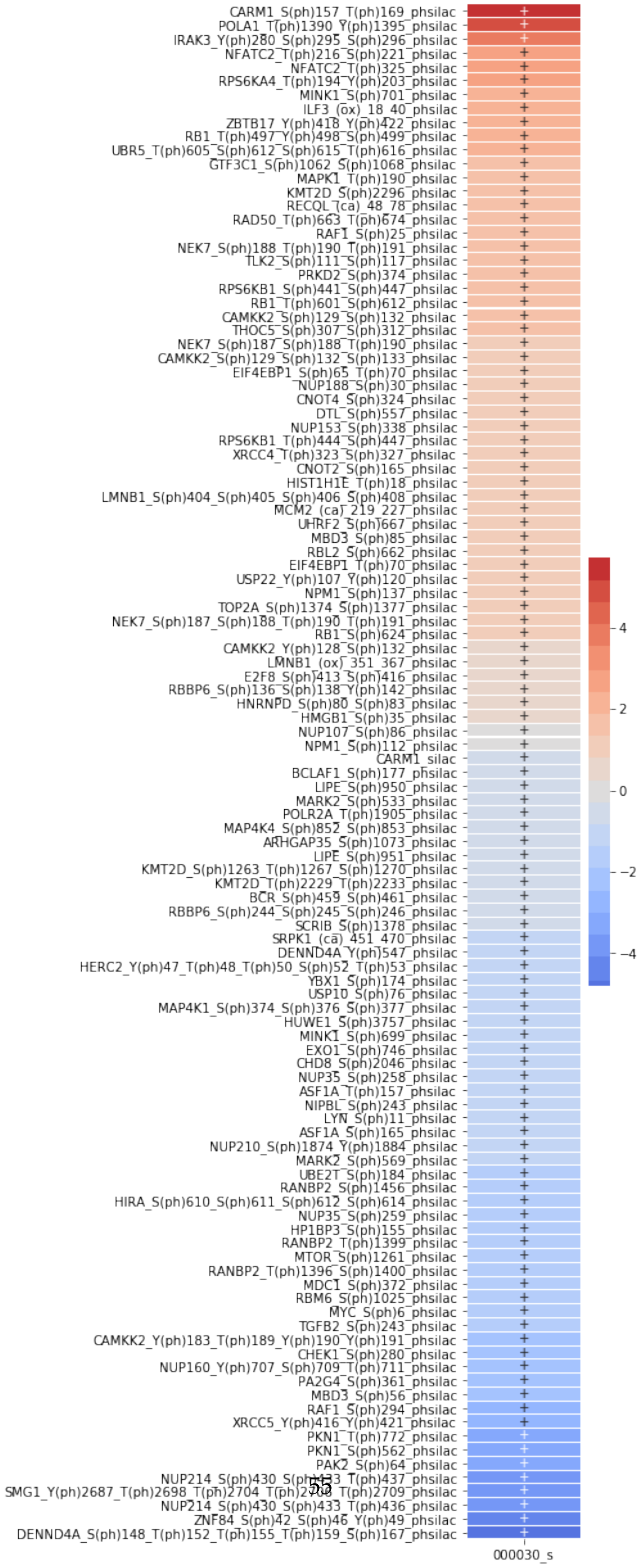
CARM1_S(ph)157_T(ph)169_phsilac
POLA1_T(ph)1390_Y(ph)1395_phsilac
IRAK3_Y(ph)280_S(ph)295_S(ph)296_phsilac
NFATC2_T(ph)216_S(ph)221_phsilac
NFATC2_T(ph)325_phsilac
RPS6KA4_T(ph)194_Y(ph)203_phsilac
MINK1_S(ph)701_phsilac
ILF3_(ox)_18_40_phsilac
ZBTB17_Y(ph)418_Y(ph)422_phsilac
RB1_T(ph)497_Y(ph)498_S(ph)499_phsilac
UBR5_T(ph)605_S(ph)612_S(ph)615_T(ph)616_phsilac
GTF3C1_S(ph)1062_S(ph)1068_phsilac
MAPK1_T(ph)190_phsilac
KMT2D_S(ph)2296_phsilac
RECQL_(ca)_48_78_phsilac
RAD50_T(ph)663_T(ph)674_phsilac
RAF1_S(ph)25_phsilac
NEK7_S(ph)188_T(ph)190_T(ph)191_phsilac
TLK2_S(ph)111_S(ph)117_phsilac
PRKD2_S(ph)374_phsilac
RPS6KB1_S(ph)441_S(ph)447_phsilac
RB1_T(ph)601_S(ph)612_phsilac
CAMKK2_S(ph)129_S(ph)132_phsilac
THOC5_S(ph)307_S(ph)312_phsilac
NEK7_S(ph)187_S(ph)188_T(ph)190_phsilac
CAMKK2_S(ph)129_S(ph)132_S(ph)133_phsilac
EIF4EBP1_S(ph)65_T(ph)70_phsilac
NUP188_S(ph)30_phsilac
CNOT4_S(ph)324_phsilac
DTL_S(ph)557_phsilac
NUP153_S(ph)338_phsilac
RPS6KB1_T(ph)444_S(ph)447_phsilac
XRCC4_T(ph)323_S(ph)327_phsilac
CNOT2_S(ph)165_phsilac
HIST1H1E_T(ph)18_phsilac
LMNB1_S(ph)404_S(ph)405_S(ph)406_S(ph)408_phsilac
MCM2_(ca)_219_227_phsilac
UHRF2_S(ph)667_phsilac
MBD3_S(ph)85_phsilac
RBL2_S(ph)662_phsilac
EIF4EBP1_T(ph)70_phsilac
USP22_Y(ph)107_Y(ph)120_phsilac
NPM1_S(ph)137_phsilac
TOP2A_S(ph)1374_S(ph)1377_phsilac
NEK7_S(ph)187_S(ph)188_T(ph)190_T(ph)191_phsilac
RB1_S(ph)624_phsilac
CAMKK2_Y(ph)128_S(ph)132_phsilac
LMNB1_(ox)_351_367_phsilac
E2F8_S(ph)413_S(ph)416_phsilac
RBBP6_S(ph)136_S(ph)138_Y(ph)142_phsilac
HNRNPD_S(ph)80_S(ph)83_phsilac
HMGB1_S(ph)35_phsilac
NUP107_S(ph)86_phsilac
NPM1_S(ph)112_phsilac
CARM1_silac
BCLAF1_S(ph)177_phsilac
LIPE_S(ph)950_phsilac
MARK2_S(ph)533_phsilac
POLR2A_T(ph)1905_phsilac
MAP4K4_S(ph)852_S(ph)853_phsilac
ARHGAP35_S(ph)1073_phsilac
LIPE_S(ph)951_phsilac
KMT2D_S(ph)1263_T(ph)1267_S(ph)1270_phsilac
KMT2D_T(ph)2229_T(ph)2233_phsilac
BCR_S(ph)459_S(ph)461_phsilac
RBBP6_S(ph)244_S(ph)245_S(ph)246_phsilac
SCRIB_S(ph)1378_phsilac
SRPK1_(ca)_451_470_phsilac
DENND4A_Y(ph)547_phsilac
HERC2_Y(ph)47_T(ph)48_T(ph)50_S(ph)52_T(ph)53_phsilac
YBX1_S(ph)174_phsilac
USP10_S(ph)76_phsilac
MAP4K1_S(ph)374_S(ph)376_S(ph)377_phsilac
HUWE1_S(ph)3757_phsilac
MINK1_S(ph)699_phsilac
EXO1_S(ph)746_phsilac
CHD8_S(ph)2046_phsilac
NUP35_S(ph)258_phsilac
ASF1A_T(ph)157_phsilac
NIPBL_S(ph)243_phsilac
LYN_S(ph)11_phsilac
ASF1A_S(ph)165_phsilac
NUP210_S(ph)1874_Y(ph)1884_phsilac
MARK2_S(ph)569_phsilac
UBE2T_S(ph)184_phsilac
RANBP2_S(ph)1456_phsilac
HIRA_S(ph)610_S(ph)611_S(ph)612_S(ph)614_phsilac
NUP35_S(ph)259_phsilac
HP1BP3_S(ph)155_phsilac
RANBP2_T(ph)1399_phsilac
MTOR_S(ph)1261_phsilac
RANBP2_T(ph)1396_S(ph)1400_phsilac
MDC1_S(ph)372_phsilac
RBM6_S(ph)1025_phsilac
MYC_S(ph)6_phsilac
TGFB2_S(ph)243_phsilac
CAMKK2_Y(ph)183_T(ph)189_Y(ph)190_Y(ph)191_phsilac
CHEK1_S(ph)280_phsilac
NUP160_Y(ph)707_S(ph)709_T(ph)711_phsilac
PA2G4_S(ph)361_phsilac
MBD3_S(ph)56_phsilac
RAF1_S(ph)294_phsilac
XRCC5_Y(ph)416_Y(ph)421_phsilac
PKN1_T(ph)772_phsilac
PKN1_S(ph)562_phsilac
PAK2_S(ph)64_phsilac
NUP214_S(ph)430_S(ph)433_T(ph)437_phsilac
SMG1_Y(ph)2687_T(ph)2698_T(ph)2704_T(ph)2705_T(ph)2709_phsilac
NUP214_S(ph)430_S(ph)433_T(ph)436_phsilac
ZNF84_S(ph)42_S(ph)46_Y(ph)49_phsilac
DENND4A_S(ph)148_T(ph)152_T(ph)155_T(ph)159_S(ph)167_phsilac
4
2
0
−2
−4
000030_s

##### 3.1.7 Exploring species of interest

[45]:
```
g2_m = ['CDK1', 'CCNB1']
cdk1_inhibitors = ['GADD45A', 'GADD45B', 'GADD45G', 'CDKN1A']

exp_data.species.heatmap(
    g2_m,
    index='label',
    subset_index='identifier',
    figsize=(4,8),
    linewidths=0.01,
    cluster_row=True,
    min_sig=1
);

exp_data.species.heatmap(
    cdk1_inhibitors,
    index='label',
    subset_index='identifier',
    figsize=(4, 4),
    linewidths=0.01,
    cluster_row=True,
    min_sig=1
);
```

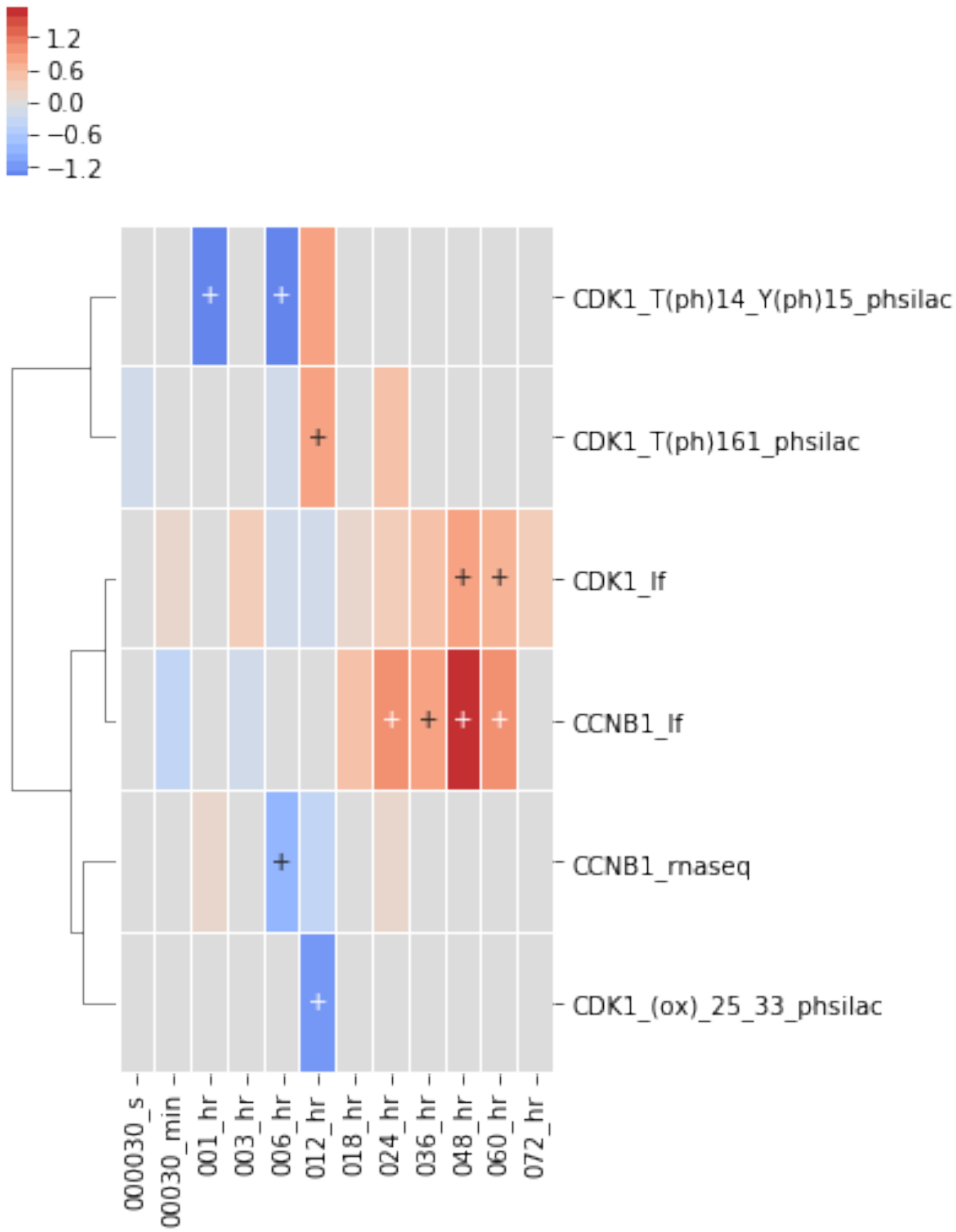

1.2
0.6
0.0
−0.6
−1.2
CDK1_T(ph)14_Y(ph)15_phsilac
CDK1_T(ph)161_phsilac
CDK1_lf
CCNB1_lf
CCNB1_rnaseq
CDK1_(ox)_25_33_phsilac
000030_s
00030_min
001_hr
003_hr
006_hr
012_hr
018_hr
024_hr
036_hr
048_hr
060_hr
072_hr

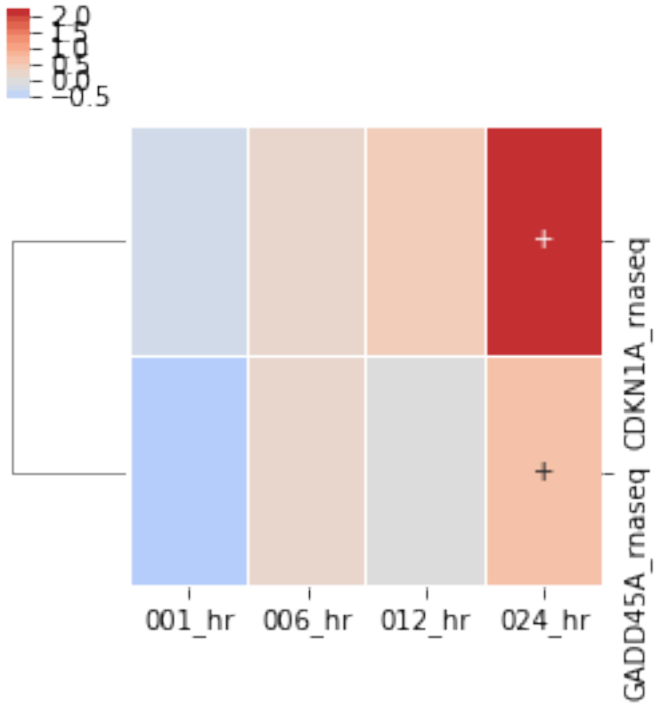


[46]:
```
expand_neigh = net_sub.expand_neighbors(
    network=None,
    nodes=g2_m,
    downstream=True,
    upstream=True,
    max_dist=1,
    include_only=reactome_only.filter_multi(category=['label_free_up']).
 ↪term_to_genes('cell cycle'),
    add_interconnecting_edges=False,
)

expand_neigh = net_sub.paths_between_two_lists(
    reactome_only.filter_multi(category=['ph_silac_up']).term_to_genes('dna␣
 ↪repair'),
    g2_m,
    max_length=3,
    include_only=reactome_only.filter_multi(category=['label_free_up']).
 ↪term_to_genes('cell cycle'),
    add_interconnecting_edges=True
)
print(len(expand_neigh.nodes))
print(len(expand_neigh.edges))
expand_neigh = utils.delete_disconnected_network(expand_neigh)
exp_data.label_free.heatmap(
```

```python
    expand_neigh.nodes,
    subset_index='identifier',
    index='label',
    rank_index=False,
    cluster_row=True,
    min_sig=3,
    annotate_sig=True,
    linewidths=0.01
);
```

```
Warning : 2 do not exist in graph
Removing from list
25
47
```

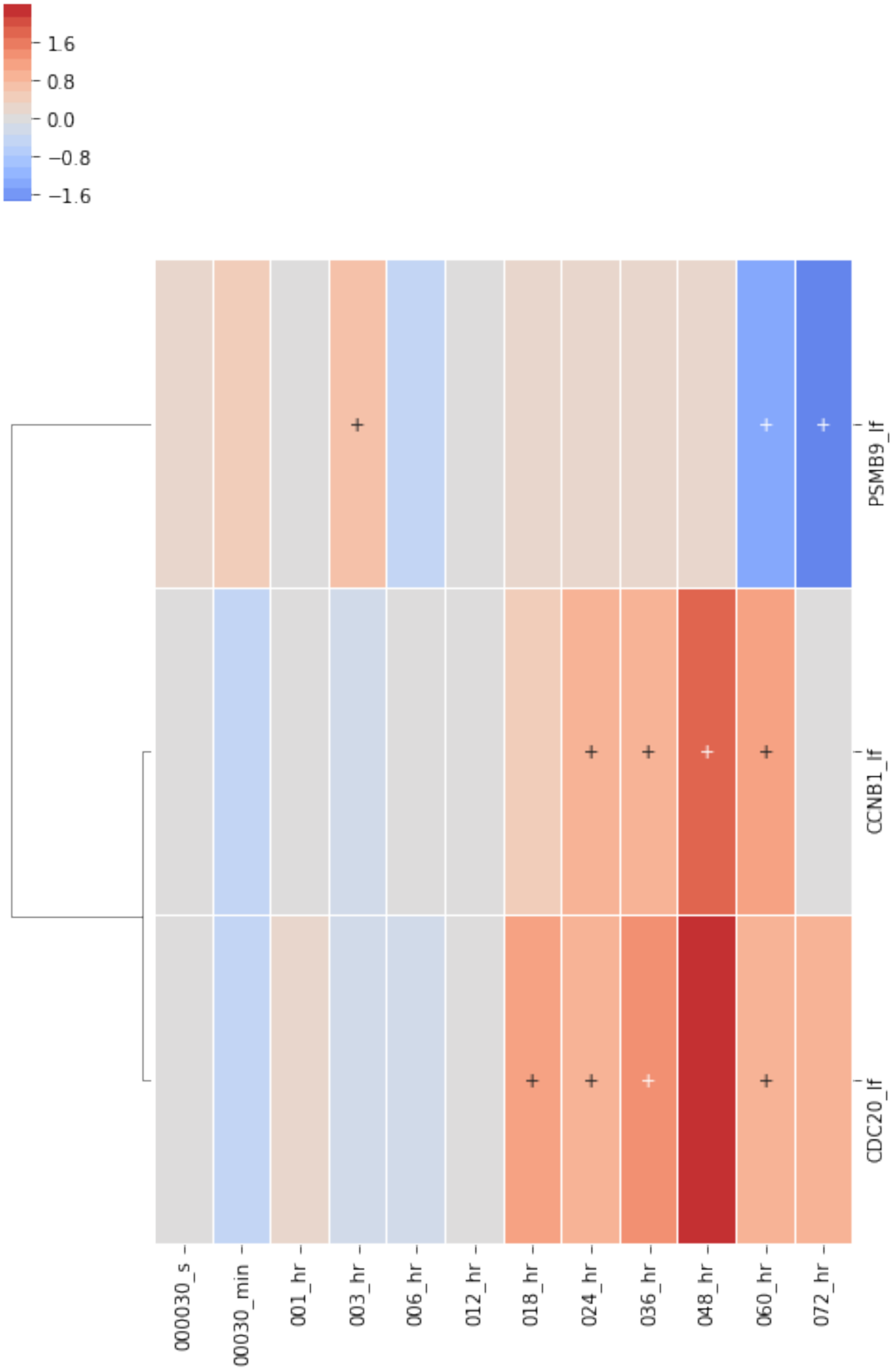
1.6
0.8
0.0
−0.8
−1.6
PSMB9_lf
CCNB1_lf
CDC20_lf
000030_s
00030_min
001_hr
003_hr
006_hr
012_hr
018_hr
024_hr
036_hr
048_hr
060_hr
072_hr

[47]:
```
#expand_neigh = utils.add_attribute_to_network(expand_neigh, g2_m, 'color',␣
↪'red', 'white')
vis.draw_cyjs(
    expand_neigh, layout='dagre',
    spacingFactor=1.,
    nodeRepulsion=100, gravity=.1,
    rankDir='TB', nodeSep=5, rankSep=25, ranker='longest-path'
)
```

```
<IPython.core.display.HTML object>
```

[48]:
```
expand_neigh = net_sub.paths_between_two_lists(
    reactome_only.filter_multi(category=['ph_silac_up']).term_to_genes('dna␣
↪repair'),
    g2_m,
    reverse=True,
    max_length=3,
    include_only=exp_data.species.sig,
    add_interconnecting_edges=False
)

expand_neigh = utils.delete_disconnected_network(expand_neigh)

print(len(expand_neigh.nodes))
print(len(expand_neigh.edges))
```

```
Warning : 2 do not exist in graph
Removing from list
29
38
```

[49]:
```
vis.draw_cyjs(
    expand_neigh, layout='dagre',
    spacingFactor=1.,
    nodeRepulsion=100, gravity=.1,
    rankDir='TB', nodeSep=5, rankSep=25, ranker='longest-path'
)
```

```
<IPython.core.display.HTML object>
```

[50]:
```
exp_data.ph_silac.heatmap(
    expand_neigh.nodes,
    subset_index='identifier',
```

```
    index='label',
    rank_index=False,
    cluster_row=True,
    min_sig=2,
    linewidths=0.01
);

exp_data.label_free.heatmap(
    expand_neigh.nodes,
    subset_index='identifier',
    index='label',
    rank_index=False,
    cluster_row=True,
    min_sig=2,
    linewidths=0.01
);
```

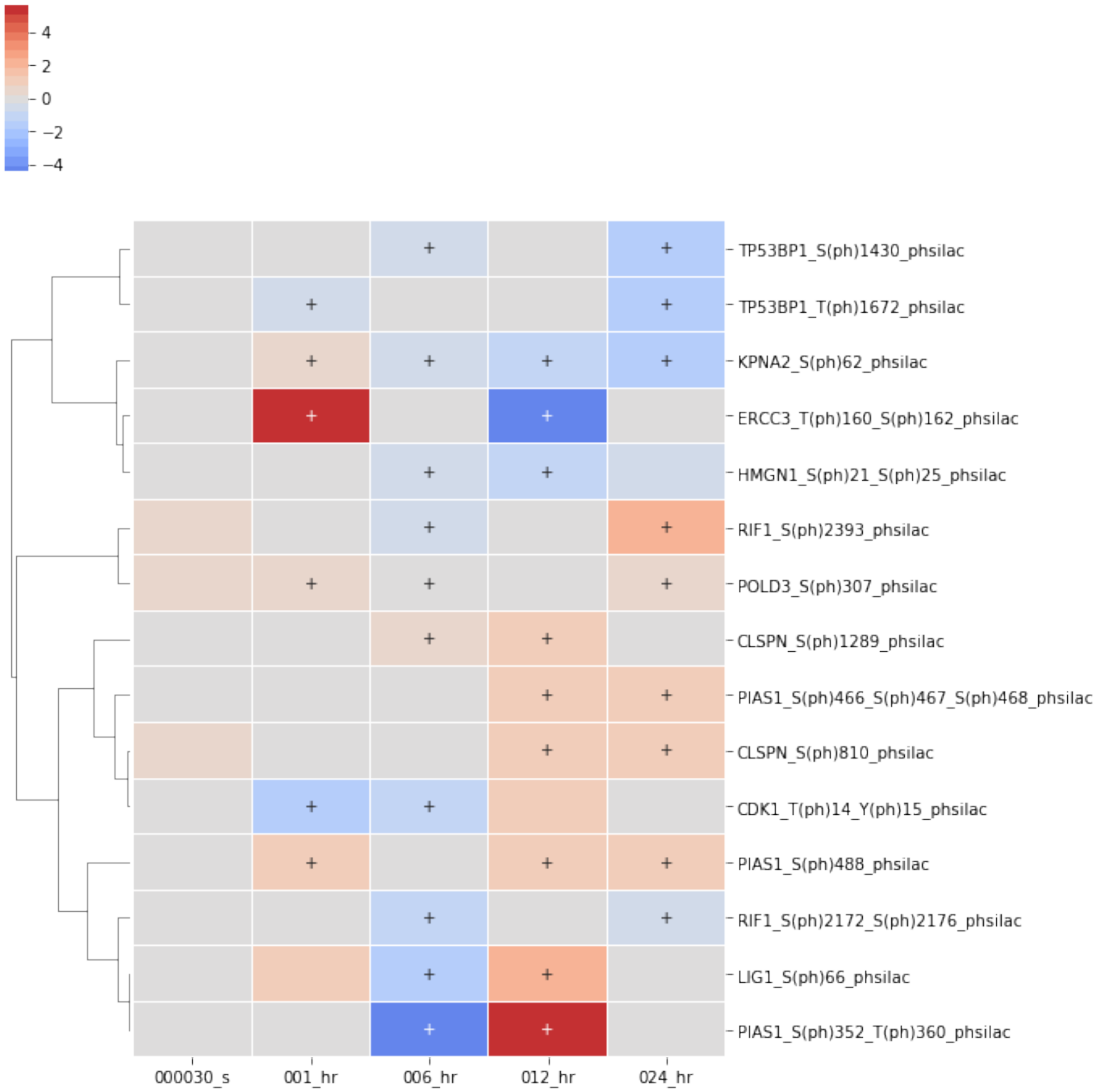
4
2
0
−2
−4
TP53BP1_S(ph)1430_phsilac
TP53BP1_T(ph)1672_phsilac
KPNA2_S(ph)62_phsilac
ERCC3_T(ph)160_S(ph)162_phsilac
HMGN1_S(ph)21_S(ph)25_phsilac
RIF1_S(ph)2393_phsilac
POLD3_S(ph)307_phsilac
CLSPN_S(ph)1289_phsilac
PIAS1_S(ph)466_S(ph)467_S(ph)468_phsilac
CLSPN_S(ph)810_phsilac
CDK1_T(ph)14_Y(ph)15_phsilac
PIAS1_S(ph)488_phsilac
RIF1_S(ph)2172_S(ph)2176_phsilac
LIG1_S(ph)66_phsilac
PIAS1_S(ph)352_T(ph)360_phsilac
000030_s
001_hr
006_hr
012_hr
024_hr

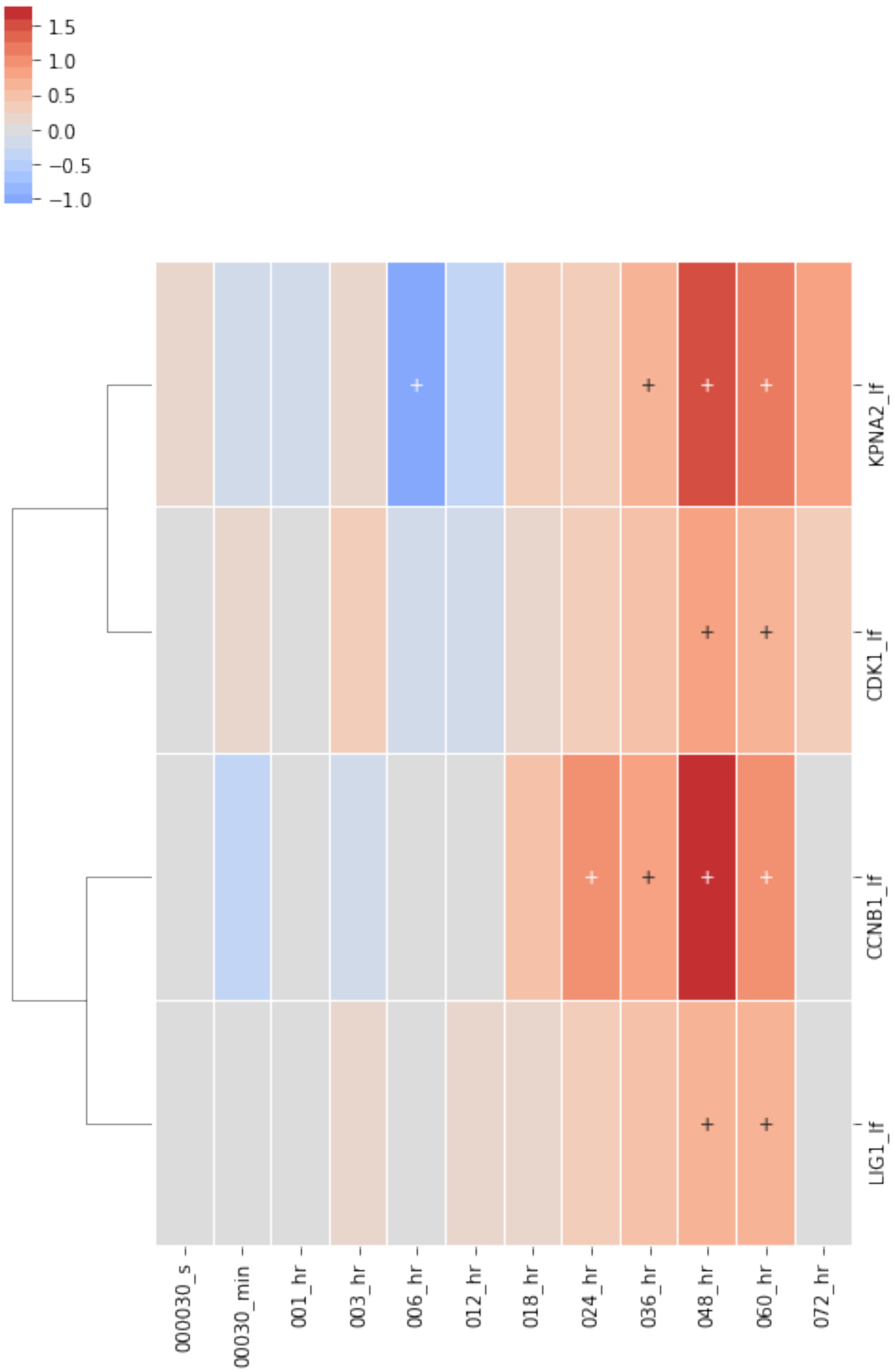
1.5
1.0
0.5
0.0
−0.5
−1.0
KPNA2_lf
CDK1_lf
CCNB1_lf
LIG1_lf
000030_s
00030_min
001_hr
003_hr
006_hr
012_hr
018_hr
024_hr
036_hr
048_hr
060_hr
072_hr

[51]:
```
down_casp3 = net_sub.expand_neighbors(
    network=None,
    nodes=['CASP3'],
    upstream=False, downstream=True,
    max_dist=1,
    include_only=exp_data.label_free.sig.require_n_sig(index='label', n_sig=2).
↪id_list
)
down_casp3 = utils.delete_disconnected_network(down_casp3)
print(len(down_casp3.nodes))
print(len(down_casp3.edges))
exp_data.label_free.heatmap(
    down_casp3.nodes,
    subset_index='identifier', index='label',
    min_sig=2, linewidths=0.01,
    cluster_row=True
);

exp_data.label_free.require_n_sig(index='label', n_sig=2).heatmap(
    down_casp3.nodes, subset_index='identifier',
    #index='label',
    rank_index=True,
    figsize=(8, 2),
    index='sample_id', columns='label',
    min_sig=2, linewidths=0.01, cluster_row=False, cluster_col=False
);
plt.yticks(rotation=0)
plt.savefig('down_from_casp3.png', dpi=300, bbox_inches='tight')
```

```
10
9
```

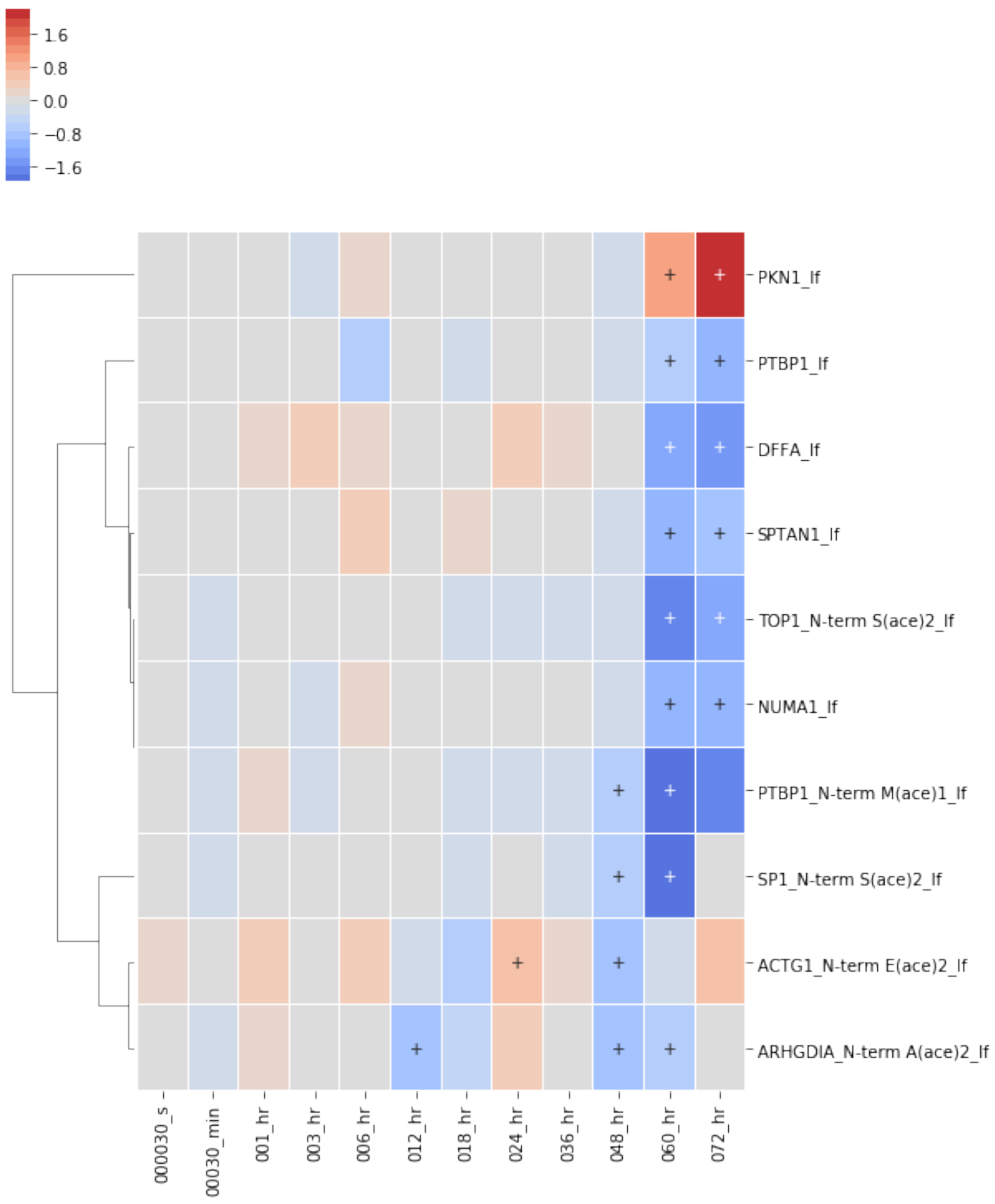
1.6
0.8
0.0
−0.8
−1.6
PKN1_lf
PTBP1_lf
DFFA_lf
SPTAN1_lf
TOP1_N-term S(ace)2_lf
NUMA1_lf
PTBP1_N-term M(ace)1_lf
SP1_N-term S(ace)2_lf
ACTG1_N-term E(ace)2_lf
ARHGDIA_N-term A(ace)2_lf
000030_s
00030_min
001_hr
003_hr
006_hr
012_hr
018_hr
024_hr
036_hr
048_hr
060_hr
072_hr

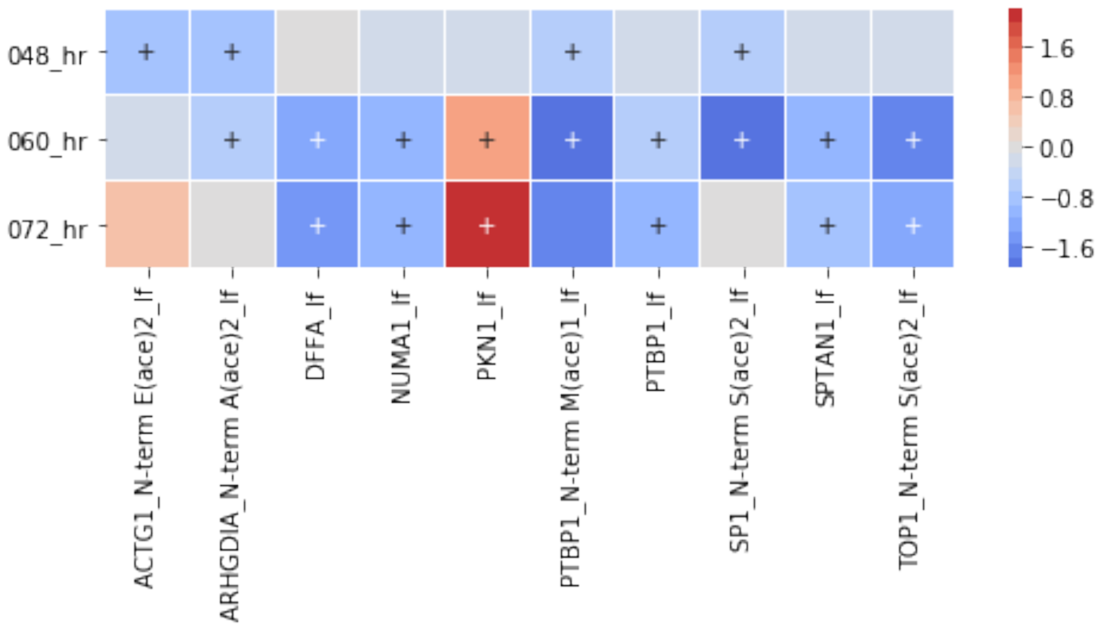


[52]:
```
vis.draw_cyjs(down_casp3)
```

```
<IPython.core.display.HTML object>
```

[53]:
```
def show_neighbors(node, df, upstream=True, downstream=False, max_dist=1,
                   include_only=None, figsize=None):

    df_copy = df.copy()
    df_copy.require_n_sig(n_sig=1, inplace=True)

    neighbors = net_sub.expand_neighbors(
        network=None,
        nodes=[node],
        upstream=upstream,
        downstream=downstream,
        max_dist=max_dist,
        include_only=include_only
    )

    neighbors = utils.delete_disconnected_network(neighbors)
    s_name =  'node_{}.png'.format(node)
    exporters.export_to_dot(neighbors, s_name, image_format='png',␣
↪engine='circo')
    display(Image(s_name, width=400))

    g = df_copy.heatmap(
```

```
        sorted(neighbors.nodes),
        subset_index='identifier',
        index='label',
        min_sig=1,
        rank_index=True,
        linewidths=0.01,
        figsize=figsize
    );

    g.savefig('{}_heatmap.png'.format(node), bbox_inches='tight', dpi=300)
show_neighbors('CASP3', exp_data.label_free, False, True, max_dist=1,
              include_only=exp_data.label_free.sig.require_n_sig(n_sig=2).
 ↪id_list)
```

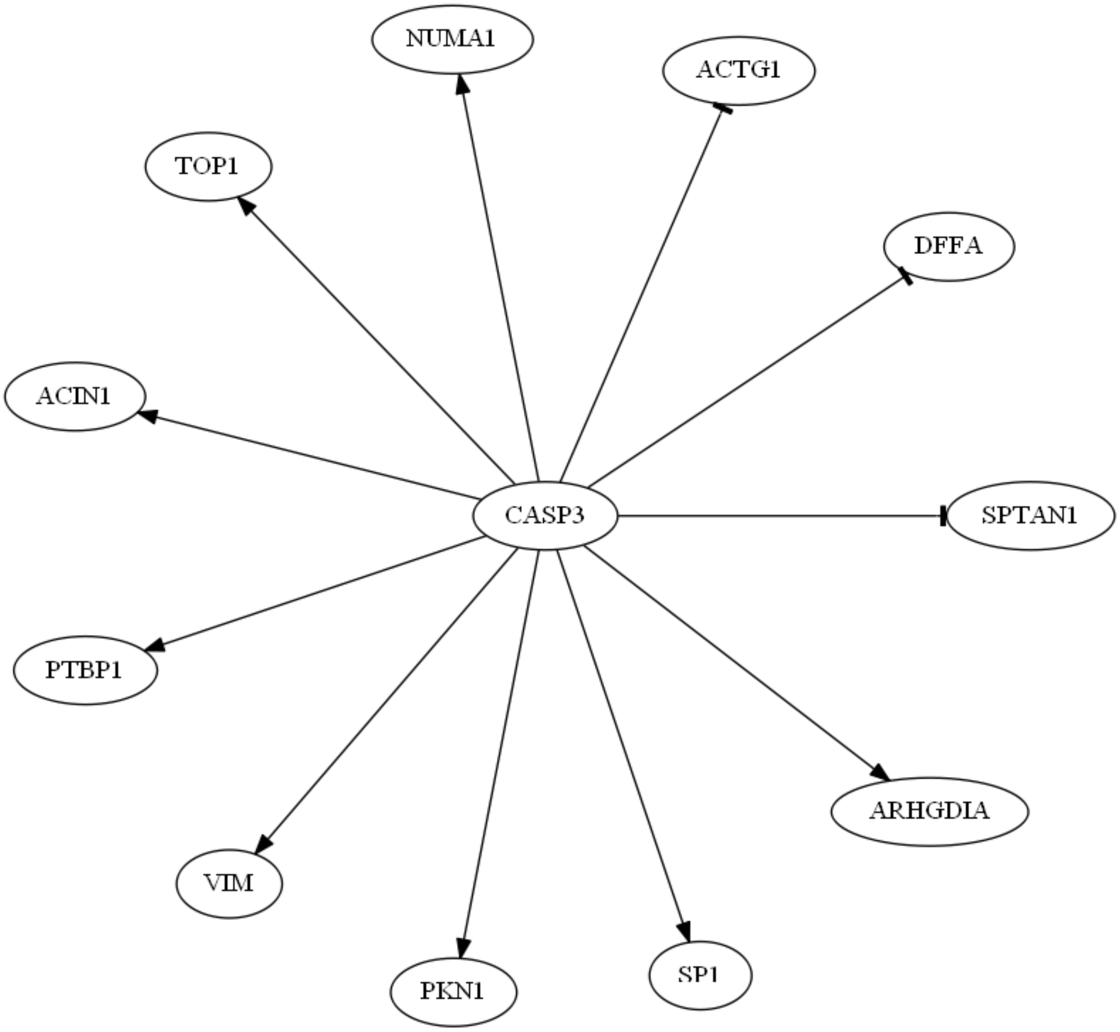

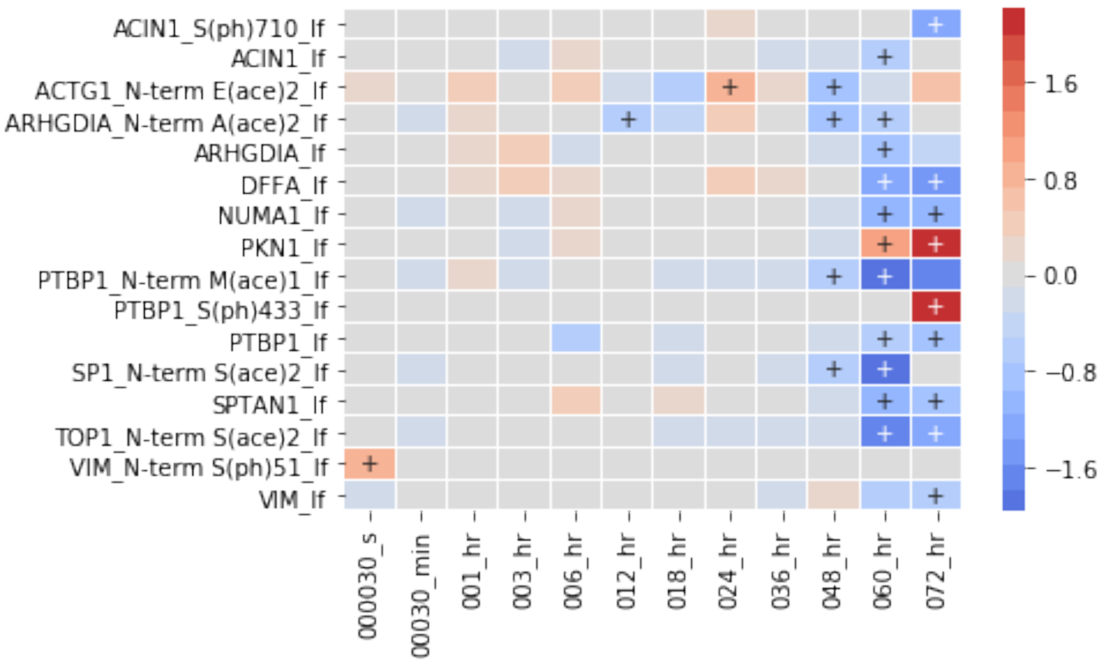


[54]:
```
show_neighbors('BAX', exp_data.species, False, True,  max_dist=1,
                include_only=exp_data.species.sig.require_n_sig(n_sig=1).id_list)
```

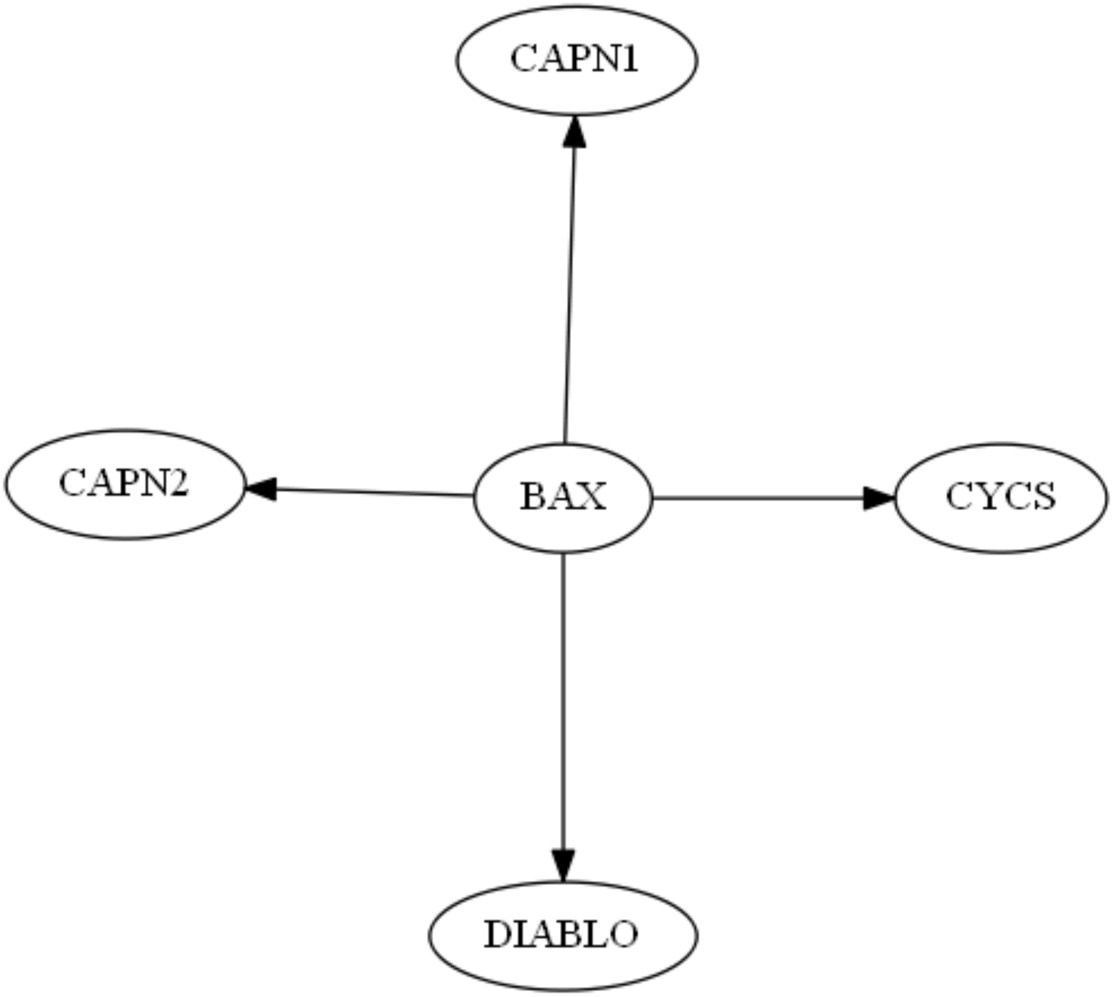
CAPN1
CAPN2
BAX
CYCS
DIABLO

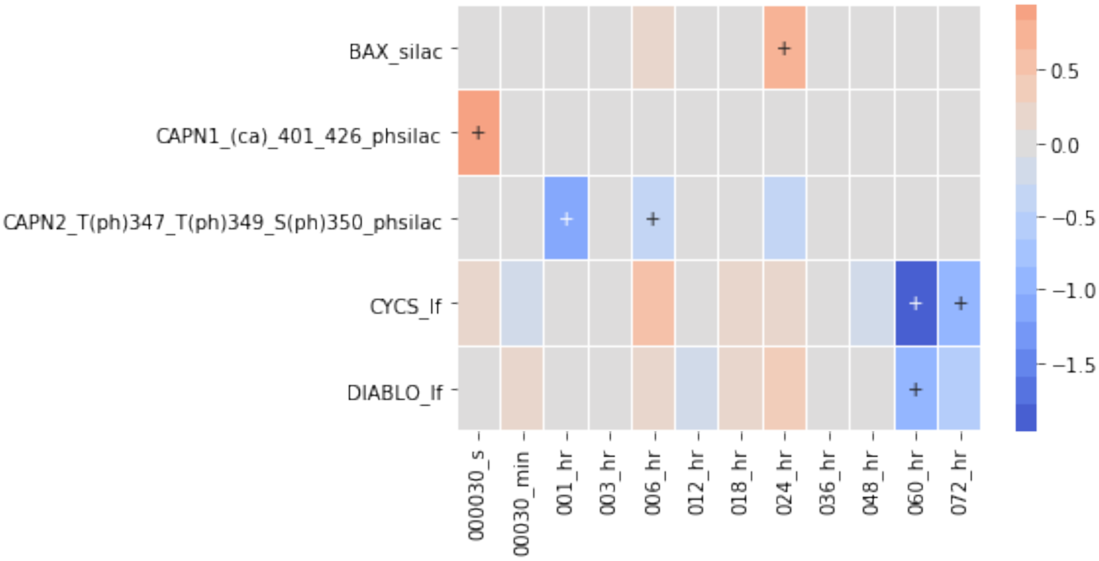
BAX_silac
CAPN1_(ca)_401_426_phsilac
CAPN2_T(ph)347_T(ph)349_S(ph)350_phsilac
CYCS_lf
DIABLO_lf
000030_s
00030_min
001_hr
003_hr
006_hr
012_hr
018_hr
024_hr
036_hr
048_hr
060_hr
072_hr
0.5
0.0
−0.5
−1.0
−1.5
