## Supplementary figures and images for "A computational framework to explore cellular response mechanisms from multi-omics datasets"

### Animated Annotated Gene-Set Network Dynamics

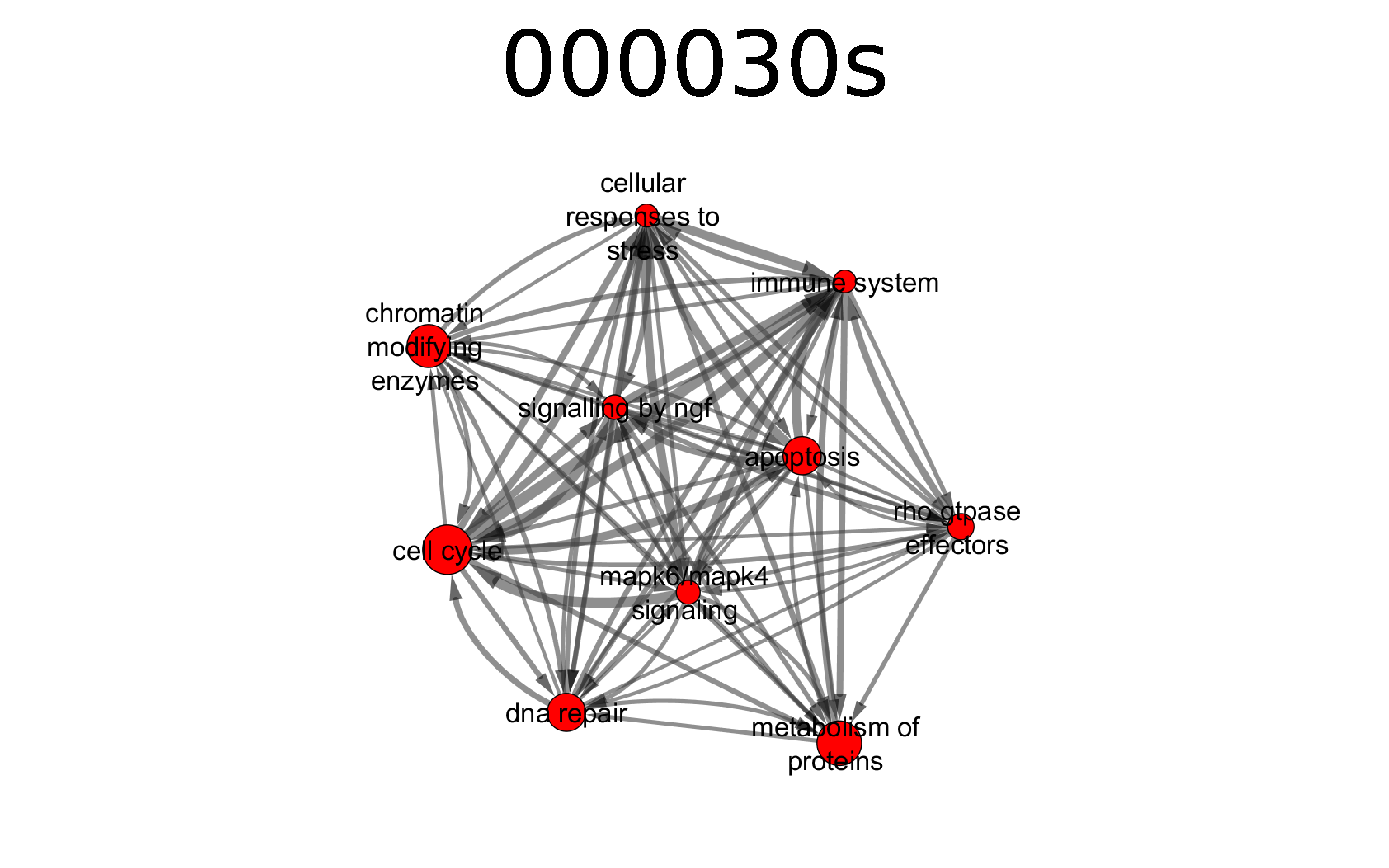
